# Reconstruction of microbial haplotypes by integration of statistical and physical linkage in scaffolding

**DOI:** 10.1101/2020.03.29.014704

**Authors:** Chen Cao, Jingni He, Lauren Mak, Deshan Perera, Devin Kwok, Jia Wang, Minghao Li, Tobias Mourier, Stefan Gavriliuc, Matthew Greenberg, A. Sorana Morrissy, Laura K. Sycuro, Guang Yang, Daniel C. Jeffares, Quan Long

## Abstract

DNA sequencing technologies provide unprecedented opportunities to analyze within-host evolution of microorganism populations. Often, within-host populations are analyzed via pooled sequencing of the population, which contains multiple individuals or ‘haplotypes’. However, current next-generation sequencing instruments, in conjunction with single-molecule barcoded linked-reads, cannot distinguish long haplotypes directly. Computational reconstruction of haplotypes from pooled sequencing has been attempted in virology, bacterial genomics, metagenomics and human genetics, using algorithms based on either cross-host genetic sharing or within-host genomic reads. Here we describe PoolHapX, a flexible computational approach that integrates information from both genetic sharing and genomic sequencing. We demonstrated that PoolHapX outperforms state-of-the-art tools tailored to specific organismal systems, and is robust to within-host evolution. Importantly, together with barcoded linked-reads, PoolHapX can infer whole-chromosome-scale haplotypes from 50 pools each containing 12 different haplotypes. By analyzing real data, we uncovered dynamic variations in the evolutionary processes of within-patient HIV populations previously unobserved in single position-based analysis.

## INTRODUCTION

Microorganisms are in a constant state of genetic flux in response to their environments. High-resolution analyses of these systems may lead to translational applications, such as clinical monitoring of antimicrobial resistance trends (Hofer 2019). However, analyzing within-host dynamics and evolution is challenging due to the difficulty of separating samples into genetically homogeneous isolates/clones, either by experimental procedures such as culturing individual strains, or current computational tools, which are unable to distinguish between many clones when haplotypes are unknown. As a pragmatic alternative, uncultured mixtures of the heterogeneous population are sequenced and analyzed based on aggregated frequencies at each segregating sites in individual hosts. This procedure ignores the fact that long-range linkage information is crucial in to many different analyses towards evolution (Sabeti, et al. 2002; Voight, et al. 2006) and association mapping (Datta and Biswas 2016). Recent advances in DNA sequencing technology led to increases in the sequencing depth per run and the length of reads, allowing us to assess genetic variants in greater detail, providing an unprecedented opportunity to understand the dynamics and evolution of these systems. Emergent single-molecule sequencing approaches that barcode short reads derived from long DNA fragments up to 50 Kb (i.e. linked-reads) (Zheng, et al. 2016; Chen, et al. 2019; Wang, et al. 2019), allow in-depth evolutionary studies into complex populations that were unresolvable from short fragment libraries (250 bp).

However, even with barcode-based linking, full resolution of short reads into long-range haplotypes is not feasible with current approaches. While third-generation technologies such as Pacific Biosciences and Oxford Nanopore Technology (Weirather, et al. 2017), produce very long reads that represent local haplotypes, computational analysis will still be essential for resolving haplotypes and estimating their relative proportions in the pool. Without this haplotype-level resolution, within-host dynamics cannot be analyzed as if the haplotypes were separated and sequenced individually. Even with barcoded linked-reads designed for single-molecule sequencing, the current tools are only applicable in a two-haplotype system, as it was designed for analyzing paternal and maternal chromosome (Mostovoy, et al. 2016) and structural variants (Elyanow, et al. 2018). Therefore, microorganism-based studies resort to computational tools to make single-cell-like analyses possible (Danko, et al. 2019).

Many tools have been developed to reconstruct haplotypes using algorithms that target data from viruses, bacteria, metagenomic data, and historically, artificially pooled human genomes. Conceptually they can be split into two categories. The first contains statistical models utilizing haplotype-block sharing between individuals (“statistical linkage disequilibrium” or “statistical LD” hereafter), mostly developed in human genetics (Long, et al. 2011; Long 2017). The second contains computational algorithms that leverage minor allele frequency and sequence reads exposing the co-occurrence of multiple alleles on the same haplotype (referred to as “physical linkage disequilibrium” or “physical LD” hereafter), mostly tailored to uncultured sequencing of viruses or bacteria (Huang, et al. 2011; Prabhakaran, et al. 2013; Pulido-Tamayo, et al. 2015; Albanese and Donati 2017; Artyomenko, et al. 2017; Posada-Cespedes, et al. 2017; Ahn, et al. 2018; Knyazev, et al. 2018; Li, et al. 2019; Nicholls, et al. 2019; Ke and Vikalo 2020; Knyazev, et al. 2020). A more detailed overview of these methods is available in the **Supplementary Notes SECTION I**.

There is a disconnect between these two approaches, and both have limitations. Methods based on genetic sharing do not consider pool-specific dynamics that can be captured by sequencing reads, and are ineffective when there is strong within-host evolution that changes allele distributions. Also, the underlying population genetic models developed in human genetics may not fit in microorganisms well. For instance, many microorganisms exchange genetic materials through gene conversions (Santoyo and Romero 2005), instead of meiotic recombination as assumed by many phasing tools (Browning and Browning 2011). On the other hand, methods based on genomic reads only work on one pool a time, without taking advantage of genetic sharing caused by transmission between hosts (Cudini, et al. 2019) and common developmental trajectories in different hosts (Toprak, et al. 2011). Additionally, organisms with differing properties such as mutation rates generate data that require field-specific assumptions to process. The result is that each haplotyping tool may be ineffective when applied to another data type.

The two current methodologies utilize disparate sources of genetic information. We expected considerable improvements by integrating them into a reconstructive model that accurately represents the composition of genetically related populations. In this work, we present a novel tool, PoolHapX, which utilizes a multi-step framework that balances cross-host shared information and host-specific information to jointly and simultaneously reconstruct haplotypes for multiple samples.

The remainder of this paper is organized as follows. PoolHapX’s design philosophy, an overview of its algorithmic framework, and examples of large-scale microorganism studies that PoolHapX facilitates are outlined in New Approaches. Thorough comparisons to other tools with both simulated and real data validating our design philosophy are presented in Results. As a practical demonstration of experimental applicability, PoolHapX is applied to a time-series of HIV samples to elucidate within-host evolutionary dynamics. More extensive descriptions of software and simulation design, including implementation details and analysis procedures, can be found in the Materials & Methods.

## NEW APPROACHES

### Applications empowered by PoolHapX

#### Assumptions

PoolHapX is designed for studies where researchers have sequenced microbial genomes in multiple samples and there is known to be genetic sharing across the samples. For example, an archetypal dataset may be several samples from patients known to be infected by the same pathogen. In that case, transmission facilitates pathogen genetic sharing. Another dataset could consist of multiple samples from the same individual at multiple timepoints during tissue development or disease progression. In this case, genetic sharing is caused by the developmental trajectory. We assume that the investigators are interested in within-host evolution at the haplotype scale, as opposed to single-nucleotide polymorphism (SNP) studies. Unlike previous tools that assume the availability of the identities of haplotypes and only estimate frequencies (Long, et al. 2011; Albanese and Donati 2017), PoolHapX can infer haplotype identities (out of the potential *2*^*n*^ candidates, where *n* is the number of segregating sites). That is why we claim a *de novo* haplotype reconstruction. At the moment, PoolHapX does not aim to assemble reference genomes *de novo* such as SPAdes (Bankevich, et al. 2012).

#### Applicable organism and sequencing protocols

PoolHapX is applicable to any microorganism with a reference genome, including viruses, bacteria and protozoans. PoolHapX is also applicable to the analysis of within-species haplotypes from metagenomics data, after the sequencing reads of the focal species are mapped to the corresponding references. PoolHapX is applicable to NGS data, including standard short reads as well as barcoded linked-reads, which was popularized by 10x-Genomics (Zheng, et al. 2016; Wang, et al. 2019), and is supported by several other vendors after 10x-Genomics’ suspension of the service (Chen, et al. 2019; Wang, et al. 2019).

### Design Philosophy of PoolHapX

PoolHapX integrates cross-host shared information (i.e., statistical LD estimated from population genetic models) and host-specific information (i.e., physical LD of alleles on the same genomic reads) in a flexible framework. Here, “pool” and “host” are both used synonymously, since “host” emphasizes the biological source of data and “pool” the experimental source. Briefly, statistical LD in a population is equivalent to non-zero correlation between the occurrences of alleles at different locations, regardless of the biological mechanism (e.g., recombination, gene conversion, or natural selection). We use hierarchical multivariate normal distributions to model and resolve the statistical LD across sub-genomic regions to generate intermediary haplotype fragments. Physical LD can be observed in paired-end reads or linked-reads, constraining the combinations of haplotypes that can occur in specific pools. We utilize this information in two ways: (1) to generate initial draft haplotypes and (2) to statistically infer final haplotypes using a regularized regression model. Below, we describe the 4-step algorithm and, intuitively, the innovation and benefit of each step, leaving more details in Materials and Methods as well as **Supplementary Notes SECTION II**.

### Overview of the PoolHapX algorithm

PoolHapX uses variant calls and read alignments to identify global haplotypes in each host, and estimate their within-host frequencies (**Supplementary Notes SECTION III Fig. 8**). The PoolHapX algorithm is comprised of four steps embedded in a divide-and-conquer framework. In Step 1, graph-coloring (Matula, et al. 1972) is employed to roughly cluster sequencing reads into initial draft haplotypes. This draft set serves as the first step of the divide-and-conquer process (**Fig. 1a, Supplementary Notes SECTION II Fig. 1**). In Step 2, our hierarchical Approximate Expectation-Maximization (AEM) algorithm is applied to infer haplotypes in local regions by incorporating information from multiple hosts. The algorithm starts with the smallest local haplotypes as the lowest hierarchy (**Fig. 1b**), and then gradually combines them into successively longer local haplotypes covering larger spans of variant positions (**Fig. 1b, Supplementary Notes SECTION II Fig. 3**). This process iterates through several rounds until reaching the top level (representing the largest local region that AEM can analyze with the available computation memory). In Step 3, the refined set of local haplotypes from the final iteration of AEM will be stitched to form candidate global haplotypes using a Breadth-First Search (BFS) algorithm (**Fig. 1c, Supplementary Notes SECTION II Fig. 6**). Finally, in Step 4, the long-range linkage implied in allele frequencies aggregated across all hosts and the short-range linkage sequencing reads from each pool are innovatively integrated in a regression model (**Fig. 1d, Supplementary Notes SECTION II Fig. 7**). This regression model is solved using a regularized objective function (Hazimeh and Mazumder 2018). The input candidate global haplotypes are from all the pools (stitched in Step 3), and the regression is conducted in individual pools sequentially. Then aggregating in-pool frequencies will lead to the cross-pool global frequencies. This step finalizes the identity and frequency of the global haplotypes.

**Figure 1.**
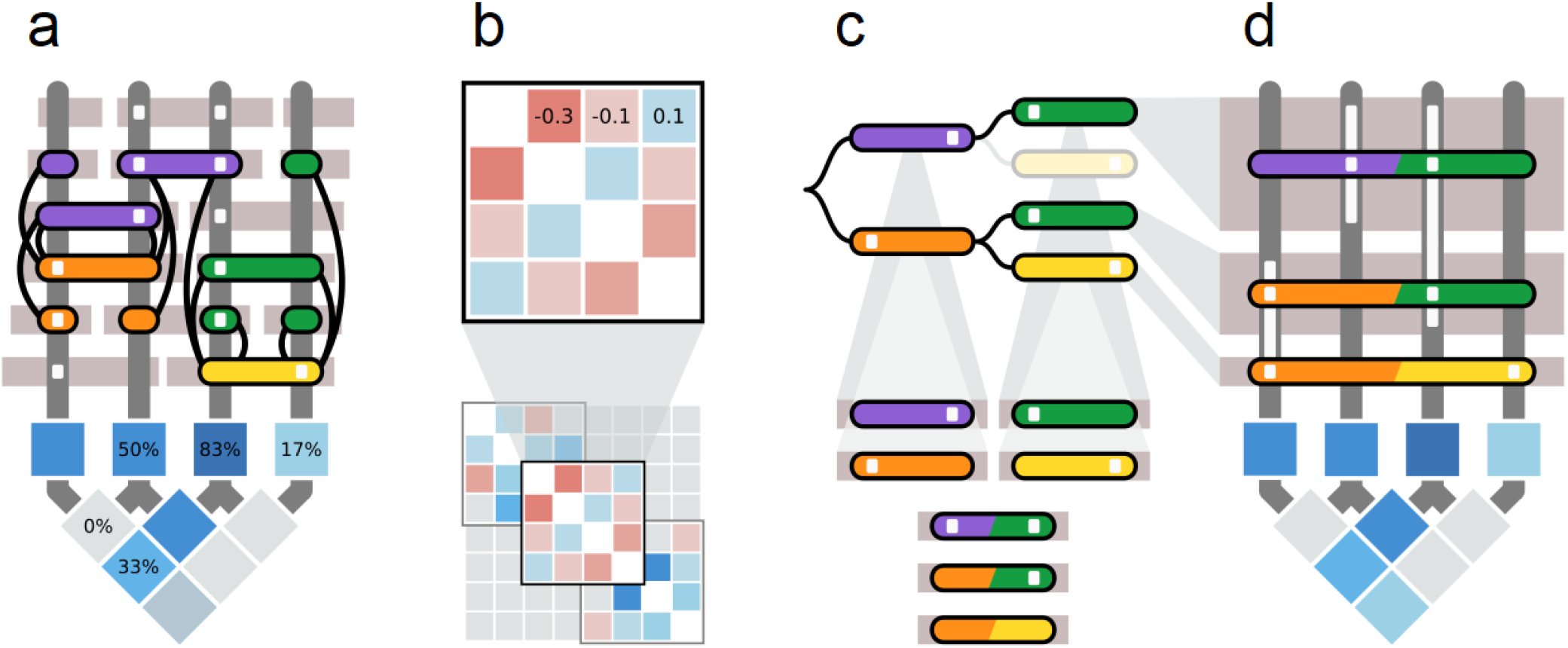
The PoolHapX algorithm. An example of the PoolHapX algorithm applied to a dataset containing reads from 3 haplotypes 0110, 1010, and 1001 with proportions 1/2, 1/3, and 1/6 respectively. Input consists of sequence reads (horizontal grey rectangles above) and allele frequencies of individual (square) and paired (diamond) sites (below). Vertical (dark grey) bars denote the locations of polymorphic sites, and white squares indicate the presence of alternate alleles. Colored rectangles represent haplotype information inferred by PoolHapX. (a) A graph is formed where nodes (coloured rectangles) are unique reads, and edges (black lines) are drawn between nodes with differing alleles at any polymorphic site. Graph coloring is applied to differentiate nodes with conflicting alleles (joined by edges), forming initial haplotypes for the next step in the algorithm (AEM). (b) Hierarchical Approximate Expectation Maximization (AEM) is applied to the initial covariance information derived from graph coloring. AEM is applied to a variance-covariance matrix representing overlapping local regions (containing 4 variants in this example). The result is then extended to a larger region (8 variants) by tiling the local regions (bottom). Here, blue squares indicate positive statistical LD between pairs of variants, and red indicates negative LD. (c) Haplotype segments from the final iteration of AEM are combined into a tree (top), whose branches represent when two adjacent segments have identical alleles in their overlapping regions (multicolored rectangles at bottom). Breadth-first tree search (BFS) is used to exhaustively search all possible branches to form global haplotypes. (d) A regularized (L0+L1) regression is used to estimate haplotype frequencies (height of rectangles above) from the individual (square) and paired (diamond) allele frequency information (below). More detailed illustrations of the algorithms are in **Supplementary Figures 1-7**.

### The innovation and benefit of each step

The above four steps incorporate physical and statistical linkage into a coherent model. Since the techniques were pioneered in multiple organismal fields to address their specific challenges, the integrated PoolHapX model might be difficult to understand as a whole. We have summarized the innovations and benefits of jointly using these techniques together below.

The innovation of Step 1 lies in its adoption of a greedy strategy to form many haplotypes (including potential false positives) for the downstream analysis, instead of the parsimony principle utilized by most tools based on graph algorithms in viral genomics (Zagordi, et al. 2011; Prosperi and Salemi 2012; Topfer, et al. 2014; Baaijens, et al. 2017). Due to its multi-step framework, PoolHapX relies on downstream steps to remove implausible haplotypes. The innovation of Step 2 is the hierarchical iteration of AEM, an established technique in human genetics (Kuk, et al. 2009). The original AEM calculated the likelihoods for all 2^n^ haplotypes (Kuk, et al. 2009), and is therefore is not scalable for genome-scale analysis. Step 3 is a standard divide-and-conquer algorithm, which links sub-genomic fragments from hierarchical iterative AEM (Step 2) into full-length candidate haplotypes for Step 4. The innovation of Step 4 lies in both its design and implementation. Traditional methods based on regressions in haplotype reconstructions for viruses (Leviyang, et al. 2017), plants (Long, et al. 2011), and metagenomics (Albanese and Donati 2017) only utilize allele frequencies at individual segregating sites. In contrast, the PoolHapX regularization model integrates both allele frequencies and physical LD between alleles co-occurring on sequencing reads. To constrain the number of reconstructed haplotypes, traditional methods solve regressions using L1 regularization, which do not distinguish between the correct solution with few haplotypes, and incorrect solutions with many haplotypes. L1-based regression models assign the same penalty to any reconstructed population as long as the sum of the allele frequencies is 1.0. To parsimoniously narrow down the set of haplotypes comprising a sample, PoolHapX uses a combination of L0 and L1 regression penalties based on a cutting-edge L0 solver (Hazimeh and Mazumder 2018).

In summary, Steps 1 and 4 utilize host-specific information (physical linkage), whereas Step 2 iteratively leverages cross-host linkage-sharing information. Step 3 stitches the sub-genomic haplotype fragments into full-length haplotype candidates. By building on existing innovations drawn from multiple fields, PoolHapX aggregates the benefits of multiple statistical models, and is thus applicable to many datasets.

## RESULTS

Using simulated and real data containing tens of haplotypes in multiple pools, we demonstrate that PoolHapX possesses the desired properties for being a universally applicable tool robust to various factors: (1) it is applicable to many fields, and in particular outperforms state-of-the-art haplotype-reconstruction tools targeting different domains, i.e., virus, bacteria, and metagenomics in their respective settings; (2) it is robust to various scenarios of within-host evolution; (3) it can reconstruct many whole-chromosome long-range haplotypes when applied to barcoded linked-reads and (4) haplotypes inferred by PoolHapX reveal novel evolutionary insights unseen in SNP-based analyses.

### Benchmarking PoolHapX with simulated data

To test the accuracy and flexibility of PoolHapX in comparison to state-of-the-art haplotyping tools (Kuk, et al. 2009; Pirinen 2009; Prabhakaran, et al. 2013; Pulido-Tamayo, et al. 2015; Albanese and Donati 2017; Ahn, et al. 2018; Knyazev, et al. 2018; Li, et al. 2019; Nicholls, et al. 2019), we simulated artificial pools of haplotypes and the sequencing reads generated from these pools. We then examined PoolHapX against tools developed for virology (Prabhakaran, et al. 2013; Ahn, et al. 2018; Knyazev, et al. 2018), bacteriology (Pulido-Tamayo, et al. 2015; Li, et al. 2019), metagenomics (Albanese and Donati 2017; Nicholls, et al. 2019), and human genetics (Kuk, et al. 2009; Pirinen 2009) (**flowchart** in **Supplementary SECTION III Fig. 8**). As each discipline has a specific method for simulating benchmarking data, we follow their corresponding conventions. For standardized assessment criteria to consistently compare reconstruction accuracy, we used two sets of metrics: (1) MCC (Matthews Correlation Coefficient) +JSD (Jensen–Shannon Divergence) that were used frequently in metagenomics field (Luo, et al. 2015; Albanese and Donati 2017; Antwis, et al. 2018) and (2) F1-Score + CRR (Correctly Reconstructed Rate) + FDR (False Discovery Rate) (**Supplementary Notes SECTION IV**) that are used in viral genomics field (Prabhakaran, et al. 2013). In the first set of metrics, we used MCC to measure the similarity between the identities of simulated ‘gold standard’ and reconstructed haplotypes, where two identical haplotypes have their MCC = 1 (the larger, the better) (Albanese and Donati 2017), we calculated the MCC of each gold-standard haplotype and the closest reconstructed haplotype (by Hamming distance), and averaged the haplotype-MCCs across all of the samples. The gold-standard haplotype and reconstructed haplotype were considered as vectors composed of 0s and 1s, and pairs of matched and mismatched alleles were counted as true-positives, false-negatives, etc. (Luo, et al. 2015; Albanese and Donati 2017; Ahn, et al. 2018). We used JSD to measure the difference between frequencies of simulated and reconstructed haplotype distributions for each host in the simulated dataset, where two identical distributions have JSD = 0 (the smaller, the better). In the second set of metrics, we first define the True Positive as the haplotypes that are correctly identified, and then define F1, CRR and FDR using standard conventions (**Supplementary Notes SECTION IV**). **Fig. 2** shows the results of this comparison using MCC/JSD, and **Supplementary SECTION VII Figs. 15 & 16** presents the same comparison in terms of F1-Score/CRR/FDR. A brief description of each domain is provided below. Details of the simulations are presented in the **Supplementary Notes SECTION III** and Online Methods, and outcomes with more parameters, showing similar trends, are presented in **Supplementary SECTION VII Figs. 12-14**).

**Figure 2.**
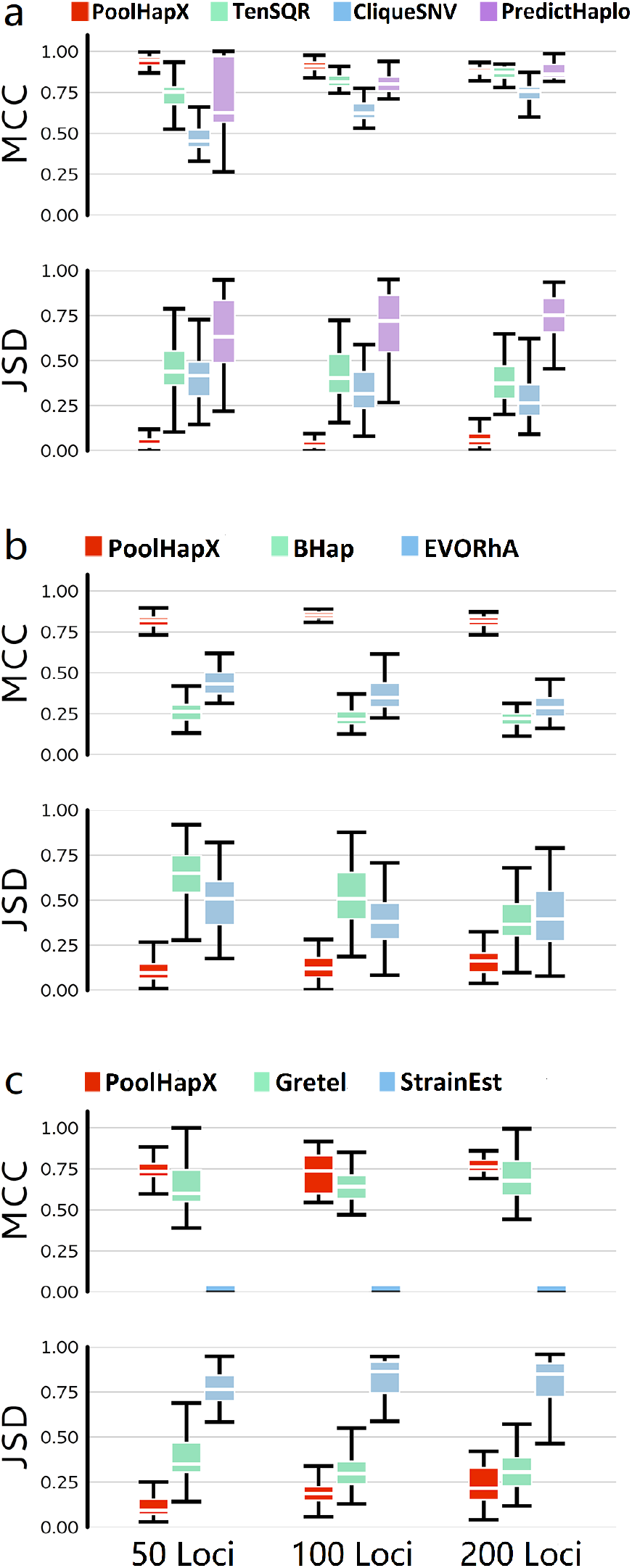
Comparison between PoolHapX and existing tools. For all panels, the upper half shows the accuracy for haplotype identity (MCC) and the lower half shows the accuracy for haplotype frequency (JSD). The x-axis denotes number of segregating sites in the haplotype. Boxes extend to the first and third quartile; whiskers extend to the upper and lower value. (a-c) Number of pools = 50. (a) tools to reconstruct viral haplotypes, TenSQR, PredictHaplo and CliqueSNV. Sequencing coverage per pool = 5000X. Number of haplotypes for 50 loci, 100 loci and 200 loci are 41, 73 and 42, respectively. (b) tools to reconstruct bacterial haplotypes, Bhap and EVORhA. Coverage = 250X. Number of haplotypes for 50 loci, 100 loci and 200 loci are 42, 43 and 43, respectively. (c) tools to reconstruct haplotypes for metagenomics, Gretel and StrainEst. Coverage = 25X. Number of haplotypes for 50 loci, 100 loci and 200 loci are 39, 37 and 36, respectively.

#### Viruses & Bacteria

In the field of viral/bacterial haplotype reconstruction, cross-host linkage sharing was not used as a source of information, despite literature evidence demonstrating extensive conservation in some genomic regions even after transmission takes place (Mak, et al. 2020). As a result, when conducting comparisons, the authors of other tools usually formed only one pool of gold standard haplotypes in each round of simulation. A reference genome is used as a template, and variants are randomly simulated with pre-specified density of SNPs to form the gold standard haplotypes. Multiple densities are used to benchmark performance with diverse configurations of data, reflecting variable mutation rates in different environments (Metzgar and Wills 2000). In general, high SNP density (between 0.5% and 2.0%) for viruses (Prabhakaran, et al. 2013), and a lower range for bacteria (between 0.005% and 0.02%) (Li, et al. 2019) are used. Our procedure is similar, except that we simulated multiple pools with some haplotype sharing between them, and reconstructed haplotypes across the pools simultaneously.

It should be noted that our approach explicitly models haplotype sharing between hosts, which PoolHapX is naturally designed for. This led to relatively better performance than other tools designed for single pools of data. To accurately simulate linkage sharing between pools, we used SLiM (Haller and Messer 2019) to simulate haplotypes under standard island models, where genomes mutate, recombine, and replicate in their own island and occasionally migrate to other islands. We embedded the simulated variants into a viral reference genome.

For viruses, we chose the human immunodeficiency virus (HIV), which is well known for its ability to form large and genetically heterogeneous within-host viral populations (Lauring and Andino 2010). Multiple haplotypes were pooled and the simulated sequencing reads were processed using a modified version of the GATK best practice pipeline (DePristo, et al. 2011) to discover variants. Details can be found in **Supplementary Notes SECTION IV**. We chose three representative viral sequencing tools for comparison: TenSQR (Ahn, et al. 2018), PredictHaplo (Prabhakaran, et al. 2013) and CliqueSNV (Knyazev, et al. 2018). The details of how we run these tools are presented in **Supplementary Notes SECTION V**. Evidently, PoolHapX outperformed these alternatives not only in the mean of the MCC and JSD values, but also their variances (**Fig. 2a** and **Supplementary SECTION VII Fig. 12**). When sequencing coverage was high (=5000X), as well as the SNP density, PoolHapX performed similarly to other tools in terms of the mean MCC, but with much smaller variance of MCC and also significantly better JSD that represents the abundance of haplotypes (**Fig. 2a**). When sequencing coverage was low, the performance of other tools decreased rapidly relative to PoolHapX (**Supplementary SECTION VII Fig. 12 a-d**). The comparison using F1-Score/CRR/FDR shows the same trend between the viral tools under comparisons (**Supplementary SECTION VII Fig. 15 a-d, Supplementary SECTION Fig. 16 a-d**).

We are not able to directly use a commonly cited real dataset with a single pool containing only 5 HIV strains (Giallonardo, et al. 2014) because PoolHapX is designed to utilize genetic sharing among multiple pools. Instead, we generated a mocked data based on the 5 HIV strains (HIV-1_HXB2_, HIV-1_89:6_, HIV-1_JR-CSF_, HIV-1_NL4-3_ and HIV-1_YU2_) to mimic the multi-pool configuration. We randomly selected 5 breakpoints in the five templates and generated recombinants by simulating recombination using the breakpoints. Out of all recombinants, we randomly selected 25 haplotypes as the “gold standard” haplotypes. Then 25 pools are formed by mixing the gold-standard haplotypes and the in-pool abundance of each “gold standard” haplotype are randomly assigned. The outcome is presented in **Supplementary SECTION VII Fig. 17** showing that PoolHapX significantly outperforms alternatives in terms of both MCC/JSD metrics and F1/CRR/FDR metrics. In a concurrent study, we used this 5-strain dataset to compare our single-host tool, WgLink, that only uses single host sequencing to existing tools (Cao, et al. 2020). We showed that our strategy of divide-and-conquer and L0+L1-regularization with cross-host information is capable of outperforming alternative tools by a large margin (Cao, et al. 2020).

For bacteria, we used a chromosome of *Vibrio cholerae* O1 biovar *El Tor* str. N16961 (Chr-2, length = 1.07Mb), a strain of the bacterium *Vibrio cholerae* and the causative agent of cholera (Cvjetanovic and Barua 1972) as the template reference genome. We used a lower SNP density (=0.005% to 0.02%) to match the bacterial genomes (Pulido-Tamayo, et al. 2015) and to be comparable to the simulation procedures of competing tools, i.e., Bhap (Li, et al. 2019) and EVORhA (Pulido-Tamayo, et al. 2015). Despite the very different simulation parameters and reference genome in the simulations of viruses and bacteria, we observed similarly good performance in contrast to other tools using both MCC/JSD metrics (**Fig. 2b** and **Supplementary SECTION VII Fig. 12 e-h**) and Fi-Score/CRR/FDR metrics (**Supplementary SECTION VII Fig. 15 e-h, Supplementary SECTION VII Fig. 16 e-h**).

In the field of metagenomics there is no established tool to infer haplotypes *de novo* without utilizing template references of known strain sequences. Note that we still rely on reference genome for the species, instead of assembling genomes of novel species completely *de novo*. Gretel, a recently developed tool (Nicholls, et al. 2019), requires high SNP density so that each SNP is within a sequencing read length of the next adjacent SNP. We simulated data to facilitate the requirements of Gretel to ensure it is runnable (although this is not a requirement of PoolHapX). We also selected StrainEst (Albanese and Donati 2017), a representative tool for strain-identification, to demonstrate whether tools that utilize templates of known strains may work for fine-scale haplotype identification. This was likely an unfair comparison, due to the lack of template references for known strains in our simulations. We used sequences from *Escherichia coli*, a typical bacterium for meta-genomics studies. In these two cases, the sequencing coverage was substantially lower than dedicated single-species sequencing (=25X). Evidently, PoolHapX outperformed Gretel, and StrainEst did not work well when templates were unavailable in terms of both MCC/JSD (**Fig. 2c**; **Supplementary SECTION VII Fig. 12 i-l)** and F1-Score/CRR/FDR (**Supplementary SECTION VII Fig. 15 i-l, Supplementary SECTION VII Fig. 16 i-l**).

In order to figure out the contributions of different modules in PoolHapX, we also analyzed simulated bacteria data using partial algorithm of PoolHapX without the last step of regularization. The outcome is depicted in **Supplementary SECTION VII Fig. 18**, showing substantial improvement achieved by the final step that is critical in linking segmental haplotypes.

### PoolHapX performance in evolutionary scenarios and linked reads

The above simulations in four domains show that PoolHapX is universally applicable across species and is robust to sequencing data with varying properties. However, except in the domain of human GWAS, existing tools do not explicitly utilize sharing across hosts. Since hosts of microbes can exert different evolutionary pressures to generate host-specific haplotypes, this can cause a biased outcome from models that use cross-host sharing. To test whether the PoolHapX module utilizing within-host physical LD can correct for this bias, we used SLiM to simulate data under three common evolutionary scenarios in population genetic analyses: positive selection, negative selection, and selective sweeps (**Supplementary Notes SECTION III**). Our analysis of these data indicated that PoolHapX is robust to host-specific haplotype patterns caused by evolution, measured by both MCC/JSD (**Fig. 3**; **Supplementary Notes SECTION VII Fig. 13**) and F1-Score/CRR/FDR (**Supplementary Notes SECTION VII Fig. 19, Supplementary SECTION VII Fig. 20**).

**Figure 3.**
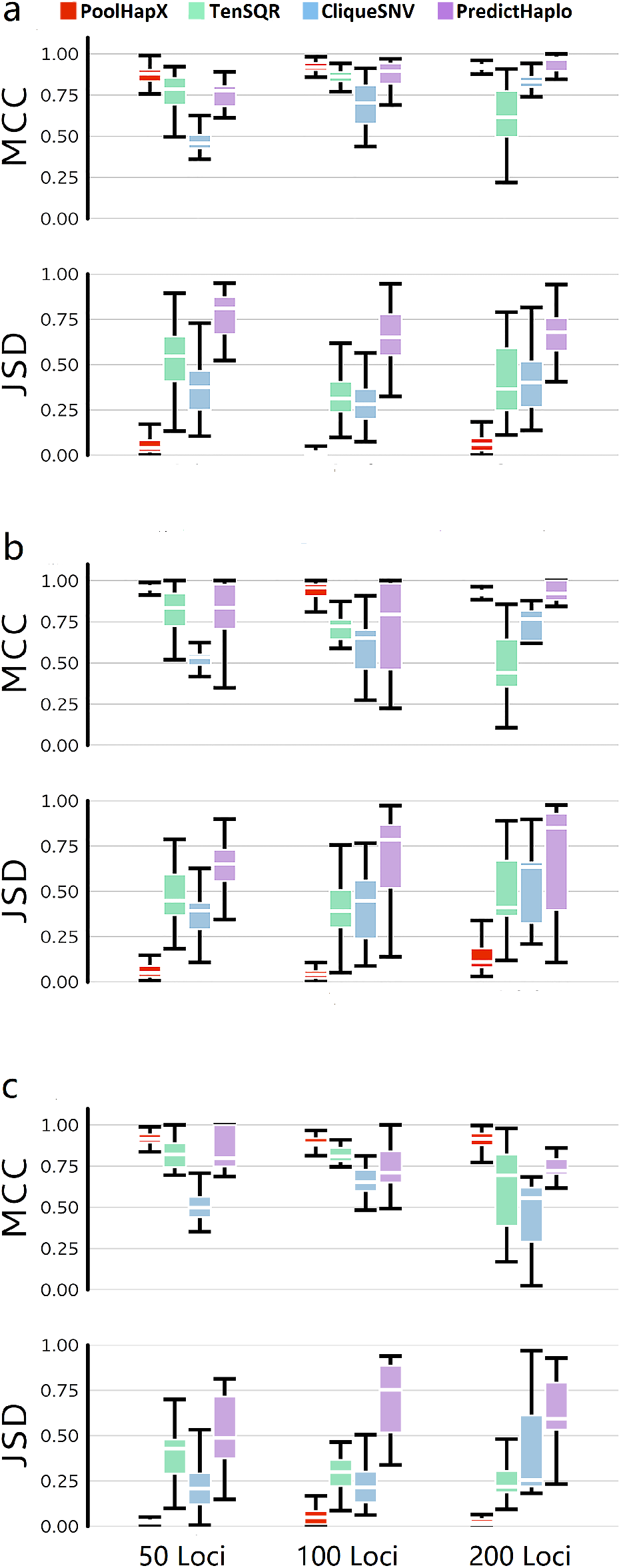
PoolHapX is robust to the within-host changes due to selective forces. The three evolutionary forces are: (a) Negative selection, number of haplotypes for 50 loci, 100 loci and 200 loci are 36, 36 and 45, respectively, (b) Positive selection, number of haplotypes for 50 loci, 100 loci and 200 loci are 41, 50 and 104, respectively, and (c) Selective Sweep, number of haplotypes for 50 loci, 100 loci and 200 loci are 23, 36 and 27, respectively. All three panels are comparing with viral tools, i.e., TenSQR, PredictHaplo and CliqueSNV. All data are simulated under coverage of 5000X, and 50 pools. The y-axis is the same as **Fig. 2**.

#### Single-molecule linked-reads

We further tested PoolHapX’s capabilities on single-*molecule* linked-reads. Based on a template of chromosome 1 of the unicellular green algae *Ostreococcus lucimarinus* (genome length of 1.15 Mb), we simulated approximately 20 gold standard haplotypes with 570 SNP positions. Using the 10X Genomics linked-read simulator LRSim (Luo, et al. 2017), with default settings of fragment length (=50Kb) in each droplet and number of linked-reads per fragment (on average 67), we simulated 10X Genomics linked-reads at various sequencing depths and numbers of pools. On average, PoolHapX achieved MCC ≥ 0.75 and JSD ≤ 0.25 (**Fig. 4**), F1-score around 0.75-1.0, CCR around 0.75-1.0 and FDR around 0.25 (**Supplementary Notes SECTION VII Fig. 21**). There results are comparable to other PoolHapX results when inferring shorter haplotypes using standard Illumina paired-end reads based on short-fragment DNA molecules. This outcome turns the promise of “single-cell” DNA sequencing into reality, enabling pathogen biologists to study within-host evolutionary changes at the individual molecule level.

**Figure 4.**
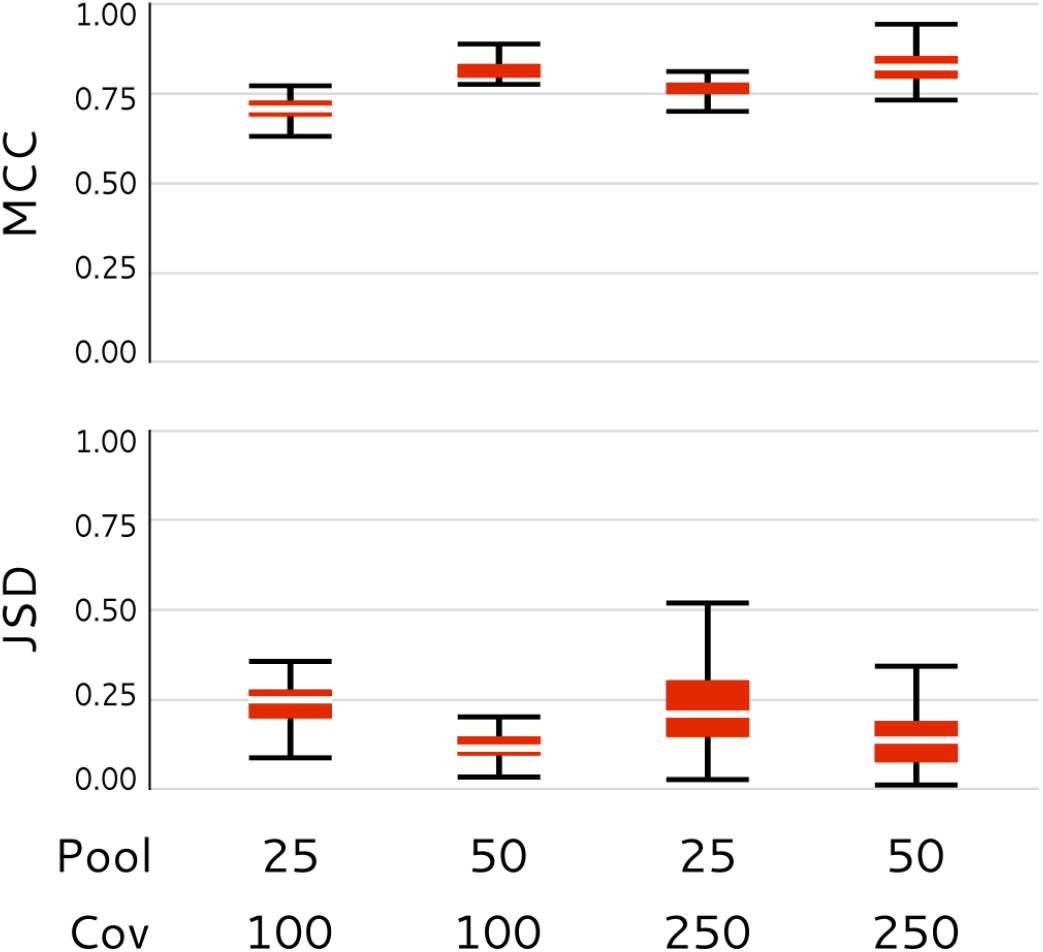
PoolHapX + single-cell linked-reads. MCC and JSD of PoolHapX applying to simulated 10x linked-reads using based on combinations of different number of pools (25,50) and sequencing coverage (100, 250).

### Applications to real data

Infections with the haploid malaria parasite *Plasmodium vixax* (Genome size =22 Mb) are known to contain multiple genotypes, which influence disease severity (Pacheco, et al. 2016). These ‘multi-clonal infections’ may derive from infection by a single mosquito bite carrying multiple strains, with meiotic recombination in the vector. Alternatively, they may be due to multiple infections from different mosquitos carrying different strains. Accurate whole-genome haplotype reconstruction will distinguish between these alternatives. We challenged PoolHapX with a collection of 49 *P. vivax* genome sequences (**Supplementary Notes SECTION VI**) to demonstrate its applicability on many pools of eukaryotic organisms with mid-sized genomes (Carlton 2003). To achieve this, we split the *P. vivax* genome into windows of 150 SNPs. PoolHapX took on average 54.79 CPU-hours per chromosome to conduct all computations. On average we found 3.3 inferred haplotypes per region per individual (**Supplementary Notes SECTION VI Table 3**), consistent to expectations of existence of multiplicity of infections (Pacheco, et al. 2016), although our inferred numbers of haplotypes are larger. It should be noted that the original study only counted a sample as multiply infected if more than one allele peak was called from PCR-based fluorescent signals. Since we are considering the entire *P. vivax* genome instead of a set of microsatellite loci, our approach will naturally find more haplotypes. Whether the whole-genome haplotypes originated from a single or multiple infection is dependent on the genetic and transmission properties of *P. vivax* itself, and indeed, for any pathogen. The distribution of haplotype frequencies along the chromosomes is shown in **Supplementary** Notes **SECTION VI Fig. 9**.

To compare the *de novo* reconstructed haplotypes with strains inferred by a template-based method in metagenomics, we reanalyzed a meta-genomics dataset collected from a gastrointestinal microbiome undergoing shifts in species and strain abundance (Sharon, et al. 2013). The original publication suggested that the abundance of *Staphylococcus epidermidis* is primarily controlled by phages 13, 14 and 46 through the *mecA* gene. Based on StrainEst (Albanese and Donati 2017) (supported by templates of known strains), other researchers analyzed the same data and inferred the identities and frequencies of the three strains in question (Albanese and Donati 2017), which we were able to replicate (**Fig. 5a**). We have re-analyzed the same data by reconstructing fine-scale haplotypes. The *Staphylococcus epidermidis* genome was divided into 110 fragments (100 SNPs per fragment). The average number of haplotypes for each fragment was 9.3, although this value changed at different time points (**Supplementary Notes SECTION VI Table 4**). All fragmentary haplotypes are aligned back to the three main strains (**Supplementary Notes SECTION VI**) to examine the aggregated haplotype frequencies of each strain. By averaging all 110 regions, the aggregated frequencies (from PoolHapX) were found to follow the same pattern of changes as these three strains (**Fig. 5b**). This demonstrates that PoolHapX correctly identified haplotypes through *de novo* inference, without the use of reference templates from known strains, as required by StrainEst.

**Figure 5.**
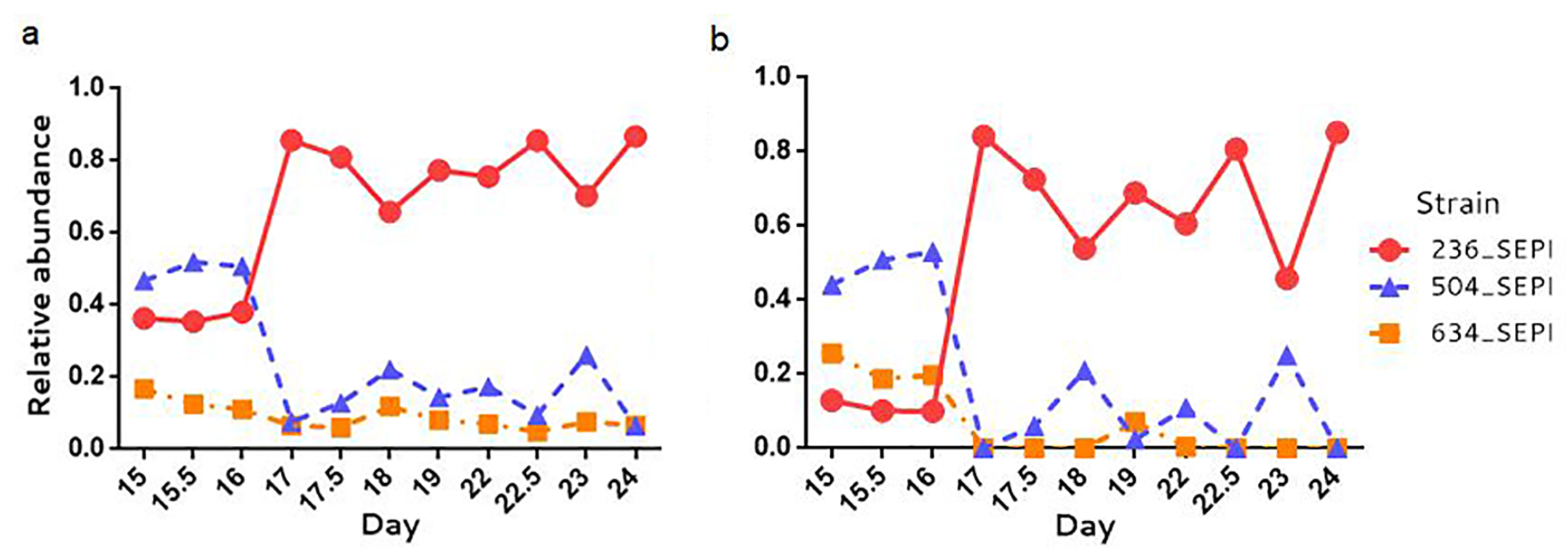
Staphylococcus epidermidis strain abundance calculated de novo by PoolHapX (a) and StrainEst based on templates of known strain (b) for early stages of infant gut colonization. All haplotypes predicted by PoolHapX are aligned to the three strains and we observe the same pattern of the changes of these three strains.

To demonstrate how PoolHapX can be used to discover novel evolutionary events, we tested PoolHapX on bulk-sequencing data from a recent intra-patient HIV study (Zanini, et al. 2015). This dataset contains longitudinal samples from multiple time points for 10 patients. We analyzed patient #1, which contains the most time points (12). We inferred haplotypes using PoolHapX, and observed 2 main haplotypes at time point 1, and 10 −13 main haplotypes at the rest time points (**Supplementary Notes SECTION VI Table 5**). We then calculated several extended haplotype homozygosity-related (EHH) summary statistics (Sabeti, et al. 2002), which measure linkage disequilibrium across a population by quantifying the probability that two randomly chosen particles are identical by descent in a certain region (see rationale in **Supplementary Notes SECTION VI**). Outlier values of the area under the EHH curve indicate that selective sweeps may have occurred (**Supplementary Notes SECTION VI**). While the size of linkage blocks decayed extremely rapidly post-infection in all genes (**Fig. 6a, b, Supplementary Notes SECTION VI Fig. 10**), it did not decrease monotonically as the HIV population adapted to the within-patient environment. To further quantify the rate and dynamics of selection within each gene, we plotted the size of windows with EHHS ≥ 0.5 at all time points and for multiple genes in the reconstructed haplotypes. The genes *gag*, responsible for assembly and structure, and *pol*, responsible for genetic reproduction (Konnyu, et al. 2013), are pictured in **Fig. 6c,d**, (other genes in **Supplementary Notes SECTION VI Fig. 11**). Within *gag* and *pol*, there was substantial heterogeneity in average window size over time, with the downstream regions of *gag* and *pol* largely fluctuating between 0 and 250 bp (**Fig. 6c, d**). These regions were highly conserved due to their respective roles in the HIV life cycle (Mayrose, et al. 2013).

**Figure 6.**
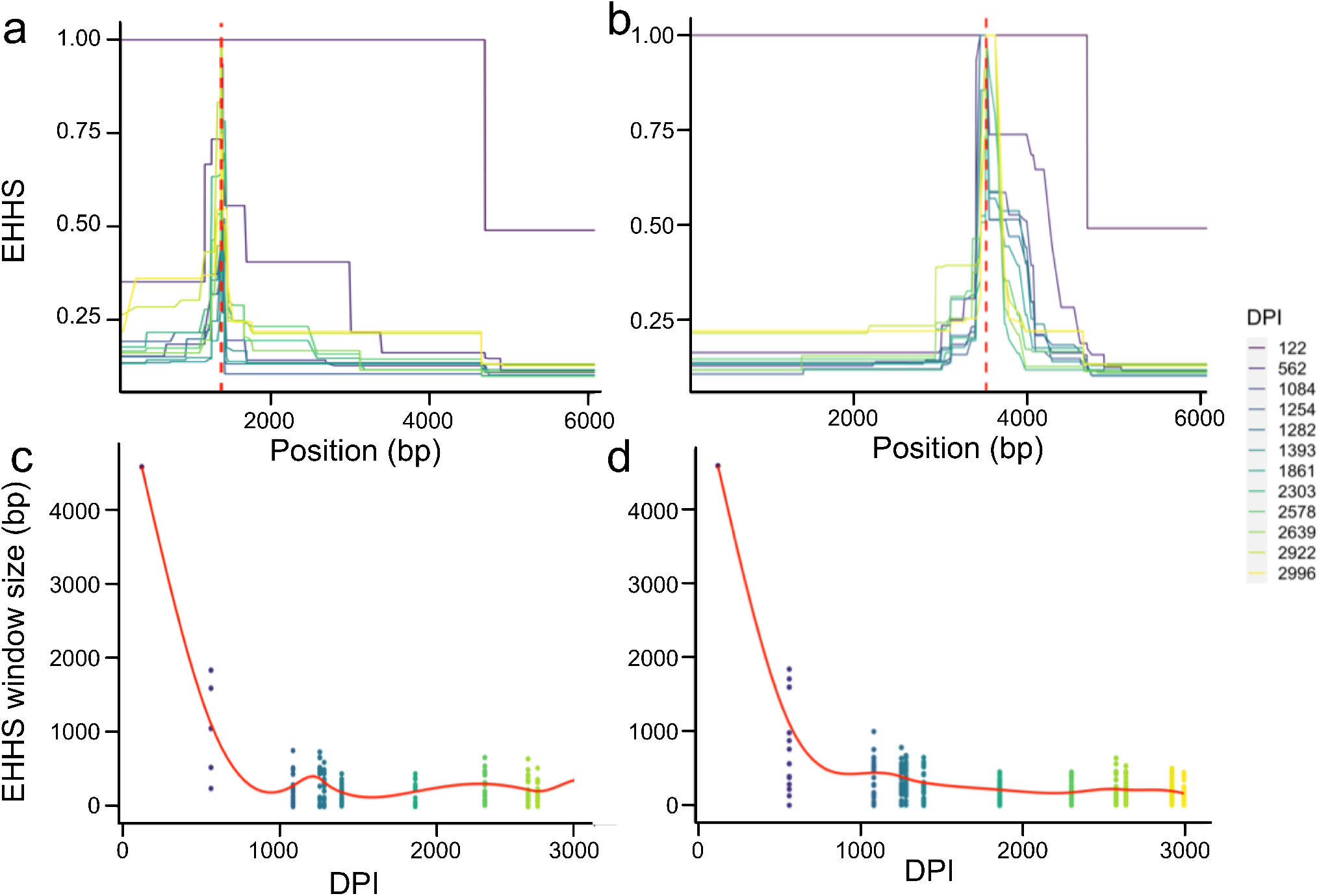
(a-b) The decay of EHHS around each SNP position in reconstructed HIV-1 haplotypes occurs rapidly during the acute phase of infection. The dashed red line indicates the location of the focal SNP position. (a) Position 1377 (Gag gene, found in p2 protein). (b) Position 3530 (Pol gene, found in p15). (c-d) The size of windows of EHHS ^3^ 0.5 fluctuate around gene-specific averages. The solid red line indicates a weighted mean across the positions in the gene. DPI refers to estimated days post-infection. Each dot represents the window size around at least one position. (c) Gag. (d) Pol. The legend (right) indicating the color corresponding to each time-point is common to panels a-d.

We have conducted a search for regions of positive selection between reconstructed haplotypes at adjacent time-points, where selective sweeps could have taken place. There are regions that are recurrently swept, most notably in the region of the *gag* polyprotein gene that encodes the p24 protein (**Supplementary Notes SECTION VI Table 7**). The occurrence of sporadic but re-occurring selective sweeps in gag, specifically p24, can be attributed to the appearance of cytotoxic T-lymphocytes (CTL) escape mutations, which reduce the ability of CTL to target virus-infected cells (Prince, et al. 2012). However, these escape mutations also decrease the replicative capacity of the virus, and a larger mutational burden corresponds to a greater decrease in capacity (Chopera, et al. 2008; Wright, et al. 2012). As such, episodic periods of positive selection at the same location would allow successful escape mutations to rise to fixation occasionally, while still allowing for genetic diversity to accumulate between selective sweeps.

## DISCUSSION

PoolHapX seamlessly integrates statistical and physical linkage information into a flexible but powerful framework for haplotype reconstruction. We have shown that PoolHapX produces more accurate haplotype reconstructions and frequencies than any other tool to date, and is robust to dynamics generated by within-host evolution. From the analysis of *Plasmodium vivax, Staphylococcus epidermidis* and HIV data, we show that PoolHapX is scalable, accurate, and infers haplotype data that is valuable for understanding the within-patient diversity of pathogens.

One of our main algorithms, AEM is borrowed from human genetics. In early genome-wide association studies (GWAS) for humans, DNA from multiple individuals was artificially pooled to save on genotyping costs. Subsequently, *in silico* methods were applied to the pooled sequencing data to reconstruct haplotypes (Kuk, et al. 2009; Pirinen 2009) for association mapping. Though technological advancements have made this cost-saving practice unnecessary, as a theoretical assessment, we compared PoolHapX against the GWAS-based haplotype reconstruction tools Hippo (Pirinen 2009) and AEM (Kuk, et al. 2009). Since there are many publicly available human genomes we did not simulate haplotypes, instead making artificial pools using phased haplotypes from the 1000 Genomes Project (Consortium 2015) (**Supplementary Notes SECTION III**). **Supplementary Notes SECTION VII Fig. 14** show that PoolHapX slightly outperformed alternative tools when there were relatively few SNPs. When there were many SNPs (>=25) in a region, however, the other tools did not finish in two weeks (using an HPC node with 48Gb memory), while PoolHapX could still produce reliable results with a large region containing as many as 200 SNPs in a few hours.

The main novelty of PoolHapX stems from the sharing of haplotype information across pools. PoolHapX performance is sensitive to the number of pools and number of haplotypes. Accordingly, we have investigated this issue using HIV as an example. By fixing the number of global haplotype as 25 and specifying the number of simulated pools from 3 to 100, we conducted analysis and used both MCC/JSD and CRR/FDR/F1-score to assess the decrease of accuracy proportional to the number of pools (**Supplementary Fig. 22**). It is observed that the performance may not be satisfactory when the number of pools equals or is lower than 5 (**Supplementary Fig. 22**). Similarly, we conducted simulations by fixing the number of pools to 25 and looked at the impact of the number of haplotypes (**Supplementary Fig. 23**). It appears that the accuracy is satisfactory when the number of global haplotypes equals or is lower than 40. These results provide some guidelines for the analysis of viral genomic data.

This implementation of PoolHapX has some limitations. We found that the method is sensitive to the inferred within-host allele frequency, and therefore high variance in allele frequency caused by very low sequencing coverage will result in high error rates. The performance of PoolHapX is also variable when we attempt to infer frequencies of more than 50 haplotypes in the pools. However, if we aim only to assess a smaller number of more abundant haplotypes (e.g. 10-20), it is robust to noise caused by rare haplotypes (**Fig. 2** and **Fig. 3**). Another limitation is that PoolHapX is not able to handle very large structural variants for the moment, while small indels can be handled in the same way as point mutations (SNPs). PoolHapX does not take the quality score of variant calls into account when reconstructing haplotypes, though several tools do (Prabhakaran, et al. 2013; Ahn, et al. 2018; Ahn and Vikalo 2018)

At present, PoolHapX is in continuing development, with ongoing work to integrate third-generation sequencing data (Check Hayden 2009) into PoolHapX, as well as algorithms using genomic assembly (Bankevich, et al. 2012) to improve haplotypes.

## MATERIAL AND METHODS

### Graph coloring algorithm

If two sequencing reads cover the same genetic segregating site but carry different alleles, it is certain that they do not belong to the same haplotype. Based on this observation, we build a graph <V, E> in which every read is a node v. For two nodes v_1_ and v_2_, we put an edge e_1,2_ between them if and only if we have information to claim they are not on the same haplotype. Then, the haplotyping problem becomes a graph-coloring problem: where each node (i.e. each read) is assigned a color, such that nodes linked by edges are colored differently. This ensures that reads belonging to different haplotypes are colored differently. After conducting this graph-coloring problem, we effectively estimate haplotypes by collecting reads of the same colors. As the standard parsimony algorithm is too slow when the number of reads is large, we have implemented a greedy algorithm to color this graph (**Supplementary Notes SECTION II**). The spatial complexity of our graph coloring is O(n^2^) and the time complexity is O(n^2^). The outcome of graph-coloring forms starting states for the whole pipeline in two respects. First, by collecting all reads of the same color, PoolHapX can generate segments of local haplotypes as the initial input to the next step, the Hierarchical AEM algorithm. Second, the gaps between local haplotypes naturally inform the initial divide & conquer plan for subsequent steps (**Supplementary Notes SECTION II**).

### Hierarchical AEM algorithm

The basic version of AEM algorithm, as described in (Kuk, et al. 2009), builds upon the multivariate normal (MVN) distribution. The LD between all pairs of *n* segregating sites is modeled as the variance-covariance matrix of an MVN distribution. Initially, all *2*^*n*^ possible haplotypes will be assigned to the same frequency (*1/2*^*n*^). Then in an iterative procedure, the likelihood ratio of observing the data with or without the presence of each haplotype is estimated using the MVN densities. These ratios are called “Importance factors” (Kuk, et al. 2009), indicating the importance of the individual haplotypes, and will be used to adjust their haplotype frequencies. This adjustment is conducted iteratively until the haplotype frequencies converge (**Supplementary Notes SECTION II**).

In our adaptation of AEM we have made the following three modifications. First, the initial haplotype configuration is no longer a uniform distribution of all *2*^*n*^ haplotypes. Instead, using the haplotypes gained from graph coloring, haplotypes with higher sequence coverage start with a higher initial frequency. As a consequence, many potential haplotypes that are not observed in graph coloring will have zero frequency. While the spatial complexity of AEM remains O(*n*^*2*^) and the theoretical time complexity remains O(n^3^ x *2*^*n*^), the number of required iterations are substantially reduced in practice due to the initial configuration being closer to the truth. Second, we use a divide & conquer algorithm to scale up the original AEM algorithm to larger regions, so that we run AEM in a hierarchical manner. The shorter haplotypes inferred from the previous AEM iteration are used in the next round of AEM to form longer haplotypes (**Supplementary Notes SECTION II Fig. 3** and **Fig. 1a**). In each round, local regions are designed to have half of their segregating sites overlap with the next region (**Supplementary Notes SECTION II Fig. 5**), in order to form tiling windows that can be stitched together at the next hierarchical level. Third, the original AEM is not robust to numerical instability if the denominator in the likelihood ratio is close to zero. However, this problem occurs more frequently in larger regions with sparse non-zero LDs in the covariance matrix. We have fixed this by adjusting the calculation of the likelihood (**Supplementary Notes SECTION II**).

### Breadth-First Search (BFS)

The iterative AEM algorithm generates successively larger regional haplotypes until an upper limit is reached, which is 96 segregating sites by default. PoolHapX will then attempt to resolve global haplotypes. The outcome of the last AEM iteration is a set of local haplotypes that span tiling windows, with many potential combinations that form global haplotypes. To resolve global haplotypes, we model the local regions as a tree, with each local haplotype as a node. Haplotypes from the first region of the genome form the first level of the tree, while haplotypes from the next tiling region form the nodes of the next level. If two haplotypes in adjacent regions have the same alleles in their overlapping segments, we add an edge linking these two nodes. Traversing the resulting tree generates an exhaustive set of all plausible combinations of local haplotypes, which forms the set of candidate global haplotypes. We implement a standard Breadth-First Search (BFS) algorithm (Cormen, et al. 2009) to conduct this traversal. Finally, we filter out some global haplotypes that are inconsistent with the sequencing reads (**Supplementary Notes SECTION II**).

### Global Regularization model

Given all candidate global haplotypes from the BFS step, we use an innovative regression model to estimate the within-host global haplotype frequencies in each pool:

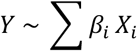

Where *β*_*i*_ is the frequency of the *i-*th global haplotype in the host (pool). Here *Y* and *X*_*i*_ are the independent and predictor variables in a standard regression model. We use two types of samples to train *Y* from *X*_*i*_, which represent two different sources of data: minor allele frequency and physical LD. Mathematically, the dimension of *Y* is *n + n(n-1)/2* (where *n* is the number of sites), representing the alternate frequency at *n* sites and their *n(n-1)/2* physical LD across pairs of sites observed in the reads. First, at each site, the sum of frequencies of haplotypes containing the alternate allele should be statistically similar to the observed alternate allele frequency based on reads from the pool. This is the same information utilized by several other tools (Pulido-Tamayo, et al. 2015; Albanese and Donati 2017). An innovation of our design is the use of a second type of sample: for each pair of sites, the sum of frequencies of haplotypes containing both alternate alleles should be equal to the frequency observed in the number of reads that cover both alternate alleles in the pool, which includes read-pairs and many barcoded reads in 10x linked-reads (**Supplementary Notes SECTION II Fig. 7**). A full description is in (**Supplementary Notes SECTION II**). The set of *β*_*i*_ that best fits these two sets of constraints is our solution.

To reduce overfitting, the objective function for training the above regression is designed as a combination of L0 and L1:

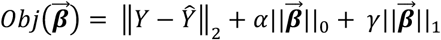

where 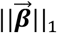 is the L1-norm, which is the sum of absolute values of all *β*_*i*_; and 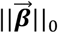 is the L0-norm, which is the number of non-zero *β*_*i*_.

Several existing papers use L1 regularization alone (Albanese and Donati 2017; Leviyang, et al. 2017) This strategy does not work for many haplotypes with small differences. This is because L1 penalizes the sum of absolute values of all regression coefficients, i.e., the haplotype frequencies in each pool. If the inference method is reasonably designed, however, the sum of haplotype frequencies in a host will always be near 1.0. This is because convex optimization-based penalties only prevent outcomes with negative frequencies, and do not distinguish between outcomes containing few haplotypes with large frequencies and outcomes containing many haplotypes with small frequencies. In essence, L1 does not produce sparse solutions when the difference between haplotypes are small, such as samples that arise when within-host recombination generates many similar haplotypes with small regions of genetic differences. To further enforce parsimony, adding a layer of L0 regularization further regularizes the number of haplotypes. This is why L0 regularization is necessary for our method, although it is much slower. Fortunately, The L0Learn package (Hazimeh and Mazumder 2018) a breakthrough in the field of computer science to solve L0-based optimization is available recently, which empowered PoolHapX to conduct this inference. Indeed, our preliminary development towards viral haplotype reconstruction in a single host (that does not utilize cross-host sharing as PoolHapX does) show that such L0+L1-regularization is very fast (Cao, et al. 2020). Finally, the cross-host (i.e., population) frequencies of each haplotype can be formed by combining the within-host frequencies.

### HIV evolutionary data analysis

For a description of Patient 1 data, the SNP position-calling pipeline, and haplotype reconstruction, see **Supplementary Notes SECTION VI**.

The R package rehh (version 3.0) was applied to survey linkage patterns within a single time-point, and changes to linkage patterns across the duration of infection monitoring (Gautier and Vitalis 2012). Several long-range haplotype-based evolutionary statistics related to extended haplotype homozygosity (EHH) (Sabeti, et al. 2002) were used to quantify the type and magnitude of selection. To search for regions of positive selection within the reconstructed genome, integrated haplotype score (iHS) (Voight, et al. 2006) and cross-population EHH scores (XP-EHH) were calculated for each time-point and between each time-point, respectively.

For more details about the rationale behind each layer of analysis, see **Supplementary Notes SECTION VI**. The scripts that generate MS (Ewing and Hermisson 2010) output format from PoolHapX output files and apply EHH-based statistics to the reconstructed haplotypes are available at (https://github.com/theLongLab/PoolHapX/tree/master/Simulation_And_Analysis/HIV_analysis_code). Parameters and settings are described in further detail within the scripts.

### Other data analyses

Processing and analyses of *Plasmodium* and other metagenomic data (*E. Coli*) can be found in **Supplementary Notes SECTION VI**. Details of simulations and comparisons (including how other tools are executed) are also included in **Supplementary Notes SECTION III and SECTION IV**.

## AVAILABILITY

PoolHapX is an open source collaborative initiative available in the GitHub repository (https://github.com/theLongLab/PoolHapX)

## SUPPLEMENTARY DATA

Supplementary Data are available at MBE Online.

## ACKNOWLEDGEMENT

*Author contributions*: Conceived the project: QL and DJ; Designed the algorithms: QL, CC, and JW; Implemented the software: CC and QL; Tested the software: LM, ML, DP and SG. Simulated data analyses: CC and JH; Real data analyses: LM, CC, and DP; Contributed data and advice: DJ, TM, SM, LS, MG, GY; Fine-art figures: DK; Write the manuscript: QL, CC, LM, JH, and DK, with contributions from all co-authors.

