## SUPPLEMENTARY MATERIALS for "Reconstruction of microbial haplotypes by integration of statistical and physical linkage in scaffolding"

### TABLE OF CONTENTS

**SECTION I: OVERVIEW OF HAPLOTYPE RECONSTRUCTION METHODS ... 3**

**SECTION II: POOLHAPX ALGORITHM ... 4**

Step 1: Graph Coloring ... 4
Step 2: Hierarchical AEM ... 5
Step 3: Breadth-first Search ... 9
Step 4: L0 and L1 Regularized Regression ... 9

**SECTION III: SIMULATION PROCEDURES ... 12**

Simulating Haplotypes for Each Pool Using SLiM ... 12
*Viral haplotype simulation* ... 13
*Bacterial haplotype simulation* ... 13
*Metagenomic haplotype simulation* ... 13
*Haplotype simulation under evolutionary scenarios* ... 14
*Haplotype simulation when provided with single-cell linked-reads* ... 14
*Example of a SLiM script* ... 14
Human Haplotype Preparation ... 17

**SECTION IV: ANALYTIC PROCEDURE ... 19**

Processing SLiM Output ... 19
Sequence Alignment and Variant Calling ... 20
PoolHapX Analysis ... 22
*Filtering of genetic variants* ... 22
*Comparison criteria* ... 22

**SECTION V: RUNNING OTHER HAPLOTYPE RECONSTRUCTION TOOLS ... 24**

Viral Quasispecies Reconstruction Tools ... 24
Bacterial Haplotype Reconstruction Tools ... 27
Metagenomics Haplotypes Reconstruction Tools ... 28
Human Haplotypes Reconstruction Tools ... 29
Processing Haplotype Reconstruction Output ... 30

**SECTION VI: REAL DATA ANALYSES ... 32**

Plasmodium vivax Analysis ... 32
Infant Gut Colonization Time Series Analysis ... 35
HIV Within-Patient Evolutionary Analysis ... 37
Longitudinal Analyses of Patient HIV Populations ... 40

**SECTION VII: OTHER SUPPLEMENTARY FIGURES ... 46**

**SECTION VIII: REFERENCES ... 59**

#### SECTION I: OVERVIEW OF OTHER HAPLOTYPE RECONSTRUCTION METHODS

The reconstruction of maternal and paternal haplotypes from diploid genotype data has excellent precision in human genetics, due to modelling recombination and coalescence(Stephens and Donnelly 2003; Scheet and Stephens 2006; Howie, et al. 2012). Large reference panels in humans(Kowalski, et al. 2019) allow routine phasing and imputation of long-range haplotypes.

However, this procedure does not extend to the inference of many haplotypes in the same host(s), or to microbial species that are not extensively studied. To facilitate the analysis of data generated by genotyping artificially pooled samples (a cost-saving strategy used in Genome-Wide Association Studies in the 2000s), several methods were developed to infer cross-pool population haplotype frequencies using pooled genotyping data. The standard PHASE algorithm (modeling coalescence)(Pirinen, et al. 2008), expectation minimization (EM) algorithm (modeling general sharing), Markov-chain Monte Carlo (MCMC) sampling (modeling both coalescence and recombination)(Kuk, et al. 2009; Pirinen 2009), and phylogenetic trees (modeling coalescence)(Efros and Halperin 2012) have been developed to reconstruct haplotypes *in silico*. These methods in general do not scale to large haplotypes with more than 25 genetic variants and cannot be applied to microorganisms.

To deconvolve viral and bacterial pooled-sequencing data, researchers have developed many tools using different computational techniques including *de novo* assembly (utilizing sequencing reads)(Baaijens, et al. 2017), tensor decomposition (utilizing sequencing reads and allele frequency)(Hashemi, et al. 2018), graph-based algorithms (utilizing sequencing reads)(Mazrouee and Wang 2014), and regularization (utilizing allele frequencies and template genomes)(Albanese and Donati 2017). As a representative technology for barcoded reads linked-reads, the 10X Genomics has provided a built-in tool for inferring paternal and maternal haplotypes and large structural variants, Long Ranger, that achieves the goals by utilizing sequencing reads that spans a large distance(Zheng, et al. 2016).

In summary, the first category of models assumes cross-host sharing via statistical models in the domain of “genetics” using statistical linkage disequilibrium; while the second focuses on within-host specifics observed by sequencing reads using “genomics” in the form of physical linkage disequilibrium.

#### SECTION II: POOLHAPX ALGORITHM

##### Problem formulation

The haplotype reconstruction problem to be addressed in this paper is intuitively outlined below: The input information of the algorithm are (1) in-pool allele frequencies in each pool, recorded in the VCF files (one VCF file for each sample); and (2) reads that are aligned to the reference genome (of the focal species) carrying multiple mutations, recoded in the BAM files. The output will be the identity of global haplotypes and their in-pool as well as across-pool frequencies.

##### Step 1: Graph coloring

The haplotyping problem is initially interpreted as a graph coloring model as follows. We form a graph <V, E> in which V denotes all nodes in the graph and E denotes the edges between nodes. Each sequencing read represents a node in the graph. An edge will be placed between two nodes if their corresponding reads have an overlapping region which contains at least one varying site between the two reads. Since the nodes linked by an edge represent reads which differ in at least one site, they must belong to two different haplotypes. Therefore, if we assign colors to each node so that every edge joins a pair of differently colored nodes, then the nodes with the same colors may belong to the same potential haplotypes. Solving this graph-coloring problem gives a preliminary haplotyping solution, although the solution is very approximate when there is insufficient read depth to distinguish between multiple valid graph-colorings.

We implemented two algorithms to solve the graph-coloring problem. We first attempted to convert this problem to a standard maximum clique problem(Bomze, et al. 1999) in graph theory in order to use miniSAT(Sorensson and Een 2005) to solve the problem. This is a parsimony algorithm that aims to search for the minimal number of haplotypes, but becomes very slow when the number of reads is large. However, the parsimony property is undesirable, since the graph-coloring step only provides the initial input for our downstream hierarchical AEM algorithm (which integrates statistical linkage disequilibrium, or LD). As a result, we have implemented a greedy algorithm as the default graph-coloring method in the current version of PoolHapX, which is described below.

During computation we maintain three data structures: the color of each node, the colors that have been assigned, and the set of colors of the neighbors of each node. Let C = {1, 2, ..., M} represent the current M colors that have been assigned to at least one node, and let color(x) denote the color of a node x. If a node has not been assigned a color, color(x) = 0. We use nbCol(x) to denote the set of different colors among all the neighboring nodes of x and size(nbCol(x)) to denote the size of this set. Our algorithm proceeds as follows: we begin by assigning color 1 to a random node. In each subsequent step, we select an uncolored node x* with the largest number of differently colored neighbors size(nbCol(x*)), and assign it to the color color(x*) = min{C \ nbCol(x)}. If there are no free colors which can be assigned to x*, meaning size(nbCol(x*)) = M, we add a new color M+1 to C and assign it to the node. We then update every uncolored node y with an edge to x so that nbCol(y) contains color(x*). This process is repeated until all nodes are colored (**Supplementary Fig. 1**).

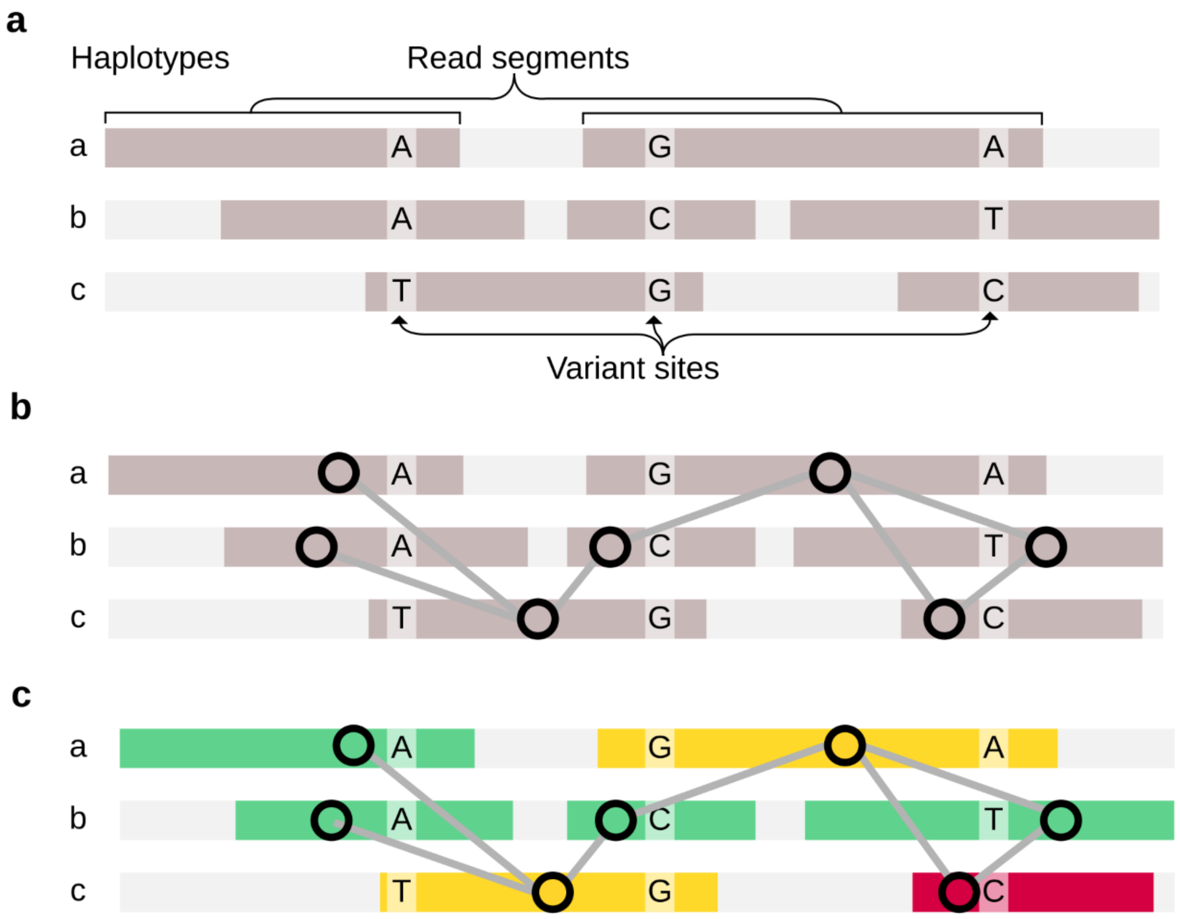


**Supplementary Figure 1**: An example of the Graph Coloring algorithm. Each node in the graph is a sequencing read (panel **a**). Two nodes are joined by an edge if their corresponding reads overlap in at least one position with differing alleles (panel **b**). A color is assigned to each node so that any two nodes linked by an edge are colored differently. As a result, nodes with the same color potentially belong to the same haplotype (panel **c**).

#### Step 2: Hierarchical AEM

The basic approximate expectation maximization (AEM) algorithm as described in Kuk *et al* (Kuk, et al. 2009) utilizes the multivariate normal (MVN) distribution. The LD between all pairs of $n$ genetic variants is modeled as the variance-covariance matrix of a MVN distribution. Suppose there are $q$ genetic variants, and therefore $2^q$ possible haplotypes. Initially, all $2^q$ possible haplotypes are assigned the same frequency $1/2^q$. Then, using an iterative procedure, the ratio of the likelihoods of observing MVN densities with or without each haplotype is estimated. These ratios indicate the relative importance of each haplotype, and are used to adjust the frequencies of all the $1/2^q$ possible haplotypes. This adjustment is conducted iteratively until the haplotype frequencies converge. A detailed mathematical formulation is outlined below (please refer to Kuk *et al* (Kuk, et al. 2009) for full description and justification).

Suppose we have $q$ genetic variants and $T$ hosts. We use $\boldsymbol{A}$ to denote the matrix of aggregated variant allele frequencies in all the pools. That is, $A_{i,j}$ denotes the aggregated allele frequency of the $j$-th genetic variant in the $i$-th pool. Note that this is the only input for the original AEM algorithm, and additional information from the sequencing reads is not integrated. Denoting $r = 2^q$ as the total number of haplotypes, let $H = (h_1, h_2, \ldots, h_r)$ be the sequence of all potential haplotypes. Using the initial haplotype frequencies $P = (p_1, p_2, \ldots, p_r)$, we calculate the mean $\mu$ and variance-covariance matrix $\Omega$ of a MVN distribution modeling the $q$ variants using the formulas: $\mu = \sum_{j=1}^{r} p_j h_j$ and the variance-covariance matrix $\Omega = \sum_{j=1}^{r} h_j h'_j p_j - \mu\mu'$ . Then the importance factors (IFs)(Kuk, et al. 2009) indicating the importance of each haplotype is defined as:

$$E[m_{ij}|A_i; \hat{\mathbf{p}}^{(t)}] \approx 2N\hat{p}_j^{(t)} \frac{\phi\left(A_i - h_j; (2N-1)\hat{\mu}_j^{(t)},\ (2N-1)\widehat{\Omega}^{(t)}\right)}{\phi(A_i; 2N\hat{\mu}^{(t)}, 2N\widehat{\Omega}^{(t)})} = \widehat{m}_{ij}^{(t)}$$

where $\phi$ denotes the multivariate normal density.

This IF is then used to update the haplotype frequencies:

$$\hat{p}_j^{(t+1)} = \frac{\sum_{i=1}^{T} \widehat{m}_{ij}^{(t)}}{2NT}$$

In our variant of the AEM algorithm, we have made the following three modifications.

First, the initial haplotype configuration is no longer a uniform distribution over all $2^q$ possible haplotypes. Instead, using the potential haplotypes determined by the Graph Coloring step, we assign higher initial frequencies to haplotypes with higher sequence coverage. Many potential haplotypes that are not observed in the graph coloring step, therefore are removed. While the theoretical spatial and time complexity of our AEM remains $O(q^2)$ and $O(q^3 \times 2^q)$ respectively, our method substantially reduces the number of iterations required in practice since our initial frequencies start much closer to the true values. As our Graph Coloring algorithm is a greedy algorithm that may produce too many haplotypes (especially when the sequencing coverage is not high), we further filter out haplotypes that are not supported by information from the sequencing reads. By default, PoolHapX requires each haplotype to have at least 2% coverage by sequencing reads at every pair of adjacent sites (**Supplementary Fig. 2**).

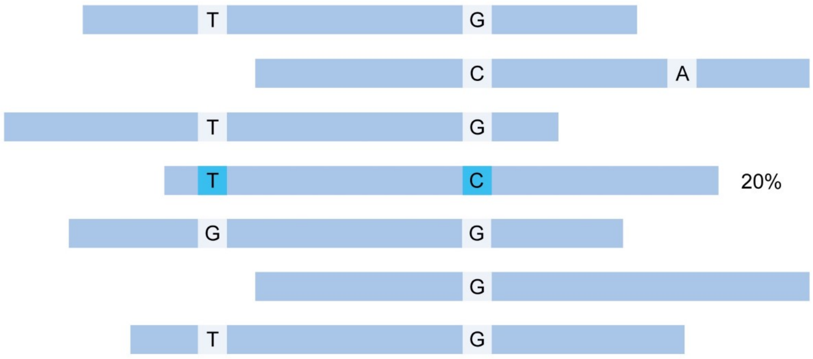


**Supplementary Figure 2**: In this example there are five reads covering two adjacent sites with variants (T/G) and (G/C) respectively. Since there is only one read with variants "T" and "C" (light blue), the percentage of T-C haplotype supported by physical LD is 20%.

Second, we apply AEM hierarchically, by using a divide-and-conquer algorithm to scale up the original AEM algorithm to larger regions. The frequency and identity of shorter haplotypes inferred by an iteration of AEM are combined into longer haplotypes for subsequent iterations (**Supplementary Fig. 3)**. In each iteration, the local regions on which AEM operates are designed to have half of their genetic variants overlap with the next region (**Supplementary Fig. 3**). This allows the local regions to form tiling windows that are stitched together at the next hierarchical level.

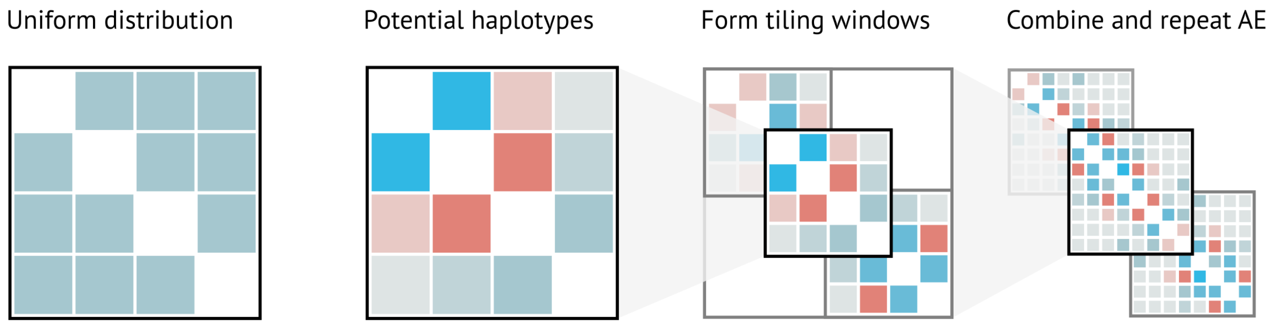


**Supplementary Fig. 3**: Divide-and-conquer algorithm to scale up the original AEM algorithm to larger regions. The darkness of the blue color indicates the strength of statistical LD.

By default, PoolHapX undergoes four iterations of the divide & conquer procedure. The division plan for Level-1 of the AEM procedure is based on the initial blocks determined by the outcome of the Graph-Coloring step (**Supplementary Fig. 4**). The gaps between graph-coloring haplotypes indicate sites with very little physical LD (mostly due to sparse sequencing coverage), and are used to divide the sequence into regions. The midpoints of successive regions are used to determine the start and endpoints of additional, overlapping regions. The combination of these overlapping regions forms the tiling regions of Level-1. This design specifically leverages statistical LD to stitch the gaps left by physical LD.

PoolHapX provides options for the users to control the size of Level-1 regions so that it is not solely decided by the positions of gaps in the physical LD (**Users Manual**). PoolHapX will then run AEM against the regions independently. After resolving the Level-1 haplotypes, adjacent regions are combined into a larger region for the Level-2 partition. Only combinations of Level-1 haplotypes with non-zero frequencies are considered as candidate haplotypes for Level-2. Specifically, adjacent regions are stitched together by the following procedure: Suppose we have 3 tiling regions, *A*, *B*, and *C*, as depicted in (**Supplementary Fig. 5**), and *a, b, c* are three haplotypes in region *A; d, e*, are two haplotypes in region *B*, and *f, g, h* are three haplotypes in region *C*, respectively. The smaller haplotypes form a longer Level-2 haplotype (such as *ah*), only if the number of mismatched alleles between the haplotype in *A* and haplotype in *B* is smaller than a cutoff value, and likewise between *B* and *C*. The user can specify this cutoff (**Users Manual**) based on their understanding of the haplotype structure and level of polymorphism for their focal species. Using the same procedure, PoolHapX can run additional rounds of the AEM algorithm, depending on the availability of the users' computing resources. In general, the size of region that is resolvable by AEM is limited by the computation cost of storing and factoring many $q \times q$ matrices. By default, PoolHapX can process regions with up to 112 genetic variants after four levels of iteration.

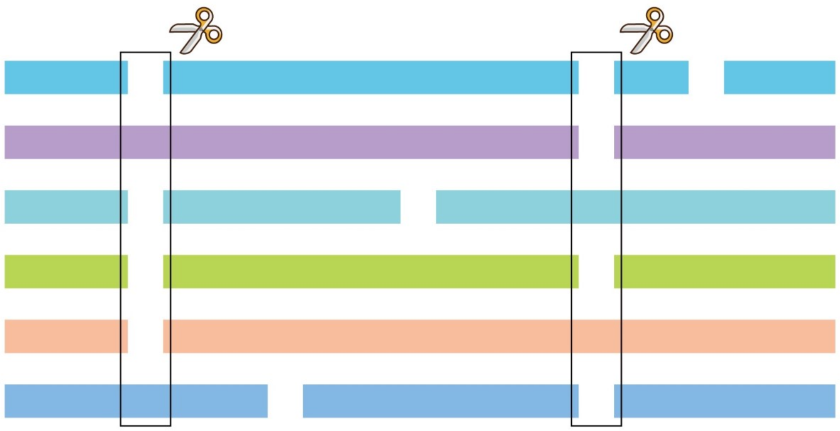

**Supplementary Fig. 4:** Regions are split at sites that have a higher number of gaps (shown as rectangles under the scissors) between graph-coloring haplotypes, since the gaps contain little information about the physical LD.

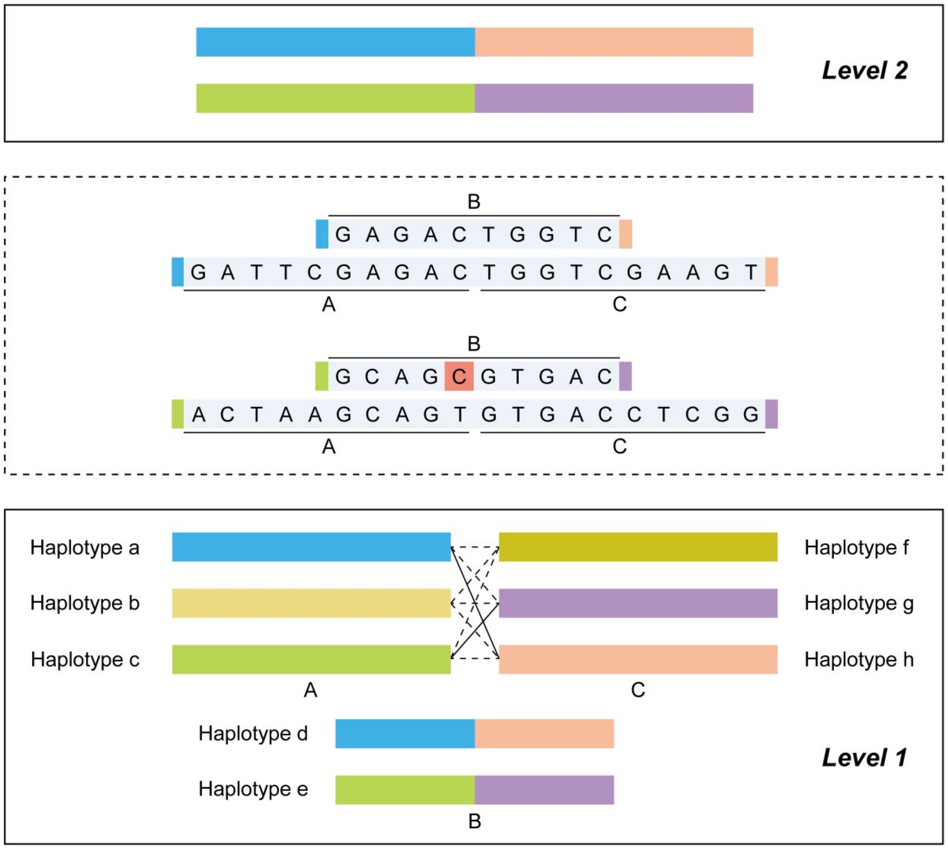


**Supplementary Fig. 5:** Three Level-1 tiling regions generate one larger Level-2 region according to the number of mismatched alleles. Suppose that Level-1 regions *A*, *B*, and *C* contain haplotypes (*a*, *b*, *c*), (*d*, *e*), and (*f*, *g*, *h*) respectively. Regions A and C can be combined in nine possible ways (*af*, *ag*, *ah*, *bf*, *bg*, *bh*, *cf*, *cg*, *ch*). However, based on their overlap with region B's haplotypes, as indicated by the colors of haplotypes *d* and *e*, only two combinations (*ah* and *cg*) are retained in Level-2, as the number of mismatched alleles in the other combinations exceeds the cutoff point.

Third, the original AEM is not robust to numerical instability if the denominator in the likelihood ratio is close to zero. When this happens, the program will converge rapidly to a sink point at which only one haplotype is dominant. This is not an important issue in the original AEM design where the number of variants is small. However, this problem occurs more frequently when applying to AEM to a larger matrix containing sparse non-zero LDs. We have resolved this problem by adjusting the calculation of the likelihood so that if both the denominator and numerator are very close to zero, the Importance Factor (IF) value will be set to a moderate constant. If only the denominator is close to zero, the IF value will be set to a large constant. These parameters are tunable by the end user.

##### Step 3: Breadth-first search

The top-level local haplotypes produced by the hierarchical AEM algorithm will be linked into global haplotypes using a breadth-first search (BFS) algorithm. As haplotypes were inferred for each of the tiling windows, the haplotypes from adjacent windows will overlap over about half of their sequence length. If the sequences of two overlapping haplotypes match, they are stitched together to form a longer haplotype. Since the overlapping regions may not be identical due to errors in the previous steps, two haplotypes may be stitched together as long as the number of mismatches in their overlapping region is less than a user-specified cutoff parameter (**Supplementary Fig. 6**). By default, PoolHapX repeats this process to form local haplotypes with a maximum of 112 genetic variants.

Next, global haplotypes are determined by traversing a tree structure which contains the locally stitched haplotypes. The *i*-th level of the tree represents the *i*-th genomic region. At each level, local haplotypes are represented by a node in the tree. We draw an edge between two nodes in neighboring levels if and only if their corresponding haplotypes can be stitched together (**Supplementary Fig. 6**).

By collecting all the paths that traverse the tree, we find the set of global haplotypes that are consistent with the outcome of the hierarchical AEM algorithm. We have implemented a standard breadth-first search (BFS) algorithm(Cormen, et al. 2009) to traverse the tree and produce a set of candidate global haplotypes for the final step.

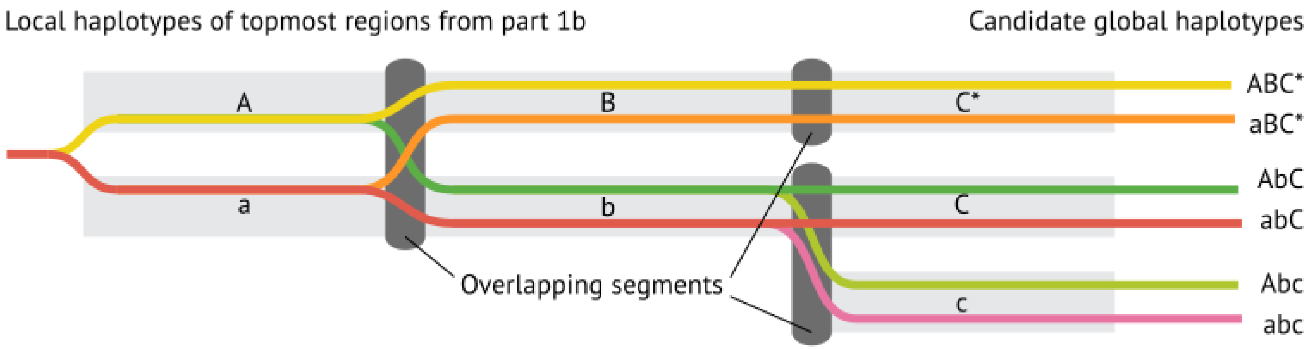


**Supplementary Fig. 6:** An example of the BFS algorithm used to determine global haplotypes. Haplotypes from adjacent windows are stitched together to form a longer haplotype if the overlapping sequences of two adjacent haplotypes match.

**Step 4: L0 and L1 regularized regression**

Using the candidate global haplotypes from the BFS algorithm, we use an innovative regression model to estimate the frequencies of within-host global haplotypes from two sources of information. First, the within-host aggregated minor allele frequency (MAF) at a given site should match the sum of the frequencies of haplotypes bearing the alternate allele at that site. This is the basis for many algorithms such as StrainEst and others(Pulido-Tamayo, et al. 2015; Albanese and Donati 2017). Second, the physical LD revealed by the sequencing reads is also integrated into the model. Specifically, consider the alternate allele (coded as 1 in our model) at a pair of variant sites *t* and *k* which are not necessarily adjacent. The sum of the frequencies of haplotypes carrying alternate alleles at both *t* and *k* should match the frequency of reads covering *t* and *k* which also carry both alternate alleles. This insight is generalizable to any sequencing technology, including standard Illumina paired-end reads, barcoded linked-reads, and third-generation long-reads.

Based on this rationale, we define the following model:

$$Y \sim \sum \beta_i X_i$$

This is a standard regression model where $Y$ and $X_i$ are independent variable and predictors. The coefficient $\beta_i$ is the frequency of the *i*-th global haplotype in the host (pool). Note that $Y$ and $X_i$ represent the two different sources of data discussed above, although denoted by the same term in the model. That is, we use two types of samples for $Y$ and $X_i$ to train this model. For individual MAFs, at each site *j* the sum of haplotype frequencies containing the minor allele at *j* should match the MAF of the observed reads at site *j*. This is the same information that has been utilized by several other tools(Pulido-Tamayo, et al. 2015; Albanese and Donati 2017). Additionally, our design utilizes physical LD information as follows: for each pair of sites *j* and *k*, the sum of haplotype frequencies containing both alternate alleles should match the frequency of observed reads containing alternate alleles at *j* and *k* (**Supplementary Fig. 7**). The dimension of $Y$ is *n* + *n(n-1)/2*, where there are *n* individual sites and MAFs, and *n(n-1)/2* pairs of sites with physical LD information. An innovation of our design is the use of the second type of sample: for each pair of two sites, the sum of frequencies of haplotype that contains both alternate alleles should be equal to the frequency of observing the number of reads that cover both alternate alleles in the pool (**Supplementary Fig. 7**). In order to adjust the relative importance of MAF versus physical LD information, the weighting of each type of information is tunable in the regression model (**Users Manual**). The set of global haplotype frequencies (i.e., the set of $\beta_i$) that best fits these two sets of constraints is our solution.

The set of $\beta_i$ that fits the above two sets of constraints best is our solution.

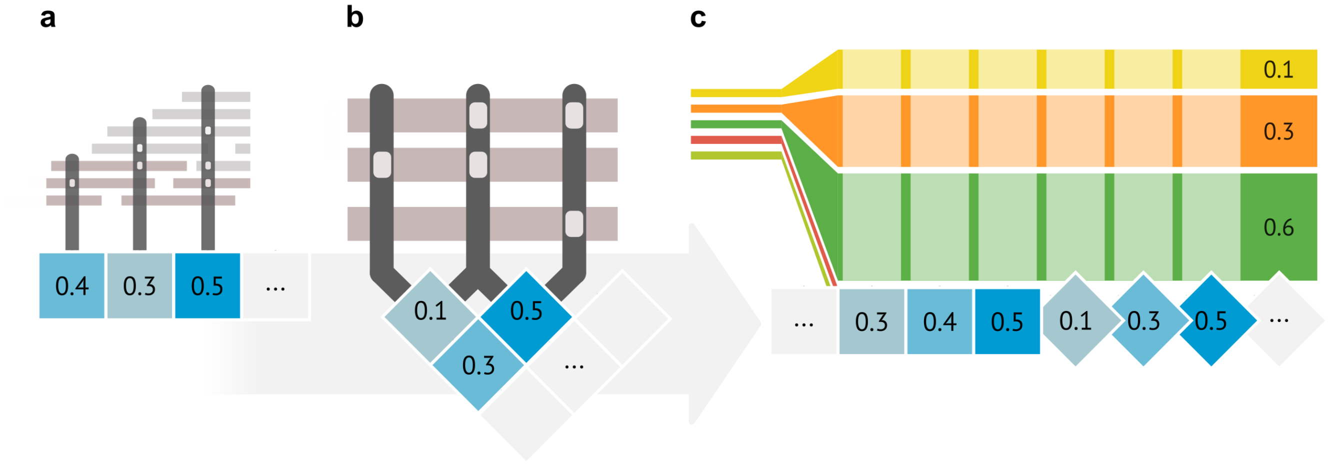


**Supplementary Fig. 7:** PoolHapX uses two types of frequency information in the L0 and L1 regularized regression step: the sum of minor allele frequencies at individual sites (panel A), and the sum of two-variant alternate allele frequencies at paired sites (panel B). Here three genetic variants provide six samples to train the model: three within-host aggregated MAFs represented by blue squares, and three pair-wise physical LDs represented by blue diamonds. A regularized (L0 and L1) optimization is used to estimate haplotype frequencies. Panel C illustrates the output of the regression model, which contains three non-zero frequency haplotypes represented by the yellow, orange and green bars. The remaining potential global haplotypes have zero frequencies, indicating that they do not exist in this pool.

To reduce overfitting, the objective function for training the above regression combines L0 and L1 regularization terms:

$$Obj(\vec{\boldsymbol{\beta}}) = \left\|Y - \hat{Y}\right\|_2 + \alpha||\vec{\boldsymbol{\beta}}||_0 + \gamma||\vec{\boldsymbol{\beta}}||_1$$

where $||\vec{\boldsymbol{\beta}}||_1$ is the L1-norm defined as $||\vec{\boldsymbol{\beta}}||_1 = \sum_{i=1}^{n} |\beta_i|$, and $||\vec{\boldsymbol{\beta}}||_0$ is the L0-norm defined as $||\vec{\boldsymbol{\beta}}||_0 =$ number of non-zero $\beta_i$, when $\vec{\boldsymbol{\beta}} = \{\beta_1, \beta_2, \dots, \beta_n\}$.

Solving L0 regularized regression is extremely challenging since it is not a convex problem. PoolHapX takes advantage of the L0Learn package, a recent development in the field of optimization(Hazimeh and Mazumder 2018), to solve this regression problem.

By applying the above regression to all hosts, the cross-host (population) frequencies of each haplotype can be determined by combining the within-host frequencies.

#### SECTION III: SIMULATION PROCEDURES

##### Simulating Haplotypes for Each Pool Using SLiM

The overall procedure for the simulations is shown in **Supplementary Figure 8**

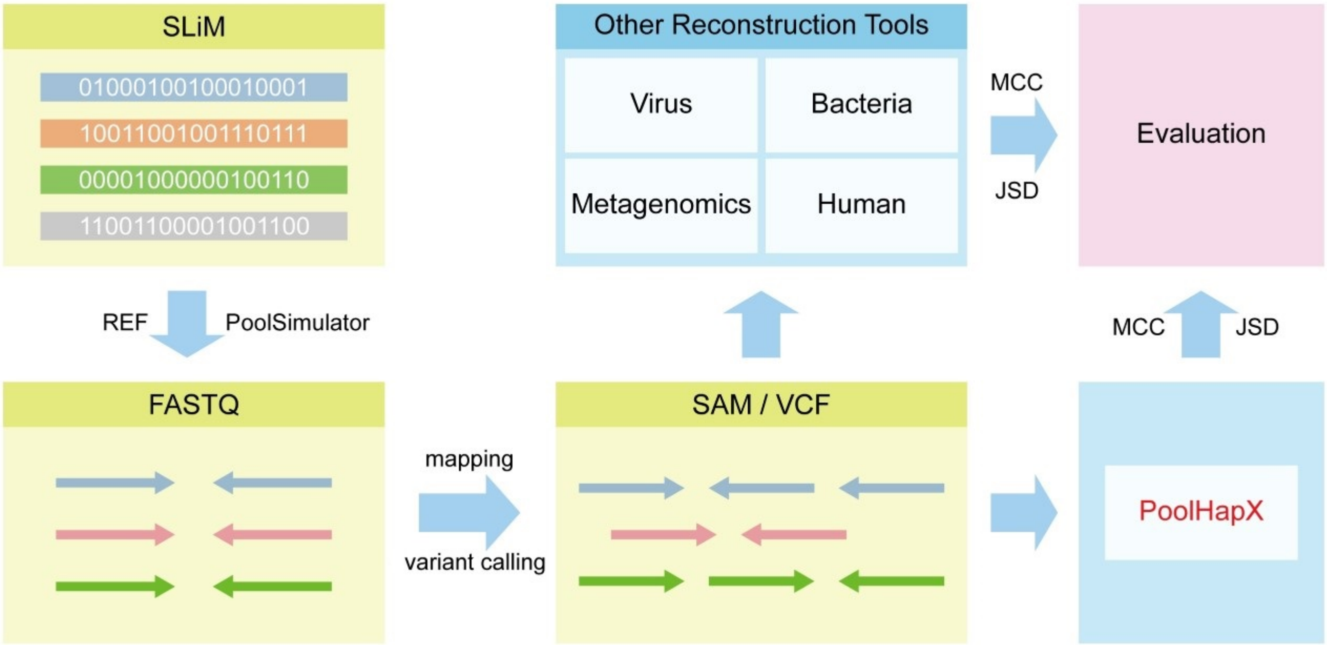


**Supplementary Fig. 8: Flowchart of the testing pipeline.** First, the forward genetic simulation framework SLiM(Haller and Messer 2019) is used to generate haplotypes for each pool. These haplotypes are collected by a Java program called PoolSimulator (https://github.com/theLongLab/PoolHapX/tree/master/Simulation_And_Analysis/PoolSimulator_SLiM.jar), which embeds the variant sites into the real reference genome and calls genome sequencing simulators (such as DWGSIM and for short reads and LRSIM for 10X Genomics barcoded linked-reads) to simulate sequencing reads. BWA(Li and Durbin 2009), Samtools(Li, et al. 2009), and tools from the GATK suite(DePristo, et al. 2011) are used to process read alignment, reference mapping, and variant calling on the FASTQ files produced by PoolSimulator. PoolHapX takes the processed SAM and CVF files as input. The reconstructed haplotypes from PoolHapX are evaluated by comparing them against the “gold standard” haplotypes generated by SLiM. Although the simulation procedures will vary for different species, the evaluation criteria are the same for all species. We compute the Jensen–Shannon divergence (JSD) and the Matthews correlation coefficient (MCC) between reconstructed haplotypes and the gold standard haplotypes in order to measure the accuracy PoolHapX against existing haplotypes reconstruction tools.

We use SLiM (version 3)(Haller and Messer 2019), a forward genetic simulation framework, to generate haplotypes for each pool. SLiM-3 provides an R-like scripting language called “Eidos” for users to customize their simulations. By default, SLiM simulates diploid individuals whereas most viruses and bacteria, including the ones we used in simulations, are haploid. Therefore, we configure SLiM to use a non-Wright-Fisher model (see the SLiM manual)(Haller and Messer 2016) to simulate haploid individuals, which is recommended by the authors of SLiM. To mimic cross-pool gene flow, we chose an island model with a constant migration rate of 5%. All simulations generated haplotypes for either 25 or 50 pools, and we set the carrying capacity for each pool as 70, which is lower than the actual number of individuals in a normal host. However, in practice we found that for simulations with carrying capacities of 100 for 50 pools with positive mutations, SLiM took over 5 days and ran out of memory on our computing cluster. To compensate for a smaller number of individuals per pool, we set a higher mutation rate so that the population mutation rate, which is the product of effective population size and the per-base mutation rate, remains reasonable.

The mutation rates are chosen differently depending on the specific application, chromosome length, and the density of single nucleotide polymorphisms (SNPs). For neutral simulations without evolutionary selection, we use only one type of mutation without any changes to fitness. We also simulate haplotypes under different evolutionary scenarios with selection, where we add another type of mutation with negative or positive changes to fitness. Following the default settings of SLiM, all mutations that became fixed (i.e., frequency of the mutations reaching 100% in the population) are removed from the population of the genomes. After simulating a set number of generations, we save mutations with a minor allele frequency (MAF) above 0.05 into SLiM's standard output format, which contains information on each mutation present and the haplotypes in each pool. Although SLiM supports the generation of real genomic bases (i.e., A, C, G, and T), simulations using reference genomes are much slower. Therefore, we specify the length of the genome and embed the resulting polymorphic sites into the reference backbone to produce an equivalent output.

*Viral haplotype simulation*

We use the HIV strain HXB2 (https://www.ncbi.nlm.nih.gov/nuccore/K03455.1?report=fasta) as the reference genome. The length of this genome is 9719 base pairs (bp). We set the mutation rate as $5\times10^{-7}$, $8\times10^{-7}$, and $2\times10^{-6}$ to simulate 50, 100, and 200 loci, respectively. We run the simulation for 10,000 generations to generate the SLiM standard output.

*Bacterial haplotype simulation*

We use chromosome 2 of Vibrio cholerae O1 biovar El Tor str. N16961 as the reference genome (https://www.ncbi.nlm.nih.gov/genome/505?genome_assembly_id=300197). The length of this chromosome is 1.07 Mb. We set the mutation rate as $1.7\times10^{-9}$, $4\times10^{-9}$, and $9\times10^{-9}$ for simulating 50, 100, and 200 variant loci, respectively. We run the simulation for 20,000 generations to generate the SLiM standard output.

*Metagenomic haplotype simulation*

We compare the performance of PoolHapX against two metagenomic haplotype reconstruction tools: StrainEst(Albanese and Donati 2017) and Gretel(Nicholls, et al. 2019). StrainEst is a reference-based method that uses the single nucleotide variant (SNV) profiles of available genomes from selected species to determine the number and identity of coexisting strains. In reality, the true number of haplotypes is generally unknown. To ensure that StrainEst is able to run, we use the SNV profiles of *E. coli* genomes which are available in the StrainEst distribution (ftp://ftp.fmach.it/metagenomics/strainest/ref) as our reference genome.

Gretel does not require a priori knowledge of the input data, such as references of many known strains or the true number of haplotypes), and is therefore directly comparable to PoolHapX. However, Gretel will fail to run if any pair of genetic variants is not covered by the same sequencing read. Therefore, using the reference genome provided by StrainEst, we simulate a genomic region with a very high variant density. In particular, we set the mutation rate as $8\times10^{-7}$, $10^{-6}$, and $2\times10^{-6}$ for simulating 50, 100, and 200 loci, respectively. The length of the simulated region is 4972 bp from position 68525 (68525 is the start point in the StrainEst

SNV profile) to 73496. We run the simulation for 6,000 generations, to generate the SLiM standard output.

*Haplotype simulation under evolutionary scenarios*

SLiM provides a mechanism of specifying multiple types of mutations in the same simulation. In the following evolutionary scenarios, we keep most mutations neutral but add some non-neutral mutations to allow selection to occur. We use HIV strain HXB2 (https://www.ncbi.nlm.nih.gov/nuccore/K03455.1?report=fasta) as the reference genome, and simulate haplotypes under three commonly studied evolutionary scenarios: positive selection, negative selection, and selective sweeps. The proportion of neutral mutations is set to 0.7 for positive and negative selections. For selective sweep, we simulate only one strongly favorable mutation and keep the remaining mutations neural.

Please note that, in SLiM simulations, we cannot absolutely control the number of loci in the final results especially when multiple evolutionary constrains are present, so we managed to adjust evolutionary parameters a bit to make the number of loci roughly 50, 100, and 200 in respective simulations.

For the positive selection model, we combined neutral mutations and mutations with a fixed positive fitness change of 0.05, as specified by a SLiM parameter for the strength of positive selection. The mutation rate is $3\times10^{-6}$, $8\times10^{-6}$, and $5\times10^{-5}$ for simulating roughly 50, 100, and 200 loci, respectively. For the negative selection model, we combine neutral mutations and mutations with a fixed negative fitness change of -0.05. The mutation rate is $8\times10^{-7}$, $1\times10^{-6}$, and $3\times10^{-6}$ for simulating 50, 100, and 200 loci, respectively. For the selective sweep model, we combine neutral mutations and mutations with a large fixed positive fitness change of 1.0 at the 100th generation. The mutation rate is $6\times10^{-7}$, $1\times10^{-6}$, and $2\times10^{-6}$ for simulating roughly 50, 100, and 200 loci, respectively. We run the simulation for 10,000 generations to generate the SLiM standard output.

*Haplotype simulation when provided with single-cell linked-reads*

We use *Chlamydia trachomatis* 434/Bu (https://www.ncbi.nlm.nih.gov/genome/471) as the reference genome. The length of this genome is 1,038,841 bp. We set the mutation rate as $9\times10^{-9}$ and run for 50,000 generations to generate 570 mutations and 31 global haplotypes.

*Example of a SLiM script*

The following example is a SLiM script used to simulate haplotypes of HIV strain HXB2(Ratner, et al. 1985) under neutral mutation for 25 pools. For all simulation scripts and detailed explanations of each command, please refer to our GitHub (https://github.com/theLongLab/PoolHapX/tree/master/Simulation_And_Analysis/Simulation_SLiM_scripts):

```
// Based on the SLiM v3
// adjustable parameters: migration_rate; fitness coefficients; maf_cutoff; number of pools

initialize() {
	initializeSLiMModelType("nonWF");
	defineConstant("K", 70);	// carrying capacity
	defineConstant("migration_rate", 0.05); // migration rate
	chromosome_len=9719;
	defineConstant("L", chromosome_len); // chromosome length
```

```
	defineConstant("H", 0.001); // Recombination probability
	initializeMutationRate(8e-7);
	initializeMutationType("m1", 1.0, "f", 0.0);
	initializeMutationType("m3", 1.0, "f", -0.03);  // deleterious
	initializeMutationType("m4", 1.0, "f", 0.05);      // beneficial
	initializeGenomicElementType("g1", c(m4,m3,m1), c(0.0, 0.0, 5.0)); // m4 or m3 not used
	initializeGenomicElement(g1, 0, chromosome_len-1);
	initializeRecombinationRate(0);   // no regular recombination in diploid sense. Instead, 
the recombination is modelled by HGT below.
}
reproduction() {
	for(p_i in sim.subpopulations){
			if (runif(1) < H)
		{
			// horizontal gene transfer from a randomly chosen individual
			HGTsource = p_i.sampleIndividuals(1).genome1;
			// draw two distinct locations; redraw if we get a duplicate
			the_break = rdunif(1, max=L-1);
			recomb = c(0, the_break, L-1);
			subpop.addRecombinant(genome1, HGTsource, recomb, NULL, NULL,      NULL);
		}
		else
		{
			// no horizontal gene transfer; clonal replication
			subpop.addRecombinant(genome1, NULL, NULL, NULL, NULL, NULL);
		}
	}

}
1 {
	metapopSide = 5;       // number of subpops along one side of the grid
	metapopSize = metapopSide * metapopSide; //
	for (i in 1:metapopSize)
		sim.addSubpop(i, 70);
}
early(){
	// random migration
	nIndividuals = sum(sim.subpopulations.individualCount);
	nMigrants = rpois(1, nIndividuals * migration_rate);
	migrants = sample(sim.subpopulations.individuals, nMigrants);
	for (migrant in migrants){
		do dest = sample(sim.subpopulations, 1);
		while (dest == migrant.subpopulation);
		dest.takeMigrants(migrant);
	}
	for (subpop in sim.subpopulations){
```

```
            subpop.fitnessScaling = K / subpop.individualCount;
        }
}
late() {
        // remove mutations in the haploid genomes that have fixed
        muts = sim.mutationsOfType(m1);
        freqs = sim.mutationFrequencies(NULL, muts);
        if (any(freqs >= 0.5))
            sim.subpopulations.genomes.removeMutations(muts[freqs >= 0.5], T);
        muts = sim.mutationsOfType(m3);
        freqs = sim.mutationFrequencies(NULL, muts);
        if (any(freqs >= 0.5))
            sim.subpopulations.genomes.removeMutations(muts[freqs >= 0.5], T);
        muts = sim.mutationsOfType(m4);
        freqs = sim.mutationFrequencies(NULL, muts);
        if (any(freqs >= 0.5))
            sim.subpopulations.genomes.removeMutations(muts[freqs >= 0.5], T);
}

10000 late() {
        outSize=p1.individualCount * 2;
        maf_cufoff=0.1;
        muts4=sim.mutationsOfType(m4);
        freqs4 = sim.mutationFrequencies(NULL, muts4);
        remove4 = muts4[freqs4 < maf_cufoff/2];
        muts1=sim.mutationsOfType(m1);
        freqs1 = sim.mutationFrequencies(NULL, muts1);
        remove1 = muts1[freqs1 < maf_cufoff/2];
        muts3=sim.mutationsOfType(m3);
        freqs3 = sim.mutationFrequencies(NULL, muts3);
        remove3 = muts3[freqs3 < maf_cufoff/2];
        removes=c(remove4, remove3, remove1);
        sim.subpopulations.genomes.removeMutations(removes, T);
        sim.outputFull();
}
```

The details including number of haplotypes, variants, average allele frequency and average pair-wise difference of the haplotypes in all the simulations are listed in the two tables below.

**Supplementary Table 1:** The number of haplotypes, number of loci, average alternative allele frequencies and average pairwise differences for our simulations (50 pools).

| Project | Hap_Count | Num_Var | Ave_Alt_Freq | Ave_Pw_Diff |
|---|---|---|---|---|
| | 41 | 43 | 14.98 | 15.91 |
| **Virus** | 73 | 102 | 27.55 | 32.79 |
| | 42 | 179 | 70.14 | 44.06 |
| | 42 | 52 | 16.07 | 20.24 |

| | | | | |
|---|---|---|---|---|
| **Bacteria** | 43 | 107 | 40.12 | 37.16 |
| | 43 | 199 | 65.27 | 68.48 |
| | 39 | 56 | 16.17 | 21.38 |
| **Metagenomics** | 37 | 99 | 33.13 | 39.35 |
| | 36 | 195 | 57.69 | 69.72 |
| | 41 | 43 | 14.97 | 15.91 |
| **Positive Selection** | 50 | 98 | 35.51 | 15.11 |
| | 104 | 193 | 53.94 | 51.98 |
| | 36 | 52 | 15.38 | 17.82 |
| **Negative Selection** | 36 | 109 | 34.75 | 41.86 |
| | 45 | 199 | 74.98 | 79.24 |
| | 23 | 48 | 15.87 | 17.18 |
| **Selective Sweep** | 36 | 111 | 36.19 | 37.56 |
| | 27 | 195 | 58.52 | 75.32 |
| **10X-linked Reads** | 40 | 570 | 199.55 | 223.69 |

**Supplementary Table 2:** The number of haplotypes, number of loci, average alternative allele frequencies and average pairwise differences for our simulations (25 pools).

| Project | Hap_Count | Num_Var | Ave_Alt_Freq | Ave_Pw_Diff |
|---|---|---|---|---|
| | 32 | 43 | 14.62 | 15.83 |
| **Virus** | 54 | 102 | 27.78 | 33.78 |
| | 33 | 180 | 69.27 | 46.34 |
| | 36 | 52 | 15.75 | 20.33 |
| **Bacteria** | 31 | 107 | 40.12 | 37.16 |
| | 36 | 199 | 65.30 | 69.66 |
| | 30 | 56 | 16.13 | 21.53 |
| **Metagenomics** | 30 | 99 | 33.00 | 39.36 |
| | 31 | 195 | 57.83 | 69.81 |
| | 32 | 43 | 14.62 | 15.83 |
| **Positive Selection** | 40 | 98 | 25.45 | 35.73 |
| | 77 | 193 | 52.62 | 50.91 |
| | 31 | 52 | 15.22 | 17.60 |
| **Negative Selection** | 31 | 109 | 34.74 | 42.10 |
| | 42 | 199 | 77.22 | 45.77 |
| | 21 | 48 | 15.90 | 17.53 |
| **Selective Sweep** | 30 | 111 | 36.57 | 38.56 |
| | 24 | 195 | 58.71 | 75.89 |
| **10X-linked Reads** | 31 | 570 | 199.29 | 219.41 |

#### Human Haplotype Preparation

As sequences of many individuals are available online, we use the phased haplotypes from the 1000 Genomes Project (https://www.internationalgenome.org/) instead of simulating haplotypes. We reconstructed haplotypes from 26 different populations from the locations listed in the following table:

| Population Code | Population Description |
|---|---|
| CHB | Han Chinese in Beijing, China |
| JPT | Japanese in Tokyo, Japan |
| CHS | Southern Han Chinese |
| CDX | Chinese Dai in Xishuangbanna, China |
| KHV | Kinh in Ho Chi Minh City, Vietnam |
| CEU | Utah Residents (CEPH) with Northern and Western European Ancestry |
| TSI | Toscani in Italia |
| FIN | Finnish in Finland |
| GBR | British in England and Scotland |
| IBS | Iberian Population in Spain |
| YRI | Yoruba in Ibadan, Nigeria |
| LWK | Luhya in Webuye, Kenya |
| GWD | Gambian in Western Divisions in the Gambia |
| MSL | Mende in Sierra Leone |
| ESN | Esan in Nigeria |
| ASW | Americans of African Ancestry in SW USA |
| ACB | African Caribbeans in Barbados AFR |
| MXL | Mexican Ancestry from Los Angeles USA |
| PUR | Puerto Ricans from Puerto Rico |
| CLM | Colombians from Medellin, Colombia |
| PEL | Peruvians from Lima, Peru |
| GIH | Gujarati Indian from Houston, Texas |
| PJL | Punjabi from Lahore, Pakistan |
| BEB | Bengali from Bangladesh |
| STU | Sri Lankan Tamil from the UK |
| ITU | Indian Telugu from the UK |

We reconstruct haplotypes in the region of the gene ZNF737 and its flanking region of 5Mb in chromosome 19. This region contains on average 200 SNPs and 34 global haplotypes cross multiple populations. The gold standard for human haplotypes is directly drawn from the original data file, which is available at

http://ftp.1000genomes.ebi.ac.uk/vol1/ftp/data_collections/1000_genomes_project/release/20190312_biallelic_SNV_and_INDEL/.

#### SECTION IV: ANALYTIC PROCEDURE

##### Processing SLiM Output

The SLiM standard output contains information for all simulated mutations and individual haplotypes within each pool. We developed a Java program called PoolSimulator to process the SLiM output. This program collects haplotypes from each pool, embeds their variant sites into the real reference genome, and calls another tool (DWGSIM(Homer 2010) for short reads or LRSIM(Luo, et al. 2017) for linked-reads to simulate sequencing reads. The source code and executable is available is downloadable from our GitHub (https://github.com/theLongLab/PoolHapX).

PoolSimulator first parses the SLiM standard output to generate global haplotypes and their corresponding frequencies within each pool. Since SLiM encodes mutated alleles as a string of 0s and 1s, a reference genome is then used to generate full sequences from the polymorphic sites generated by SLiM. The true cross-pool and within-pool haplotype distributions are written in separate files to be used as the gold standard for evaluating PoolHapX's performance. Once we have a mixture of haplotypes for each pool, a genome-sequencing simulator is used to generate sequencing reads. We use DWGSIM(Homer 2010) (https://github.com/nh13/DWGSIM) to generate paired-end reads, and LRSIM(Luo, et al. 2017), a 10x Genomics Reads Simulator (https://github.com/aquaskyline/LRSIM), to generate single-cell linked-reads. The average read length is 150 with an outer distance of 400, and an error rate of 0.001 errors per base. Reads are simulated using different combinations of coverage (50, 100, 250 or 1000) depending on the simulation type. The simulated reads are saved as FASTQ files in the input directory for further processing.

We use property files to more easily run multiple simulations and store simulation parameters for future reference. In the file we record parameters including the locations of input files, intermediate files, gold standard files, and SLiM output. We run PoolSimulator with the following command:

> *java –Xmx4G -jar PoolSImulator.jar PoolSimulator.properties*

The following example is a PoolSimulator.properties file for processing viral haplotypes simulated by SLiM from the HIV HXB2 reference genome. All property files are available in our GitHub repository (https://github.com/theLongLab/PoolHapX/tree/master/Simulation_And_Analysis)

```
# Project name
Proj_Name = project_name
# Path to difference directories
Input_Dir = /path/to/input
Intermediate_Dir = /path/to/intermediate
Gold-Standard_Dir = /path/to/gold_standard
Slim_Output_Path = /path/to/slim_out
# Path to reference genome and its genome length
Reference_Seq= /path/to/HIV_HXB2.fa
Ref_Seq_Len = 9719
# Path to DWGSIM
DWGSIM = /path/to/dwgsim
# Number of pools
Num_Pools = 25
```

```
# Error rate per base
Error_Rate_Per_Base = 0.001
# Haplotype frequency cutoff
Hap_Freq_Cutoff = 0.0
# Coverage
Coverage = 5000
# Read length
Read_Len = 150
# Outer distance (i.e., the distance of the two farthest points of the two reads)
Outer_Dist = 400
Weak_Length = 550
# Is_Perfect = true will generated VEF file for running PoolHapX
Is_Perfect = false
# Using island model
Slim_Model = island_haploid
# For island model, set below two parameters as false
Is_Single_Population = false
Is_Ms_Output = false
# This is to control number of individual haplotypes per pool
Num_Haps_Pool = 30
```

##### Sequence Alignment and Variant Calling

We use BWA, Samtools, and tools from the GATK suite to process read alignment, reference mapping, and variant calling on the FASTQ files produced by PoolSimulator. The allele-annotated reads generated by this pipeline may contain false positive segregating sites and false negative (missed) sites due to errors in pre-processing, such as read or variant dropout due to poor mapping or base quality. This mirrors the imperfect data processing in practise and challenges PoolHapX's ability to robustly handle data with errors.

Our sequence alignment and variant calling workflow is as follows:

For each pool, use 'gunzip' to extract the "fastq.gz" files into two paired-end FASTQ files. The placeholder "{prefix}" has been substituted for the name of the actual file to be processed.

```
gunzip /path/to/{prefix}.bwa.read1.fastq.gz
gunzip /path/to/{prefix}.bwa.read2.fastq.gz
```

Using BWA-MEM, map the simulated reads to the reference genome (name replaced by placeholder "{ref}"), and then generate an output in the BAM binary format.

```
> bwa mem
      -R /path/to/{ref}
     /path/to/{prefix}.bwa.read1.fastq
     /path/to/{prefix}.bwa.read2.fastq|samtools view -Shub →/path/to/{prefix}.bam
```

Using Samtools-sort, sort the BAM file by coordinate and create an output file with the suffix "srt.bam".

```
> samtools sort
```

```
        -I /path/to/{prefix}.bam
        -o /path/to/{prefix}.srt.bam
```

For each pool using GATK-AddOrReplaceReadGroups, assign all reads in a file to a single new “read group”. The pool ID is given as “{task_ID}” and the pool name is given as “{pool_name}”.

```
> gatk AddOrReplaceReadGroups
        -I /path/to/{prefix}.srt.bam
        -O /path/to/{prefix}.rg.bam
        -ID {task_ID}
        -LB NPD -PL Illumina -PU NPD
        -SM {pool_name}
```

For each pool using Samtools-index, generate an index file for each sorted BAM file. Then, use GATK-HaplotypeCaller to call variants into a raw gVCF file.

```
> samtools index  /path/to/{prefix}.rg.bam

> gatk
        --java-options "-Xmx20g -XX:+UseConcMarkSweepGC -XX:ParallelGCThreads=4"
        HaplotypeCaller
        -R /path/to/{ref}
        -I /path/to/{prefix}.rg.bam
        -ERC GVCF
        -ploidy 8
        --heterozygosity 0.01
        --max-alternate-alleles 1
        -O /path/to/{prefix}.raw.g.vcf
```

Using GATK-CombineGVCFs, join all pool-specific raw gVCF files into a joint gVCF file. Then, using GATK-GenotypeGVCFs, merge the gVCF records for variant discovery into a raw VCF file. Finally, use GATK-SelectVariants to select subsets of variants from the raw VCF file.

```
> gatk CombineGVCFs
        -R /path/to/{ref}
        {ALL_Pools}.raw.g.vcf
        -O /path/to/{prefix}.g.vcf
>gatk GenotypeGVCFs
        -R /path/to/{ref}
        -V /path/to/{prefix}.g.vcf
        -ploidy 8
        -O /path/to/{prefix}.raw.vcf
> gatk SelectVariants
        -R /path/to/{ref}
        -V /path/to/{prefix}.raw.vcf
        -O /path/to/{prefix}.vcf
```

Convert each pool-specific sorted BAM file into a sorted SAM file.

```
> samtools view
```

*-ho /path/to/{prefix}.sam*
*/path/to/{prefix}.srt.bam*

#### PoolHapX Analysis

*Filtering of genetic variants*

PoolHapX operates on BAM files, which record the read alignment status, and VCF files, which record the discovered genetic variants. Note that non-reference bases may be discovered in the read sequences of the BAM files, but such bases are not included in the haplotype reconstruction process if they are not present in the VCF files. Uncalled non-reference bases are likely to be sequencing or alignment errors. When comparing reconstructed haplotypes to gold standard, only correct variants are used. As PoolHapX does not intend to analyze very rare global haplotypes, we remove all sites with MAF lower than 0.05 from the VCF files.

The parameters used to run PoolHapX are recorded in property files, which are available in our GitHub repository (https://github.com/theLongLab/PoolHapX/tree/master/Simulation_And_Analysis). We use default parameters in most cases, with the only exception being the analysis of *P. vivax* data for which we have used stronger regularization.

*Comparison criteria*

*MCC+JSD:* Although the simulation procedure varies across different species, the assessment criteria are the same for all species. For each haplotype reconstruction tool, we compute the Jensen–Shannon divergence (JSD)(Fuglede and Topsoe 2004) and the Matthews correlation coefficient (MCC)(Powers 2011) between the simulated (gold standard) and reconstructed haplotypes to measure the reconstruction accuracy. Each reconstructed haplotype is first mapped to its nearest gold standard haplotype using Hamming distance(Tangherloni, et al. 2019). Both haplotypes are represented by a string composed of 1s and 0s (representing reference and alternate alleles). MCC is calculated between such pairs and then averaged across all haplotypes in the pools.

The Matthews correlation coefficient (MCC) is used to measure the similarity of the simulated and reconstructed haplotype. The MCC is defined as

$$MCC = \frac{TP \times TN - FP \times FN}{\sqrt{(TP + FP)(TP + FN)(TN + FP)(TN + FN)}}$$

where TP is the number of true positives, TN the number of true negatives, FP the number of false positives, and FN the number of false negatives. The MCC returns a value in the interval [−1,1] where 1 represents an exact prediction, 0 represents a random prediction, and –1 indicates total disagreement between the predicted and actual haplotypes.

The Jensen–Shannon divergence (JSD) evaluates the divergence between the frequencies of simulated and reconstructed haplotype distributions. The probability distributions A and P represent the frequency distributions of haplotype in the pool, where A is the distribution of gold standard haplotypes, and P is the reconstructed haplotype distribution. The JSD is defined as

$$JSD(A \parallel \mathrm{P}) = \frac{1}{2}\mathrm{D}(A \parallel \mathrm{K}) + \frac{1}{2}\mathrm{D}(P \parallel \mathrm{K})$$

where

$$K = \frac{1}{2}\mathrm{D}(A + \mathrm{P}).$$

*D*(*A*||*P*) is the Kullback–Leibler divergence from A to P, which is defined as

$$\mathrm{D}(A \parallel \mathrm{P}) = \sum_{i} A(i) \log_2 \frac{A(i)}{P(i)}.$$

The JSD is symmetric for the comparing distributions (i.e., *A* and *P*), and for two probability distributions has a value in the interval [0,1] (Fuglede and Topsoe 2004), where 0 represents an exact prediction.

*F1-Score* + *CRR* + *FDR*: Here, the same as the MCC/JSD case, only the reconstructed haplotypes with frequency higher than 0.01 are counted in the comparison. As the targeted application is to reconstruct many haplotypes among many pools, we do not require the true positives to be defined as exactly perfect match. Instead, among these reconstructed haplotypes with frequency higher than 0.01, the ones that are close enough to a gold-standard haplotype (MCC >0.8) is considered as true positives, and the rest are considered as false positives. If there is no reconstructed haplotype with frequency higher than 0.01 can match a gold-standard haplotype, then this gold-standard haplotype will be deemed as a false negative.

With the above definition, we can define the following metrics:

precision = true_positives / (true_positives + false_positives)

recall = true_positives / (true_positives + false_negatives)

F1-Score = 2 * (precision * recall) / (precision + recall)

Additionally, the CRR (correctly reconstructed rate) is defined as the number of correctly reconstructed haplotypes divided by the total number of the gold-standard haplotypes. FDR (false discovery rate) is defined by the number of incorrectly reconstructed haplotypes divided by the total number of the haplotypes detected by the corresponding tool.

#### SECTION V: RUNNING OTHER HAPLOTYPE RECONSTRUCTION TOOLS

To evaluate the performance of PoolHapX, we compared its performance against state-of-the-art haplotype reconstruction tools in four fields: viral, bacterial, metagenomic, and human genetics. Every haplotype reconstruction tool is benchmarked with the same input data provided to PoolHapX. We describe our procedure for running these tools below.

##### Viral Quasispecies Reconstruction Tools

We compared PoolHapX with three viral quasispecies reconstruction tools: CliqueSNV(Knyazev, et al. 2018), PredictHaplo(Prabhakaran, et al. 2013), and TenSOR(Ahn, et al. 2018).

###### *CliqueSNV*

CliqueSNV is a reference-based method for reconstructing viral variants from next generation sequencing (NGS) short reads, which constructs an allele graph based on linkage between single nucleotide variations and identifies true viral variants by merging cliques of that graph via combinatorial optimization techniques. The open source implementation of CliqueSNV is available at (https://github.com/vyacheslav-tsivina/CliqueSNV). We run CliqueSNV using the default setting listed under the "Usage example" section, which is the following command:

### Run CliqueSNV software to reconstruct haplotypes

```
> java -Xmx20g -jar path/to/clique-snv.jar
        -m snv-illumina
        -in path/to/{prefix}.rg.bam
```

The parameters are interpreted below:
-Xmx. Controls the maximum memory used by the Java runtime environment (JVM).
path/to/clique-snv.jar. Indicates full path to the CliqueSNV jar file.
-m snv-illumina. Sets CliqueSNV to run on Illumina data-in. Indicates alignment file to use.

###### *PredictHaplo*

PredictHaplo implements a fully probabilistic approach to quasispecies reconstruction. Given a set of aligned reads, PredictHaplo uses a Bayesian mixture model with a Dirichlet process prior to adaptively estimate the unknown number of underlying haplotypes(Prabhakaran, et al. 2013). Inference is carried out via a Markov chain Monte Carlo sampling scheme, which is optimized to scale with real-world NGS data sets. We perform global haplotype reconstruction over long genomic regions using the PredictHaplo software available at (https://bmda.dmi.unibas.ch/software.html), with the following commands:

### Using BWA-MEM, map the simulated reads to the reference sequence "{ref}", and generate an output "{input_sam}" in the SAM format:

```
> bwa mem    -R /path/to/{ref}
        /path/to/{prefix}.bwa.read1.fastq
        /path/to/{prefix}.bwa.read2.fastq
        > /path/to/{input_sam}
```

Then we run PredictHaplo to reconstruct haplotypes:
*>/path/to/PredictHaplo_Paired {config_file}*
Parameters:
/path/to/PredictHaplo_Paired. Indicates full path to the PredictHaplo_Paired executable file. As our simulation produces paired-end reads, we use PredictHaplo_Paired instead of PredictHaplo.
{config_file}. Indicates the input configuration file.

To run PredictHaplo, we have downloaded their example configuration file, which can be downloaded using the link https://bmda.dmi.unibas.ch/software.html . Inside the folder, there is a file named as "config_5V" for us to start. However, the default parameters are not runnable on our side. So we have adjusted two parameters: First, we have lower down the '% min_readlength' to from its default of 220 to 100, because our simulated read length is 150. Second, following a published paper comparing tools(Schirmer, et al. 2014), we have We set the parameter '% entropy_threshold' to 5e-3. Then the software is runnable on our side.

The following is an example of our configuration file for PredictHaplo:

```
% configuration file for the PredictHaplo
% prefix  # The name for the file
{predix}
% filename of reference sequence (FASTA) #Indicates the full path to the reference file.
/path/to/{ref}
% do_visualize (1 = true, 0 = false)
0
% filname of the aligned reads (sam format) # Indicates the full path to the input sam
file.
/path/to/{input_sam}
% have_true_haplotypes  (1 = true, 0 = false)
0
% filname of the true haplotypes (MSA in FASTA format) fill in any dummy filename if
there is no true haplotypes)
null
% do_local_analysis  (1 = true, 0 = false) (must be 1 in the first run)
1
% max_reads_in_window
10000
% entropy_threshold
5e-3
%reconstruction_start
0
%reconstruction_stop
9719
%min_mapping_qual
30
%min_readlength
```

```
100
%max_gap_fraction (relative to alignment length)
0.05
%min_align_score_fraction (relative to read length)
0.35
%alpha_MN_local (prior parameter for multinomial tables over the nucleotides)
25
%min_overlap_factor (reads must have an overlap with the local reconstruction window of at least this factor times the window size)
0.85
%local_window_size_factor (size of  local reconstruction window relative to the median of the read lengths)
0.7
% max number of clusters (in the truncated Dirichlet process)
25
% MCMC iterations
501
% include deletions (0 = no, 1 = yes)
0
```

*TenSQR*

TenSQR utilizes a tensor factorization framework to analyze high-throughput sequencing data and reconstruct viral quasispecies characterized by highly uneven frequencies of their components(Ahn, et al. 2018). TenSQR performs clustering with successive data removal to successively infer strains in a quasispecies from most to least abundant. As each strain is inferred, the sequencing reads originating from that strain are removed from the dataset. The open source implementation of TenSQR is available at (https://github.com/SoYeonA/TenSQR).

To run this tool with its default settings, we have downloaded a default config provided by the tool website (https://github.com/SoYeonA/TenSQR/blob/master/config ). We only edited the reconstruction start and stop positions, and kept all the rest parameters default.

We run TenSQR with the following commands:

Using BWA-MEM, map the simulated reads to the reference sequence "{ref}", and generate an output "{input_sam}" in the SAM format:

```
> bwa mem
    -R /path/to/{ref}
    /path/to/{prefix}.bwa.read1.fastq
    /path/to/{prefix}.bwa.read2.fastq >  /path/to/{input_sam}
```

Run TenSQR to reconstruct haplotypes:

```
/path/to/ExtractMatrix {config_file}
Python /path/to/TenSQR.py {config_file}
```

The parameters are:

/path/to/ExtractMatrix. Indicates full path to the executable file ExtractMatrix.
/path/to/TenSQR.py. Indicates full path to the executable file TenSQR.py.
{config_file}. Indicates the input configuration file.

Here is an example of the configuration file that we used:

```
% configuration file for TenSQR
filename of reference sequence (FASTA):
/path/to/{ref}
filname of the aligned reads (sam format) :
/path/to/{input_sam}
SNV_thres : 0.01
reconstruction_start : 1
reconstruction_stop: 9719
min_mapping_qual : 40
min_read_length : 100
max_insert_length : 400
characteristic zone name : {predix}
seq_err (assumed sequencing error
rate(%)) : 0.1
MEC improvement threshold : 0.0312
initial population size :100
```

#### Bacterial Haplotype Reconstruction Tools

We compared PoolHapX with two bacterial haplotype reconstruction tools: EVORhA(Pulido-Tamayo, et al. 2015) and Bhap(Li, et al. 2019).

##### *EVORhA*

EVORhA is a haplotype reconstruction method that combines phasing information in regions of overlapping reads with the estimated frequencies of inferred local haplotypes. EVORhA first reconstructs haplotypes at the local level, and then links local reconstructions into extended haplotypes using information from overlapping reads. The implementation of EVORhA can be downloaded from http://bioinformatics.intec.ugent.be/kmarchal/EVORhA/.

We run EVORhA with the following commands, which are their default settings:

```
>java -jar /path/to/evorha.jar
        completeAnalysis /path/to/{ref}
        /path/to/{input_bam}
```

The parameters are quite simple: reference genome and input BAM file.

##### *Bhap*

BHap is a tool for reconstructing bacterial haplotypes from next generation sequencing (NGS) data, and is based on fuzzy flow networks. The implementation of BHap can be downloaded from http://hulab.ucf.edu/research/projects/BHap/BHap.html.

We run BHap with the following commands, which are their default settings:

```
>python run_BHap.py
        -d /path/to/{prefix}.bhap_output
        -r $ref
        -t fastq -1 /path/to/{prefix}.bwa.read1.fastq -2 /path/to/{prefix}.bwa.read2.fastq
        -l 150 -c 100 -g $ref.length
```

Parameters:
/path/to/{prefix}.bhap indicates the full path to the BHap executable file
ref indicates the full path to the reference genome
-l, sets read length to 150
-c, sets coverage to 100

##### Metagenomics Haplotypes Reconstruction Tools

We compared PoolHapX with two metagenomic haplotype reconstruction tools: StrainEst(Albanese and Donati 2017) and Gretel(Nicholls, et al. 2019).

###### *StrainEst*

StrainEst is a reference-based method that uses genomic data from selected species to reconstruct complex strain profiles from metagenomic sequencing. This tool can quantify the number and identity of coexisting strains and their relative abundances in mixed metagenomic samples. The source code is available at https://github.com/compmetagen/strainest.

We run StrainEst with the following commands, which are their default settings:

```
>/path/to/strainest est
        E_coli/snp_clust.dgrp
        /path/to/{input_bam}
        output_dir
```

/path/to/strainest est. Full path to executable file strainest est
snp_clust.dgrp. This file corresponds to the genetic variants (SNP) profiles of the available genomes of E.coli strains, available at ftp://ftp.fmach.it/metagenomics/strainest/ref
/path/to/{input_bam}. Full path to sorted BAM files
output_dir. Full path to the output directory

###### *Gretel*

Gretel is able to recover haplotypes from metagenomic data. Gretel parses an alignment of reads into a Hansel matrix and uses SNP pairs observed in contiguous reads to probabilistically reconstruct the most likely haplotypes. Gretel uses an L-th order Markov chain model to reconstruct likely sequences of variants that constitute real haplotypes in the metagenome. The source code is available at https://github.com/SamStudio8/gretel.

We run Gretel with the following commands, which are their default settings:

Using BWA-MEM, map the simulated reads to the reference sequence “{ref}”, and generate an output “{input_sam}” in the SAM format:

```
> bwa mem
    -R /path/to/{ref}
    /path/to/{prefix}.bwa.read1.fastq
    /path/to/{prefix}.bwa.read2.fastq
    >  /path/to/{input_bam}
```

Run Gretel:

```
    /path/to/gretel
    /path/to/{prefix}.raw.vcf.gz
    -s 1
    -e 4972
    --master /path/to/{ref}
    -o /path/to/{prefix}.gretel_output
```

Parameters:
/path/to/Gretel. The full path to the Gretel executable file.
/path/to/{prefix}.raw.vcf.gz. The full path to the VCF files, which must be compressed using bgzip and indexed with tabix.
-s. Start position.
-e. End position.
--master. Full path to the reference genome.
-o. Full path to the output directory.

**Human Haplotypes Reconstruction Tools:**

*Hippo*
Hippo (Haplotype estimation under incomplete prior information using pooled observations) uses a multinormal approximation of the likelihood observing genotyping data and a reversible-jump Markov chain Monte Carlo (rjMCMC) algorithm to estimate the existing haplotypes in the population and their frequencies.

We run Hippo with the following commands, which are their default settings:

```
>/path/to/hippo parameters_file
```

Parameters:
/path/to/hippo. Full path to hippo
parameters_file. Each row of the parameters file contains a label and the corresponding parameter value, separated by whitespace. An example of the parameter file is below:

```
data_file {prefix}
n_loci 25
n_pools 25
```

{prefix} The name of the data file. The data file contains one row for each pool, for a total of *n_pools* rows. Each row contains 1+*n_loci* integer entries separated by whitespace as shown:

*poolsize a_1 a_2 ... a_n_loci*, where *poolsize* is the number of individuals in the pool, and *a_i* is the number of alleles of type 1 at locus *i*, where *i = 1,...,n_loci*. Alleles are labelled either 0 or 1. An example of the data_file where *n_pools = 25*, *n_loci = 10*, and *poolsize = 100*:

```
100 50 83 22 20 48 53 82 29 20
41
100 91 56 52 18 73 94 53 66 18
74
… …
```

n_loci: (int) number of loci
n_pools: (int) number of pools
All other parameters are set to default values, excepting the three parameters described above.

*Approximate expectation maximization*

The approximate expectation maximization (AEM) algorithm infers haplotypes from pools using an approximate EM algorithm(Kuk, et al. 2009).

We run AEM using the following commands, which are their default settings:

```
> /path/to/aem(genodata, N=50)
```

Parameters:
genodata is an R matrix denoting the count of alternate alleles, e.g.:
matrix(c(50,83,22,20,48,53,82,29,20,41 91,56,52,18,73,94,53,66,18,74…), nrow=25, byrow=T)

N=50: The number of individuals in each pool

As AEM outputs the frequencies of all $2^N$ possible haplotypes, we selected only the 50 most frequent haplotypes for comparison.

**Processing Haplotype Reconstruction Output**

We converted all output data into a custom tab-delimited output format. The first row of the output file lists the haplotype IDs, and the second row lists their corresponding frequencies. The remaining rows record the position of each varying site and list the alleles of each haplotype at that site. Below is an example of the output format:

```
Hap     h0      h1      h2      h3      h4 …
Freq    0.02870493      0.02202937      0.03137516      0.06108144      0.12449933 …
0;211;211;0:1 0 1       1       1       1 …
0;231;231;0:1 1 0       1       0       1 …
0;310;310;0:1 0 1       1       0       0 …
… …
```

We evaluated the performance of each haplotype reconstruction tool by comparing their reconstructed haplotypes with the gold standard simulated haplotypes using the following command:

```
>java -jar PoolHapX.jar evaluate
```

We measured the accuracy of each tool by calculating the MCC and JSD of their reconstructed haplotypes.

#### SECTION VI: REAL DATA ANALYSES

##### *Plasmodium vivax* Analysis

We identified the haplotypes present in the hosts of multiple strains of *Plasmodium vivax*, a protozoan which causes malaria. Short sequence reads from previously published *P. vivax* strains(Hupalo, et al. 2016) were downloaded from ENA (https://www.ebi.ac.uk/ena) and mapped against a combined human (hg38) *P. vivax* (strain P01, version 39) genome using NextGenMap (v0.5.5)(Sedlazeck, et al. 2013) with the following command:

```
>ngm
    -r [genome, fasta]
    -1 [1st read pairs]
    -2 [2nd read pairs]
    -b
    -o [sample].bam
```

Read mappings were sorted, and duplicate reads were removed using SAMtools (v 1.9)(Li, et al. 2009):

```
>samtools sort [sample].bam > [sample].sorted.bam
>samtools index [sample].sorted.bam
>samtools rmdup -S [sample].sorted.bam [sample].rmdup.bam
>samtools index [sample].rmdup.bam
```

The read group names were changed using Picard tools (http://broadinstitute.github.io/picard):

```
>picard AddOrReplaceReadGroups
    I=[sample].rmdup.bam
    O=[sample].nrg.bam
    RGID='sample' RGLB=lib_
    'sample' RGPL=illumina
    RGPU=unit1 RGSM='sample'
```

Reads were split into separate *P. vivax* chromosomes using the SAMtools view function. For each chromosome we run:

```
>samtools view
    -bh [sample].nrg.bam 'chr' > [sample_chr].bam
```

Indels were realigned using the GATK package (v 4.0.11)(McKenna, et al. 2010):

```
>java -jar GenomeAnalysisTK.jar
    -T RealignerTargetCreator
    -R [fasta file of chromosome]
    -I [sample_chr].bam
    -o [sample_chr].forIndelRealigne, r.intervals
>java -jar GenomeAnalysisTK.jar
    -T IndelRealigner
    -R [fasta file of chromosome]
    -I [sample_chr].bam
    -targetIntervals [sample_chr].forIndelRealigner.intervals
```

```
	-o [sample_chr].realigned.bam
```

A CHR file is constructed for each chromosome to list the names of all resulting BAM files of the form [sample_chr].realigned.bam.

SNPs are called using GATK HaplotypeCaller, keeping SNPs with a QUAL score above 30.

```
>java -jar GenomeAnalysisTK.jar
	-R [fasta file of chromosome]
	-T HaplotypeCaller
	-I [chr, list of bam]
	--genotyping_mode DISCOVERY -o [chr].gatk.raw.vcf
>cat [chr].gatk.raw.vcf | bcftools filter --include 'QUAL>30 && MQ>30' > [chr].gatk.filter.vcf
```

SNPs are called using freebayes (v 1.2.0)(Garrison and Marth 2012), keeping only SNPs with a QUAL score above 30 (Danecek, et al. 2011):

```
>freebayes
	--bam [chr, list of bam]
	--min-alternate-count 5
	--fasta-reference [fasta file of chromosome]
	--dont-left-align-indels
	--vcf [chr].freebayes.raw.vcf
>vcftools
	--vcf [chr].freebayes.raw.vcf
	--minQ 30
	--stdout
	--recode
--recode-INFO-all > [chr].freebayes.filter.vcf
```

The VCF files produced by GATK and freebayes are merged together, keeping the genotyping information produced by freebayes:

```
>vcftools
	--vcf [chr].freebayes.filter.vcf
	--positions [chr].gatk.filter.vcf
	--stdout
	--recode --recode-INFO-all > [chr].union.vcf
```

Columns of VCF files are reordered so that samples appear in a consistent order across all files. We pick a random VCF file [template vcf] and reorder all other VCF files to match:

```
> vcf-shuffle-cols -t [template vcf] [chr].union.vcf | bgzip > final.[chr].vcf.gz tabix final.[chr].vcf.gz
```

VCF files for all individual chromosomes are combined together:

```
> vcf-concat final.[chr1].vcf.gz final.[chr2].vcf.gz (..) final.[chrN].vcf.gz > pvivax.vcf.gz
```

The *P. vivax* data was then obtained in the form of VCF files. The data was separated by chromosome and their respective BAM files were checked for coverage using SAMtools. The pool or strain ID for *P. vivax* was noted. The VCF files were then filtered for SNP (i.e., no indels), sequencing coverage of the pools, and SNP allele frequencies greater than 0.05. Once the VCFs were filtered they were formatted and analyzed by PoolHapX using its default parameters. PoolHapX was able to identify multiple haplotypes on different strains of *P. vivax*. The numbers of haplotypes are presented in **Supplementary Table 3** and an example of visualized frequencies are illustrated in **Supplementary Figure 9.**

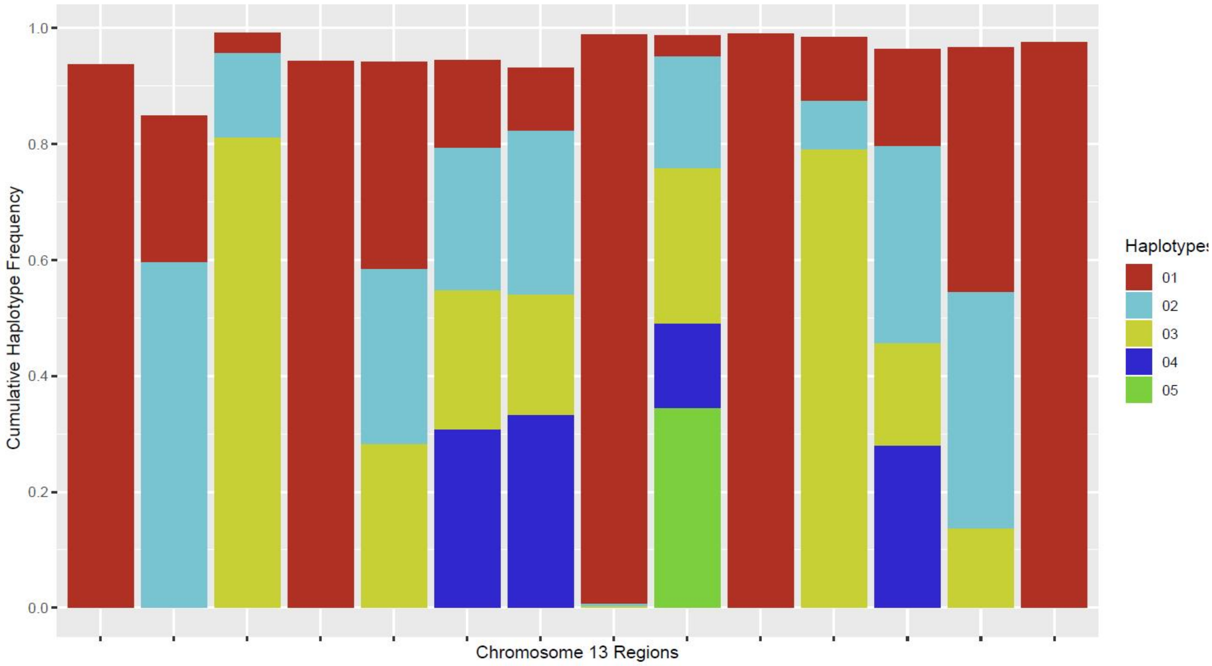


**Supplementary Fig. 9:** The graph shows the distribution of predicted haplotypes in Chromosome 13 of *Plasmodium vivax* strain SRS693934. The chromosome was divided into separate regions to enable PoolHapX to process the data. Each stacked bar in the graph represents one of these regions and each stack represents a haplotype. The height of a stack depicts the haplotype's frequency.

**Supplementary Table 3:** Each chromosome of each strain was divided into multiple regions. The average number of haplotypes identified per regions of each chromosome for each of the 49 strains of *Plasmodium vivax* is detailed below.

| Strain ID | Haplotypes per Region per Chromosome |
|---|---|
| **M_18_0048** | 4.94 |
| **M_18_0049** | 4.25 |
| **SRS258091** | 4.49 |
| **SRS258178** | 4.45 |
| **SRS258286** | 4.63 |
| **SRS363192** | 4.59 |
| **SRS365050** | 4.17 |
| **SRS417747** | 4.69 |
| **SRS693267** | 4.07 |

| | |
|---|---|
| **SRS693269** | 4.74 |
| **SRS693273** | 4.02 |
| **SRS693307** | 4.07 |
| **SRS693346** | 4.28 |
| **SRS693348** | 4.00 |
| **SRS693354** | 4.20 |
| **SRS693364** | 4.13 |
| **SRS693469** | 4.00 |
| **SRS693476** | 4.80 |
| **SRS693491** | 1.79 |
| **SRS693551** | 4.09 |
| **SRS693569** | 4.00 |
| **SRS693582** | 3.63 |
| **SRS693583** | 4.07 |
| **SRS693897** | 4.21 |
| **SRS693900** | 3.61 |
| **SRS693902** | 4.14 |
| **SRS693903** | 3.81 |
| **SRS693908** | 3.65 |
| **SRS693917** | 4.00 |
| **SRS693934** | 4.16 |
| **SRS693950** | 4.05 |
| **SRS693956** | 3.82 |
| **SRS693980** | 3.46 |
| **SRS694229** | 4.14 |
| **SRS694231** | 4.01 |
| **SRS694244** | 3.87 |
| **SRS694245** | 4.13 |
| **SRS694247** | 4.34 |
| **SRS694248** | 4.09 |
| **SRS694250** | 4.04 |
| **SRS694255** | 4.35 |
| **SRS694267** | 3.77 |
| **SRS696214** | 4.19 |
| **SRS696216** | 3.77 |
| **SRS696222** | 4.70 |
| **SRS819436** | 4.40 |
| **SRS819493** | 4.40 |
| **SRS819716** | 3.63 |
| **SRS819740** | 4.35 |

##### Infant Gut Colonization Time Series Analysis

StrainEst focuses on species of interest by defining their population structure through a clustering of their genetic variants profiles. In each cluster of SNP profiles, StrainEst can only detect the representative strain which is central to that cluster. However, StrainEst cannot investigate the subtle difference between haplotypes and reconstruct them *de novo*. In addition, while each cluster contains strains that are similar across the whole genome, the strains within the cluster may differ greatly in specific regions. StrainEst uses a popular time series of infant gut colonization data to demonstrate its ability to analyze real metagenomic data. In this dataset, the dominant strain was 504_SEPI at the start of the sampling period (15 to 16 days), after which the dominant strain switches to 236_SEPI (17 days onward). This finding is consistent with the conclusion of the original paper(Sharon, et al. 2013). In our analysis, we apply PoolHapX to the same data and map our predicted haplotypes back to the three main strains, to examine whether the aggregated haplotype frequencies follow a similar trend to the original estimation(Albanese and Donati 2017) based on reference templates of known strains.

The FASTA files of the three main strains 236_SEPI, 504_SEPI, and 634_SEPI, were downloaded from https://www.ncbi.nlm.nih.gov/genome/genomes/155#. The *Staphylococcus epidermidis* reference genomes (SR.fasta) were downloaded from https://www.ncbi.nlm.nih.gov/genome/155?genome_assembly_id=299299.

We align the *S. epidermidis* genomes using the "mapgenomes" function of StrainEst:

```
> strainest mapgenomes ref/236_SEPI.fa ref/504_SEPI.fa ref/634_SEPI.fa  SR.fasta MR.fasta
```

Below is an example of the output FASTA format which aligns the reference templates:

```
GCA_001073525.1_ASM107352v1_genomic.fna,Ref,236_SEPI.fa,504_SEPI.fa,634_SEPI.fa
4,G,A,G,G
184,C,C,T,C
598,A,A,T,A
635,A,A,C,A
868,T,T,C,T
1093,G,A,A,G
1153,T,A,A,T
1156,C,T,T,C
```

We then use our pipeline to call SNPs for the 11 available samples from 11 different time points. From the output, the variants with minor allele frequency less than 0.05 were removed. In total there are 10,975 SNPs. As PoolHapX cannot process such a large number of SNPs, and we also expect the haplotype structure in different regions to vary, we divided the 10,975 SNPs into 110 regions so that each region contains around 100 SNPs and can be quickly processed by PoolHapX.

The running parameters of PoolHapX can be found in the property file in our GitHub (https://github.com/theLongLab/PoolHapX/tree/master/Simulation_And_Analysis). Average numbers of haplotypes at each time point are presented in **Supplementary Table 4.**

**Supplementary Table 4:** Average number of haplotypes at 11 time points. The *S. epidermidis* whole genome is divided into 110 regions. The average number of haplotypes in each region at different time points is shown below.

| Day | Average Number of Haplotypes |
|---|---|
| **15** | 12.5 |
| **15.5** | 12.7 |
| **16** | 10.7 |
| **17** | 5.9 |
| **17.5** | 12.3 |
| **18** | 3.9 |
| **19** | 11.6 |
| **22** | 9.9 |
| **22.5** | 9.9 |
| **23** | 7.2 |
| **24** | 6.8 |

We then conduct a comparison between the template-based inference by StrainEst and PoolHapX's *de novo* inference. The abundance of the three *S. epidermidis* strains 236_SEPI, 504_SEPI, and 604_SEPI in the infant gut colonization time series was predicted by StrainEst using a template-based method. However, StrainEst only predicted the abundance of the three representative strains, which is not directly comparable with the fine-scale haplotypes predicted by PoolHapX. We therefore align all the haplotypes predicted by PoolHapX to their nearest representative *S. epidermidis* strains with the lowest MCC, in order to check whether the abundance of the three strains predicted by both tools have a similar trend for the majority of regions. Of the 110 regions, in around 37, 40, and 17 regions of the three respective strains we observed significantly difference between strain abundance predicted by StrainEst and the aggregated frequencies of PoolHapX-predicted haplotype frequencies (two-tailed paired T-test, p-value < 0.05). The rest regions, which are the majority enjoy similar frequencies. Overall, the average of the 110 regions shows a fairly similar pattern over the longitudinal samples. This result is depicted in **Figure 5** of the main text.

#### HIV Within-Patient Evolutionary Analyses

*HIV genomic data*

We selected patient 1 from the dataset generated by Zanini *et al*(Zanini, et al. 2015), due to the number of time-points available and the relative completeness of genomic coverage at each time-point. For more information about the inclusion criteria of study patients, the clinical origins of the sample data, and sequencing procedures, we refer the reader to Zanini *et al*(Zanini, et al. 2015). Patient #1 is a female with HIV-1 subtype 01_AE. The first blood plasma sample was taken from an estimated 122 days post-infection, and the last sample at 8.2 years post-infection. By applying PoolHapX to the data, we reconstructed haplotypes in each time point and the number and frequencies of haplotypes are listed in **Supplementary Table 5.**

**Supplementary Table 5:** Number of haplotypes and their frequencies at 12 time points for HIV patient 1.

| Time Point | Number of haplotypes | Haplotype abundance |
|---|---|---|
| **1** | 2 | 0.51;0.49 |
| **2** | 12 | 0.17;0.16;0.13;0.12;0.12;0.1;0.06;0.04;0.04;0.03;0.03;0.02 |
| **3** | 13 | 0.22;0.1;0.1;0.09;0.07;0.07;0.07;0.07;0.06;0.05;0.04;0.02;0.01 |
| **4** | 13 | 0.18;0.14;0.1;0.09;0.07;0.07;0.07;0.06;0.06;0.05;0.04;0.03;0.02 |
| **5** | 13 | 0.24;0.11;0.1;0.09;0.07;0.07;0.07;0.06;0.05;0.05;0.04;0.04;0.02 |
| **6** | 12 | 0.19;0.14;0.11;0.1;0.09;0.08;0.08;0.07;0.06;0.06;0.04;0.02 |
| **7** | 13 | 0.26;0.19;0.11;0.11;0.08;0.05;0.05;0.04;0.04;0.04;0.04;0.03;0.01 |
| **8** | 10 | 0.17;0.15;0.14;0.13;0.09;0.07;0.07;0.07;0.06;0.05 |
| **9** | 12 | 0.15;0.13;0.11;0.09;0.09;0.09;0.08;0.05;0.05;0.05;0.04;0.04 |
| **10** | 12 | 0.21;0.15;0.1;0.1;0.09;0.08;0.07;0.06;0.05;0.04;0.04;0.02 |
| **11** | 11 | 0.25;0.14;0.12;0.12;0.07;0.07;0.07;0.07;0.05;0.03;0.02 |
| **12** | 13 | 0.29;0.12;0.11;0.11;0.1;0.07;0.06;0.04;0.04;0.04;0.04;0.04;0.02 |

Within each time-point-specific set of haplotypes, we calculate the site-specific extended haplotype heterozygosity (EHH) at every SNP position. To examine the rate and dynamics of linkage decay within each gene, we calculated at each time-point the size of regions around each SNP position in which site-specific extended haplotype heterozygosity (EHHS) is equal to or larger than 0.5.

*EHH-based analyses of the within-patient population*

Scripts for generating well-formatted files from PoolHapX output files and apply extended haplotype homozygosity-based statistics to the reconstructed haplotypes, are available in our GitHub repository (https://github.com/theLongLab/PoolHapX/tree/master/Simulation_And_Analysis/HIV_analysis_code). The scripts include detailed descriptions of our parameters and settings.

Using the sequence alignment and variant calling procedure described in the previous sections, 208 SNP positions were identified from the sequencing data from all 12 time-points. The sequencing reads from each time-point were re-mapped to the consensus whole-HIV genome sequence from patient 1's first sample to generate consistent SNP positions. The reads from the SAM files of each time-point were annotated with either a 0 or a 1 at each SNP position, depending on whether they carried the reference or an alternate allele. PoolHapX was subsequently applied to reconstruct haplotypes jointly across the longitudinal samples of patient 1, with a set of haplotypes reconstructed for each time-point (default parameters are used).

For each time-point, a single PoolHapX output file was generated and converted to MS(Ewing and Hermisson 2010) output files. The within-population frequencies of each haplotype scaled to the number of viral particles per milliliter of patient plasma(Zanini, et al. 2015). MS is generally used to generates samples with the assumption of neutral evolution, and the default output format encodes haplotypes as a string of 0s (reference allele) and 1s (alternate allele) at each SNP position(Hudson 2002).

The R package rehh (version 3.0) was applied to survey linkage patterns within a single time-point, as well as changes to linkage patterns across the duration of infection monitoring(Gautier, et al. 2017). Several long-range haplotype-based evolutionary statistics related to extended haplotype homozygosity(Sabeti, et al. 2002) were used to quantify the type and magnitude of selection.

Within each set of haplotypes specific to a time-point, the site-specific extended haplotype heterozygosity (EHHS) was calculated at every SNP position(Sabeti, et al. 2007). EHHS quantifies the probability that any pair of randomly chosen haplotypes are homozygous over the region between the focal SNP position and another SNP position. As with(Gautier and Vitalis 2012; Gautier, et al. 2017), the EHHS of the region between focal SNP position s and another SNP position *t* is

$$EHHS_{s,t} = \frac{1}{n_s(n_s - 1)} \sum_{k=1}^{K_{s,t}} n_k(n_k - 1)$$

$n_s$ is the number of haplotypes at SNP position s, which is equivalent to the number of haplotypes at that time-point since all PoolHapX-reconstructed haplotypes are complete across the span of SNP positions. $K_{s,t}$ is the number of different kinds of haplotypes that are homozygous over the region between SNP positions s and t (inclusive). $n_k$ is the number of haplotypes that are in the category $K_{s,t}$.

To examine the rate and dynamics of linkage decay within each gene, we calculated at each time-point the size of regions around each SNP position where EHHS is equal to or larger than 0.5. The size of the region between the rightmost and leftmost positions such that $EHHS_{s,t} >= 0.5$ (same notation as above) was plotted for each time-point and each gene.

To search for regions of positive selection within the reconstructed genome, we calculated the integrated haplotype score (iHS) for each time-point, as well as cross-population EHH scores (XP-EHH) between each time-point. The rehh function scan_hh() calculates the integrated haplotype homozygosity (iHH) for both the ancestral and derived alleles respectively, and the rehh function ihh2ihs() calculates the iHS from the ancestral and derived iHH. The iHH of an allele is the area under the EHH curve of that allele, which quantifies the decay of linkage around that allele. The minimal threshold of EHH calculation (limehh) is set to 0, and EHH integration continues to the end of the reconstructed region (for patient 1 haplotypes, this is from positions 1 to 6078).

The iHS of a SNP position calculates the relative homozygosity around the derived allele relative to the ancestral allele. When the distribution of iHS from other SNP positions with similar allele frequencies is standardized, p-values can be calculated for each SNP position to indicate outliers where either the derived or ancestral allele at that position is significantly more homozygous than the other. Alleles are binned by frequency in increments of 0.1, and the p-value threshold for outlier status was set to 0.05. The equations are the same as the ones in Gautier *et al* (Gautier, et al. 2017).

$$uniHS(s) = ln\left(\frac{iHH_a(s)}{iHH_d(s)}\right)$$

$$iHS(s) = \frac{uniHS(s) - mean(uniHS|p_s)}{sd(uniHS|p_s)}$$

uniHS(s) is the un-standardized log-ratio of ancestral iHH to derived iHH at SNP position s. mean(uniHS|$p_s$) and sd(uniHS|$p_s$) refer to the mean and standard deviation of SNP positions belonging to the same allele frequency bin ($p_s$), so that their frequencies are within 0.1 of each other. We standardize iHS(s) by subtracting the mean and dividing by the standard deviation of uniHS.

Selection tends to produce regions with multiple outliers(Tang, et al. 2007). To identify regions where homozygosity is maintained to a significant degree relative to the rest of the genome at a given time-point, the rehh function calc_candidate_regions() was applied to intervals of 250 base pairs (bp) containing two or more SNP positions with outlier iHS scores, in a sliding window fashion with 125 bp of allowable overlap. If overlapping 250 bp intervals each contain two or more outliers, the intervals are merged and reported as a single region. We chose the interval size of 250 bp since it was slightly less than the average window size of EHHS >= 0.5 in the envelope gene. This is the smallest average window size out of all genes analyzed (gag, pol, vif, vpu, tat, env).

To identify sites where haplotype homozygosity surrounding one allele or the other has undergone significant longitudinal change, the XP-EHH of each SNP position was calculated by comparing its integrated EHHS (analogous to iHH, but with EHHS instead of EHH) at time-point *i + 1* relative to the previous time-point *i*. As with iHS, integrated EHHS must also be standardized relative to the rest of the inter-time-point comparisons(Gautier, et al. 2017).

$$unXP - EHH(s) = ln\left(\frac{iES_{pop1}(s)}{iES_{pop2}(s)}\right)$$

$$XP - EHH(s) = \frac{unXP - EHH(s) - mean\big(unXP - EHH(s)\big)}{sd\big(unXP - EHH(s)\big)}$$

unXP-EHH(s) is the unstandardized log-ratio of the population 1 iES (at time-point *i*) to the population 2 iES (at time-point *i + 1)*, at SNP position *s*. mean(unXP-EHH(s)) and sd(unXP-EHH(s)) refer to the mean and standard deviation of SNP positions within the same allele frequency bin ($p_s$) of width 0.1. We standardize XP-EHH(s) by subtracting the mean and dividing by the standard deviation of unXP-EHH(s).

Similar to the iHS regions obtained above, regions where homozygosity is significantly increased relative to the previous time-point are calculated by applying the rehh function calc_candidate_regions() to the XP-EHH scores of each time-point's SNP positions. The user-defined parameters are the same as described for the outlier iHS regions above.

**Longitudinal Analyses of Patient HIV Populations Reveal General and Gene-specific Linkage Patterns**

Typically, prior investigations estimated the recombination rate in genes targeted by antiretroviral drugs or immune selection either by clonal sequencing, or indirectly through longitudinal SNP comparisons(Messer 2013; Rouzine, et al. 2014). To demonstrate the utility of haplotype reconstruction for evolutionary analyses of complex populations, we explored longitudinal patterns of linkage across the HIV genome. For more sample details and a thorough analysis of SNP-based trends in genetic diversity and divergence, we refer the reader to Zanini et al (Zanini, et al. 2015).

*The applicability of EHH-based selection statistics to within-patient HIV populations*

Given its high mutation rate(Rambaut, et al. 2004), HIV does not necessarily follow the assumption of identity by descent, and the maintenance of regional homozygosity may not indicate population expansion due to the advantage of a single allele. The same mutation may have arisen multiple times within the span of the longitudinal samples. As a result, there may be multiple haplotypes that are identical due to random drift and not by descent, which dilutes the signal of positive selection for that haplotype. Additionally, most SNP positions are assumed to be governed by neutral selection. Though the genes surveyed in this analysis are not as immunogenic as the envelope gene (of which only a few base pairs are reconstructed), several proteins in Gag, such as p24, do accumulate cytotoxic T-lymphocytes (CTL) escape mutations at some SNP positions due to their physical location on the exterior of the HIV capsid(Prince, et al. 2012). Thus, it is likely, although not certain, that the rehh-defined p-value (against a null hypothesis of no selection when all positions follow a Gaussian distribution) holds.

However, the maintenance of any particular linkage block is difficult under a high rate of recombination, and therefore the strength of linkage in a block of alleles relative to the whole genome is still indicative of the relative fitness of that regional haplotype. Thus, EHH-based summary statistics can still be considered as general tests for assessing positive selection via unusually long-range LD.

The integrated haplotype score (iHS) and cross-population EHH score (XP-EHH) are both used because they are complementary. Whereas XP-EHH, which is similar to Tang et al.'s Rsb statistic(Tang, et al. 2007), has the power to detect selective sweeps that have reached fixation, iHS is better for comparing sweeps which are in progress, in which case there exist two alleles with intermediate frequencies.

*Linkage decays extremely rapidly post-infection, but does not decrease monotonically over time*

Within each time-point-specific set of haplotypes, we calculated the site-specific extended haplotype heterozygosity (EHHS) at every SNP position(Sabeti, et al. 2002). EHHS measures the decay of haplotype homozygosity as a weighted combination of the ancestral and the derived allele frequencies. An EHHS of 0.5 indicates that 50% of the haplotypes carrying an allele at the focal position also carry the same allele at the distant position.

Although the Gag-Pol Polyprotein gene is in complete LD at the first time-point, over a year later at time-point 2 (562 days post infection (DPI)), the window of SNP positions where the EHHS is larger than 0.5 has shrunk to at most 1841 bp at positions 1412 to 1670, which overlap the p7, p1, and p6 genes in Gag. However, even at the first time-point, Gag-Pol SNP positions are not in complete linkage with the genes downstream of Pol which we reconstructed, including the genes Tat, Vif, Vpu, and the first 386 bp of Env (**Supplementary Fig. 10**).

*Linkage is measured on the order of several hundred base-pairs consistently across two-thirds of the HIV genome*

To examine the rate and dynamics of linkage decay within each gene, we calculated the average size of regions around each SNP position where EHHS is larger than 0.5. We plotted the longitudinal decay of homozygosity for each focal position (**Figure 6C, D** and **Supplementary Fig. 10**). Since EHHS decays monotonically (**Figure 6A, B** and **Supplementary Fig. 10**) with increasing genomic distance from each SNP position, the sizes of these regions represent the average size of a linkage block at that time-point.

Strikingly, for all genes the decrease in average window size appears to level out around day 1282, approximately 3.5 years post infection. After this point, the average window size across all positions in the remaining time-points is 217 bp, and the standard deviation is 170 bp. In intermediate progressors, this is the time-point in disease progression where CD4+ T-cell counts start to decline(Langford, et al. 2007). After this point, instead of decreasing monotonically towards an asymptotic value, the average window size fluctuates around a range constrained to several hundred bp, depending on the gene. These fluctuations can occur quickly, with increases and decreases in the hundreds of bp on scales as low as a month (**Supplementary Fig. 10**).

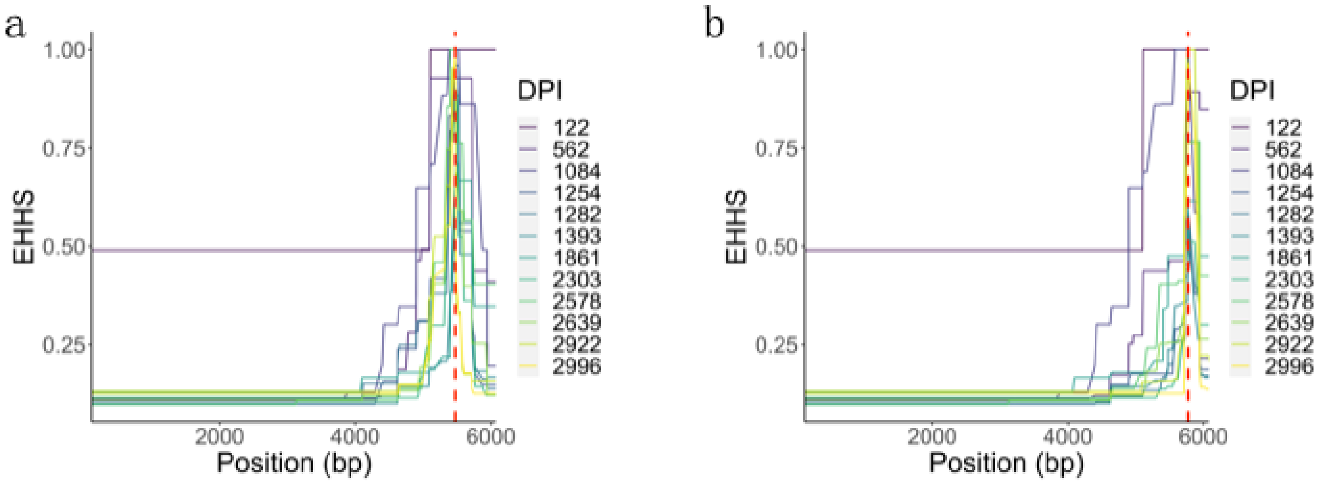


**Supplementary Fig. 10:** The decay of EHHS around each SNP position in reconstructed HIV-1 haplotypes occurs rapidly during the acute phase of infection. The dashed red line indicates the location of the focal SNP position. A) Position 5474 (Vpu gene). B) Position 5769 (Env gene, found in the env peptide).

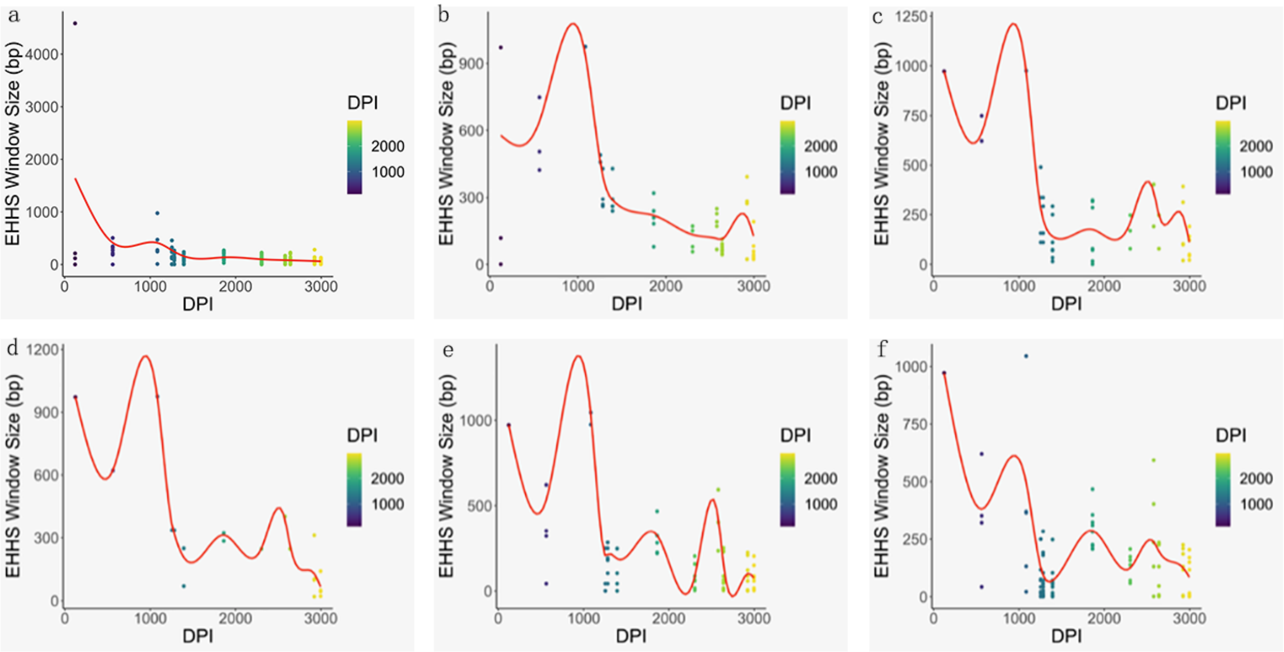


**Supplementary Figure 11:** The windows where EHHS ≥ 0.5 fluctuate in size around gene-specific averages. The solid red line indicates a weighted mean across the positions in the gene. DPI refers to estimated days post-infection. Each dot represents the window size around at least one position. A) Vif. B) Vpr. C) Tat. D) Rev. E) Vpu. F) 5' region of the envelope gene (Env).

Within Gag and Pol, there is substantial heterogeneity in average window size, with the downstream regions of Gag and Pol largely fluctuating between 0 and 250 bp (**Figure 6C,D**). Although the first half of the Pol gene, which spans from 1551 to 3470 bp fluctuates in window size between 0 and 500 bp, the latter half of the Pol gene behaves more similarly to the genes downstream of Gag-Pol, with the average size fluctuating between 0 and 250 bp. Even within Gag and Pol, where the rate of nonsynonymous evolution is slower than in accessory and envelope proteins(Zanini, et al. 2015), there is no SNP position with a consistent window of high EHHS.

Zanini et al.(Zanini, et al. 2015) previously characterized linkage using D', an approach that relies on the average allelic content residing on a genomic fragment. After reconstructing HIV haplotypes from the first 6000 bp, we find that the average linkage block size is at least twice as large as previously estimated, and the standard deviation is nearly as large as the average. Nevertheless, our estimate of the recombination rate, which is based on an average linkage block size of 217 bp and an average separation of 261 days between sampling times, is $1.76015 \times 10^{-5}$ crossovers/day, which agrees with both the study of Zanini et al. as well as several others(Shriner, et al. 2004; Neher and Leitner 2010; Batorsky, et al. 2011; Song, et al. 2018). Furthermore, the limited range of sizes observed in the average linked regions indicate that the first 6000 bp of the HIV genome appear to be governed by similar recombination rates after approximately three years post-infection.

*In the absence of drug selection in immunogenic genes, selective sweeps still occur occasionally and recurrently in the same regions*

We have conducted a search for regions of positive selection within the reconstructed genome where selective sweeps could have taken place. To identify regions that are positively selected relative to other regions within the same time-point, we identified 500 bp windows where there are 2 or more SNP positions that were outliers in terms of their integrated haplotype score (iHS). The iHS measures whether a derived allele displays significantly higher extended haplotype homozygosity (EHH) than the ancestral allele, or vice versa(Sabeti, et al. 2007) (**Supplementary Table 6**). To identify positively selected regions relative to the previous time-point, we identified 500-bp windows where there are 2 or more SNP positions that were outliers in terms of their cross-population EHH (XP-EHH)(Sabeti, et al. 2007) (**Supplementary Table 7**).

**Supplementary Table 6:** Regions with outlier integrated haplotype scores (iHS) relative to the rest of the reconstructed genome at a specific time-point. DPI refers to the number of days post-infection that the sample was taken. Start and end refer to the position relative to the patient's consensus founder (see Zanini et al more information)(Zanini, et al. 2015). Num. Extr. Positions refers to the number of SNP positions with outlier iHS within the region. Max. iHS and p-value refer to the iHS within the region with the largest magnitude and its relative significance.

| DPI | Start | End | Gene | Num. Extr. Positions | Tot. Markers | Max. iHS | p-value |
|---|---|---|---|---|---|---|---|
| **1254** | 5250 | 5625 | Vpr,Vpu | 2 | 9 | 2.583 | .010 |
| **1393** | 5625 | 5875 | Vpu,Env | 2 | 5 | 2.285 | .022 |
| **2639** | 2125 | 2375 | Pol | 2 | 6 | -2.532 | .011 |
| **2639** | 5250 | 5625 | Vpr,Vpu | 2 | 18 | 1.988 | .047 |
| **2922** | 4875 | 5625 | Vif,Vpu | 7 | 33 | -2.963 | .003 |

**Supplementary Table 7:** Regions with outlier cross-population extended haplotype homozygosity scores (XP-EHH) relative to the same region at the previous time-point. DPI refers to the number of days post-infection that the sample was taken. Start and end refer to the position relative to the patient's consensus founder (see Zanini et al for more information)(Zanini, et al. 2015). Num. Extr. Positions refers to the number of SNP positions with outlier XP-EHH within the region. Max. XP-EHH and p-value refer to the XP-EHH within the region with the largest magnitude and its relative significance.

| DPI | Start | End | Gene | Num. Extr. Positions | Tot. Markers | Max. XP-EHH | p-value |
|---|---|---|---|---|---|---|---|
| **562** | 3250 | 3625 | Pol | 2 | 15 | -3.103 | .002 |
| **562** | 4375 | 4875 | Pol,Vif | 10 | 16 | -3.081 | .002 |
| **1084** | 1250 | 1625 | Gag | 2 | 20 | -2.089 | .037 |
| **1084** | 5625 | 5875 | Vpu,Env | 2 | 8 | 2.443 | .015 |
| **1254** | 500 | 875 | Gag | 2 | 14 | 2.109 | .035 |
| **1254** | 1000 | 1375 | Gag | 5 | 12 | 2.559 | .010 |

| | | | | | | | |
|---|---|---|---|---|---|---|---|
| **1254** | 5500 | 6000 | Rev,Env | 4 | 21 | -3.119 | .002 |
| **1282** | 3250 | 3625 | Pol | 3 | 15 | 3.205 | .001 |
| **1393** | 5250 | 5625 | Vpr,Vpu | 5 | 18 | -3.010 | .003 |
| **1861** | 5750 | 6250 | Env | 11 | 13 | 2.629 | .009 |
| **2303** | 1125 | 1500 | Gag | 3 | 22 | 2.415 | .016 |
| **2639** | 3000 | 3375 | Pol | 4 | 16 | -2.797 | .005 |
| **2639** | 4500 | 5000 | Pol,Vif | 5 | 26 | 3.061 | .002 |
| **2922** | 1 | 250 | 5’ LTR | 2 | 2 | 2.205 | .027 |
| **2922** | 5250 | 5750 | Vpr,Vpu | 6 | 23 | -2.383 | .017 |
| **2996** | 1250 | 1625 | Gag | 8 | 20 | 3.013 | .003 |

The region approximately between positions 5250 to 5625, which overlaps the Vpr and Vpu genes, is repeatedly identified as an outlier (**Supplementary Table 1**). The surrounding sequence space, which overlaps neighboring genes such as Vif and Env, is also included. Though the region between Pol and Env is not uniformly identified as an outlier at all time-points, its consistent identification as a site of selective sweeping (relative to the rest of the genome) flags it as a location to further investigate estimates of baseline recombination rates, especially given that none of those proteins are immunogenic. The exception in the set is the region between positions 2125 and 2375 identified at time-point 10 (day 2639), which is near the 5’ end of the reverse transcriptase gene in Pol. It is unknown why this occurs.

In contrast, when comparing regions between time-points, there is both significant heterogeneity in the locations of the outlier regions, as well as limited consistency in terms of the affected gene between adjacent time-points (**Supplementary Table 2**). Regions that are recurrently swept include positions 5500 – 6000 that overlap Vpu and parts of Vpr and Env, and positions 3250 – 3625 in Pol that overlap the p15 gene, which undergo sweeps at multiple time-points throughout the 10 year span of the study data. Interestingly, different regions of Gag appear as the outlier at multiple time-points, accounting for 5 out of 16 entries and appearing at 4 out of 12 time-points.

#### SECTION VII: OTHER SUPPLEMENTARY FIGURES CITED IN THE MAIN TEXT

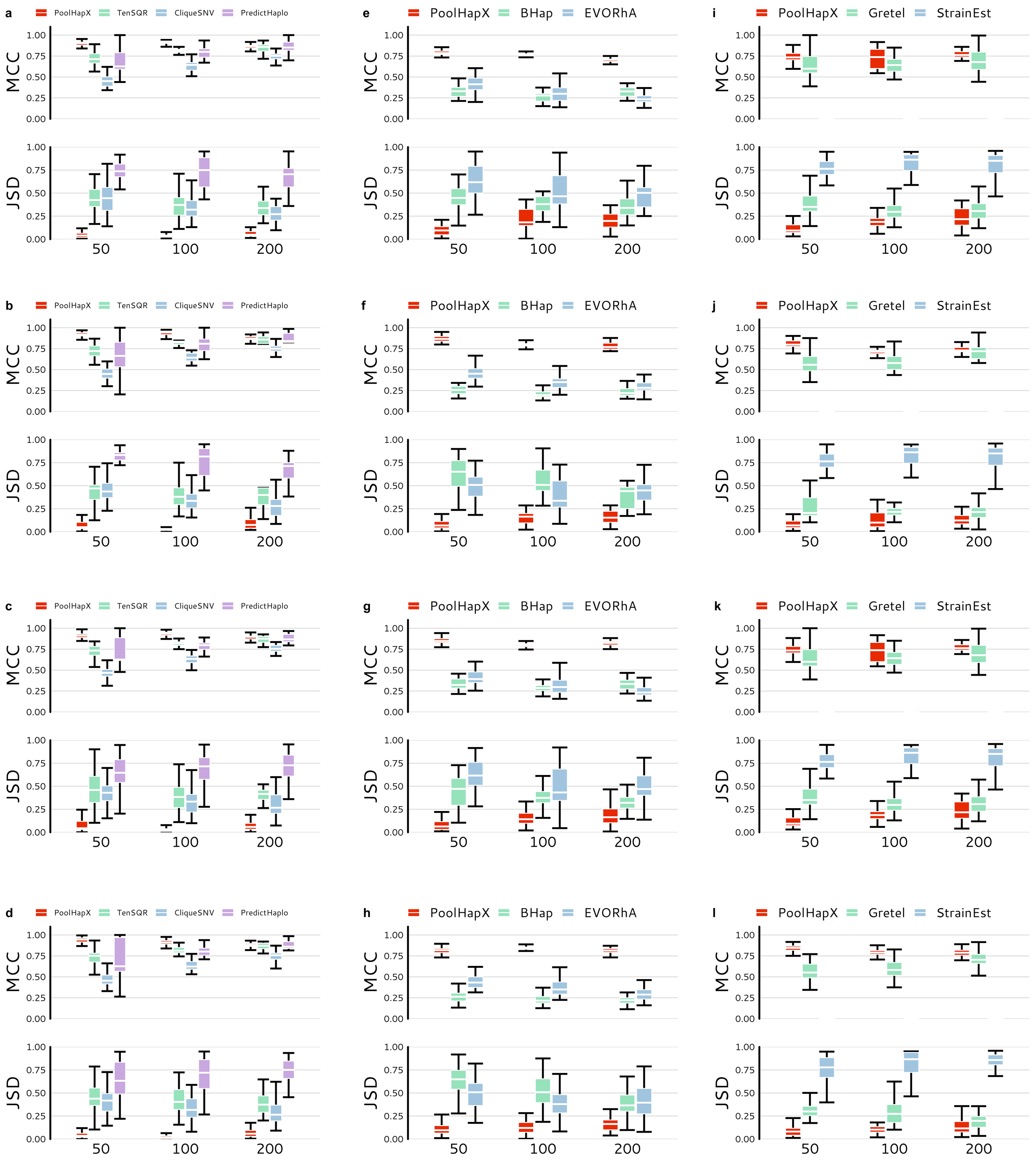

**Supplementary Fig. 12: Comparison between PoolHapX and existing haplotype reconstruction tools**. In all panels, the upper set of box plots show the accuracy of haplotype identities (MCC) and the lower set shows the accuracy of haplotype frequencies (JSD). The x-axis denotes number of genetic variants in the haplotype. Boxes extend to the first and third quartile, and whiskers extend to the upper and lower value. **Figures a-d** compare PoolHapX against the viral haplotype reconstruction tools TenSQR, PredictHaplo and CliqueSNV. **a.** Sequencing coverage per pool = 2500X, number of pools =

25. Number of haplotypes for 50 loci, 100 loci and 200 loci are 32, 54 and 33, respectively. **b.** Sequencing coverage per pool = 5000X, number of pools = 25. Number of haplotypes for 50 loci, 100 loci and 200 loci are 32, 54 and 33, respectively. **c.** Sequencing coverage per pool = 2500X, number of pools = 50. Number of haplotypes for 50 loci, 100 loci and 200 loci are 41, 73 and 42, respectively. **d.** Sequencing coverage per pool = 5000X, number of pools = 50. Number of haplotypes for 50 loci, 100 loci and 200 loci are 41, 73 and 42, respectively. **Figures e-h** compare PoolHapX against the bacterial haplotype reconstruction tools Bhap and EVORhA. **e.** Sequencing coverage per pool = 100X, number of pools = 25. Number of haplotypes for 50 loci, 100 loci and 200 loci are 36, 31 and 36, respectively. **f.** Sequencing coverage per pool = 250X, number of pools = 25. Number of haplotypes for 50 loci, 100 loci and 200 loci are 36, 31 and 36, respectively. **g.** Sequencing coverage per pool = 100X, number of pools = 50. Number of haplotypes for 50 loci, 100 loci and 200 loci are 42, 43 and 43, respectively. **h.** Sequencing coverage per pool = 250X, number of pools = 50. Number of haplotypes for 50 loci, 100 loci and 200 loci are 42, 43 and 43, respectively. **Figures i-l** compare PoolHapX against the metagenomic haplotype reconstruction tools Gretel and StrainEst. For StrainEst, it only reconstructed a reference haplotype, there is no TP value, so the MCC value cannot be calculated. Therefore, for MCC, we only showed the comparisons between PoolHapX and Gretel. **i**. Sequencing coverage per pool = 25X, number of pools = 25. Number of haplotypes for 50 loci, 100 loci and 200 loci are 30, 30 and 31, respectively. **j.** Sequencing coverage per pool = 50X, number of pools = 25. Number of haplotypes for 50 loci, 100 loci and 200 loci are 30, 30 and 31, respectively. **k.** Sequencing coverage per pool = 25X, number of pools = 50. Number of haplotypes for 50 loci, 100 loci and 200 loci are 39, 37 and 36, respectively. **l.** Sequencing coverage per pool = 50X, number of pools = 50. Number of haplotypes for 50 loci, 100 loci and 200 loci are 39, 37 and 36, respectively.

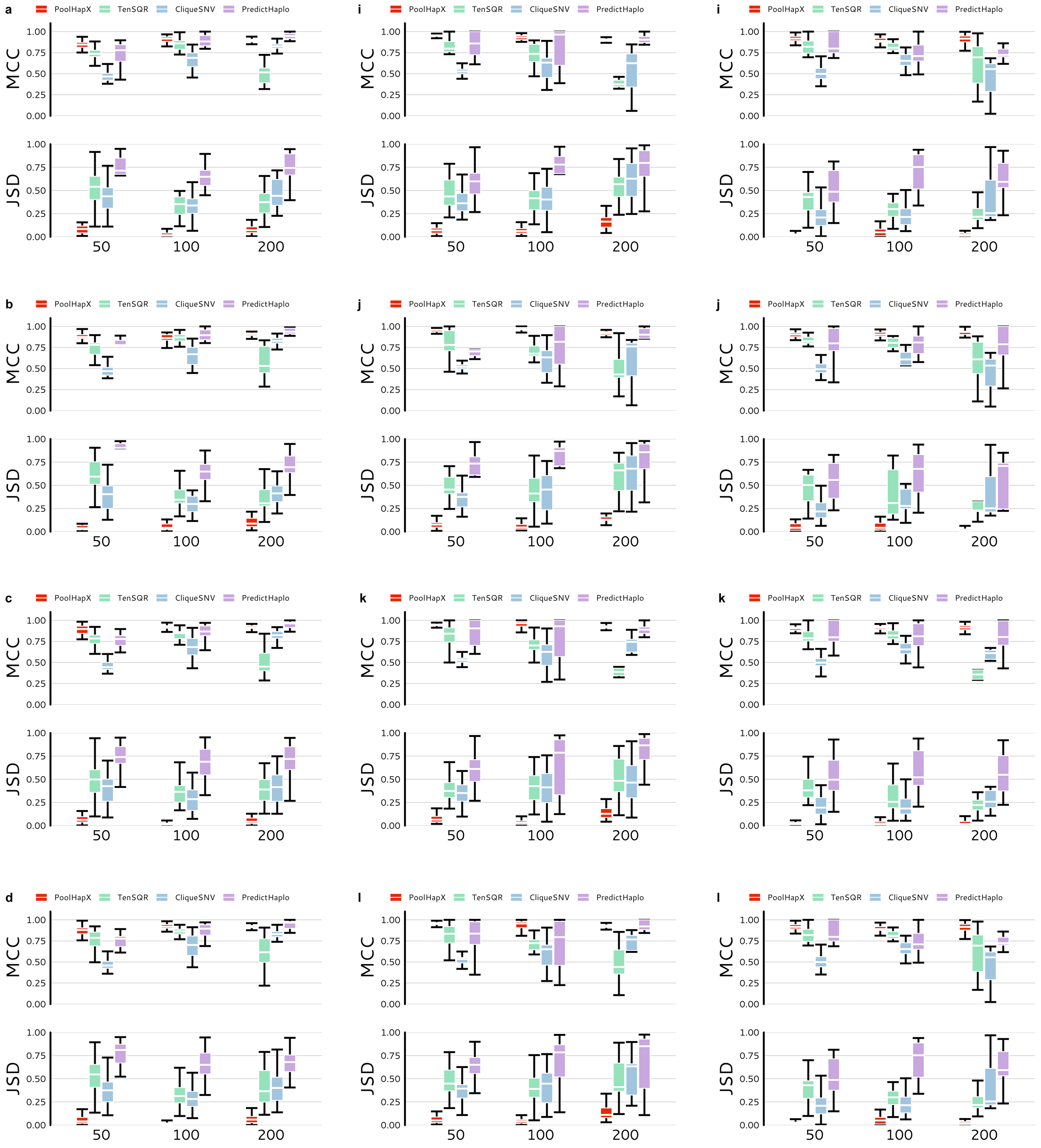


**Supplementary Fig. 13: PoolHapX is robust to the within-host changes due to selective forces**. In all panels, the upper set of box plots show the accuracy of haplotype identities (MCC) and the lower set shows the accuracy of haplotype frequencies (JSD). The x-axis denotes number of genetic variants in the haplotype. Boxes extend to the first and third quartile, and whiskers extend to the upper and lower value. Comparing to viral reconstruction tools, TenSQR, PredictHaplo and CliqueSNV. **Figures a-d** compare haplotype reconstruction tools under the scenario of negative selection. **a.** Sequencing coverage per pool = 2500X, number of pools = 25. Number of haplotypes for 50 loci, 100 loci and 200 loci are 31, 31 and 42, respectively. **b.** Sequencing coverage per pool = 5000X, number of pools = 25. Number of haplotypes for 50 loci, 100 loci and 200 loci are 31, 31 and 42, respectively. **c.** Sequencing coverage per pool = 2500X, number of pools = 50. Number of haplotypes for 50 loci, 100 loci and 200 loci are 36, 36 and 45, respectively. **d.** Sequencing coverage per pool = 5000X, number of pools = 50. Number of haplotypes for 50 loci, 100 loci and 200 loci are 36, 36 and 45, respectively. **Figures e-h** compare haplotype reconstruction tools under the scenario of positive selection. **e.** Sequencing coverage per pool = 2500X, number of pools = 25. Number of haplotypes for 50 loci, 100 loci and 200 loci are 32, 40 and 77, respectively. **f.** Sequencing coverage per pool = 5000X, number of pools = 25. Number of haplotypes for 50 loci, 100 loci and 200 loci are 32, 40 and 77, respectively. **g.** Sequencing coverage per pool = 2500X, number of pools = 50. Number of haplotypes for 50 loci, 100 loci and 200 loci are 41, 50 and 104, respectively. **h.** Sequencing coverage per pool = 5000X, number of pools = 50. Number of haplotypes for 50 loci, 100 loci and 200 loci are 41, 50 and 104, respectively. **Figures i-l** compare haplotype reconstruction tools under the scenario of selective sweep. **i.** Sequencing coverage per pool = 2500X, number of pools = 25. Number of haplotypes for 50 loci, 100 loci and 200 loci are 21, 30 and 24, respectively. **j.** Sequencing coverage per pool = 5000X, number of pools = 25. Number of haplotypes for 50 loci, 100 loci and 200 loci are 21, 30 and 24, respectively. **k.** Sequencing coverage per pool = 2500X, number of pools = 50. Number of haplotypes for 50 loci, 100 loci and 200 loci are 23, 36 and 27, respectively. **l.** Sequencing coverage per pool = 5000X, number of pools = 50. Number of haplotypes for 50 loci, 100 loci and 200 loci are 23, 36 and 27, respectively.

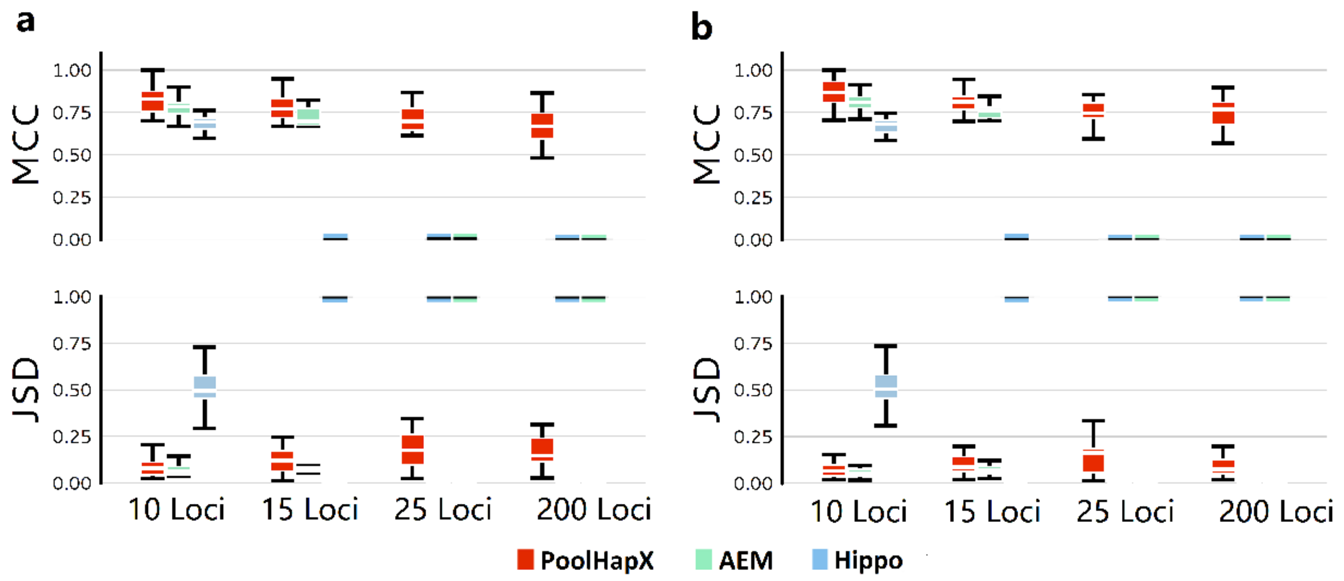


**Supplementary Fig. 14: Comparison between PoolHapX and existing human haplotype reconstruction tools**. For all panels, the upper half shows the accuracy for haplotype identity (MCC) and the lower half shows the accuracy for haplotype frequency (JSD). The x-axis denotes number of genetic variants in the haplotype. Boxes extend to the first and third quartile; whiskers extend to the upper and lower value. Comparing with tools to reconstruct human haplotypes, Hippo and AEM (using 26 populations in 1000 Genome Project data). **a**. Coverage at 100X. **b**. Coverage at 250X.

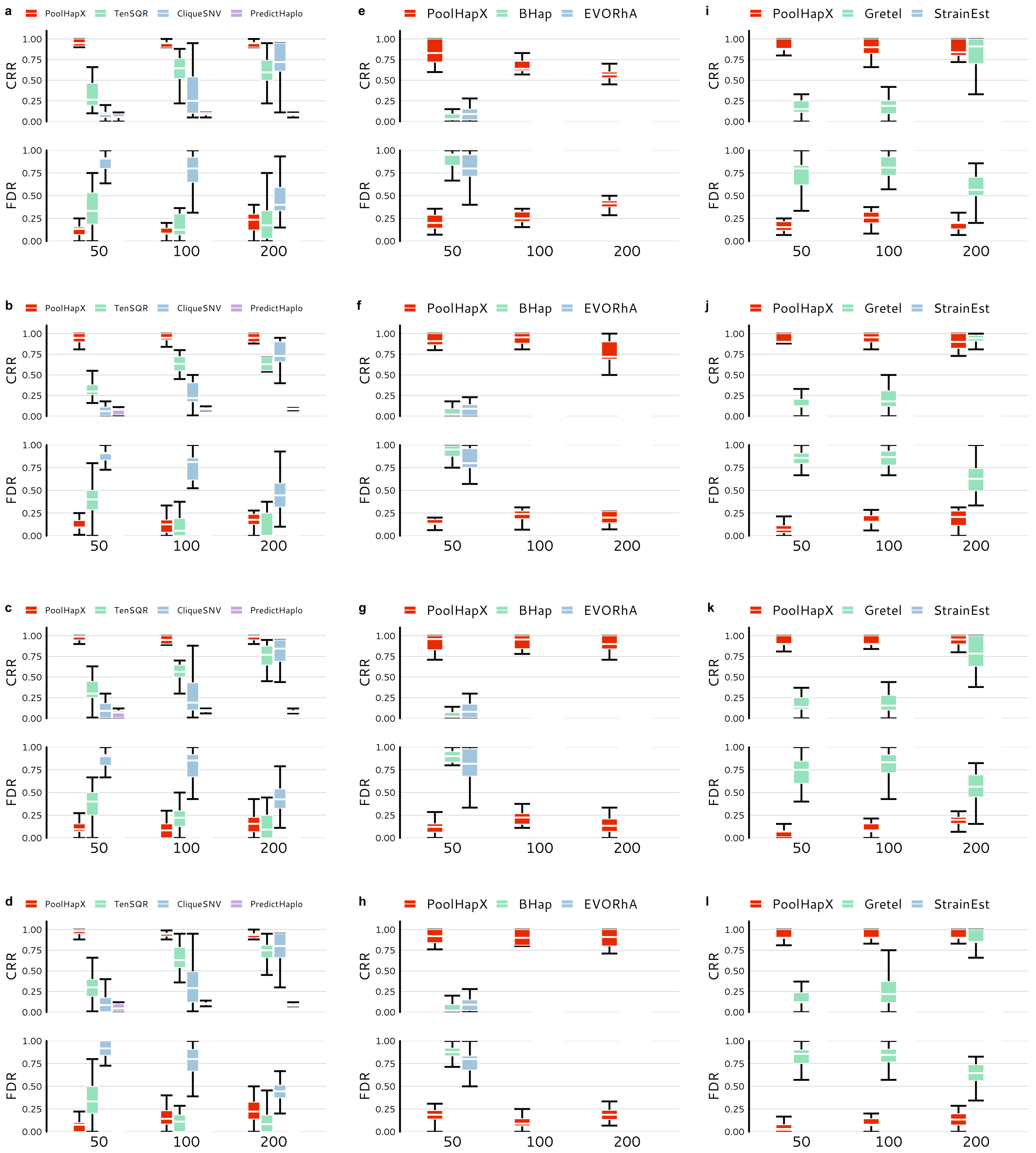


**Supplementary Figure 15: Comparison between PoolHapX and existing haplotype reconstruction tools in terms of Correctly Reconstructed Rate and False Discovery Rate**. In all panels, the upper set of box plots show the values of Correctly Reconstructed Rate (CRR) and the lower set shows the values of False Discovery Rate (FDR). The x-axis denotes number of genetic variants in the haplotype. Boxes extend to the first and third quartile, and whiskers extend to the upper and lower value. **Figures a-d** compare PoolHapX against the viral haplotype reconstruction tools TenSQR, PredictHaplo and CliqueSNV. Because PredictHaplo only reconstructed one or two haplotypes for each pool, the FDR was either 0 or 1. These results made it difficult to present with box plots, therefore, for FDR we only compared the PoolHapX with TenSQR and CliqueSNV in our figures. **a.** Sequencing coverage per pool = 2500X, number of pools = 25. Number of haplotypes for 50 loci, 100 loci and 200 loci are 32, 54 and 33, respectively. **b.** Sequencing coverage per pool = 5000X, number of pools = 25. Number of haplotypes for 50 loci, 100 loci and 200 loci are 32, 54 and 33, respectively. **c.** Sequencing coverage per pool = 2500X, number of pools = 50. Number of haplotypes for 50 loci, 100 loci and 200 loci are 41, 73 and 42, respectively. **d.** Sequencing coverage per pool = 5000X, number of pools = 50. Number of haplotypes for 50 loci, 100 loci and 200 loci are 41, 73 and 42, respectively. **Figures e-h** compare PoolHapX against the bacterial haplotype reconstruction tools Bhap and EVORhA. For Bhap and EVORhA, when the number of loci equals to 100 and 200, their MCC values are all below 0.8, leading no true positives. Since there are no true positives, their CRR values are all 0 and FDR values are all 1. **e.** Sequencing coverage per pool = 100X, number of pools = 25. Number of haplotypes for 50 loci, 100 loci and 200 loci are 36, 31 and 36, respectively. **f.** Sequencing coverage per pool = 250X, number of pools = 25. Number of haplotypes for 50 loci, 100 loci and 200 loci are 36, 31 and 36, respectively. **g.** Sequencing coverage per pool = 100X, number of pools = 50. Number of haplotypes for 50 loci, 100 loci and 200 loci are 42, 43 and 43, respectively. **h.** Sequencing coverage per pool = 250X, number of pools = 50. Number of haplotypes for 50 loci, 100 loci and 200 loci are 42, 43 and 43, respectively. **Figures i-l** compare PoolHapX against the metagenomic haplotype reconstruction tools Gretel and StrainEst. For StrainEst, it only reconstructed a reference haplotype, there is no TP values, so the MCC values cannot be calculated. Since both CRR and FDR are calculated based on the MCC values to define the true positives or false positives, the CRR and FDR values cannot be calculated either. **i**. Sequencing coverage per pool = 25X, number of pools = 25. Number of haplotypes for 50 loci, 100 loci and 200 loci are 30, 30 and 31, respectively. **j.** Sequencing coverage per pool = 50X, number of pools = 25. Number of haplotypes for 50 loci, 100 loci and 200 loci are 30, 30 and 31, respectively. **k.** Sequencing coverage per pool = 25X, number of pools = 50. Number of haplotypes for 50 loci, 100 loci and 200 loci are 39, 37 and 36, respectively. **l.** Sequencing coverage per pool = 50X, number of pools = 50. Number of haplotypes for 50 loci, 100 loci and 200 loci are 39, 37 and 36, respectively.

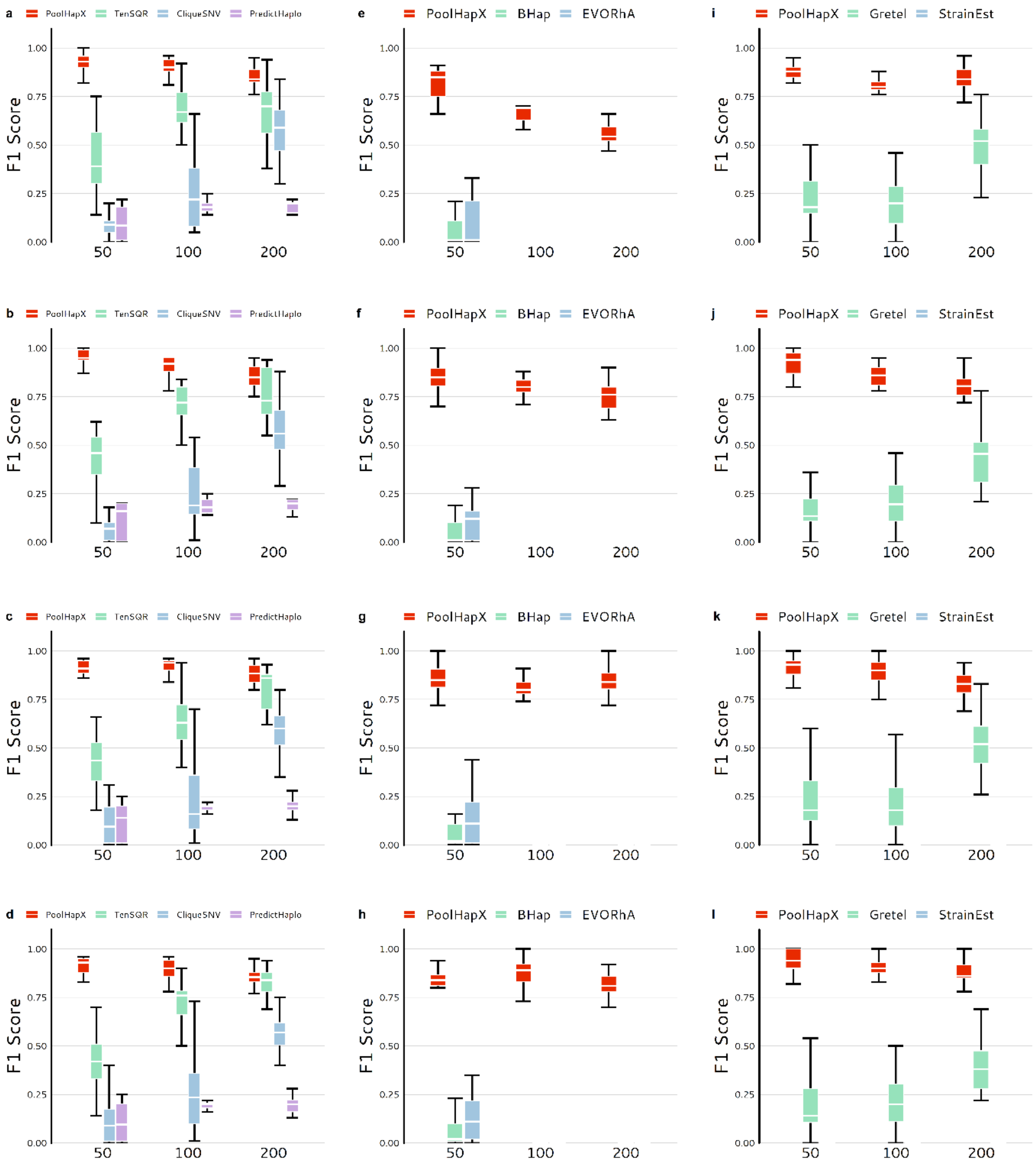


**Supplementary Figure 16: Comparison between PoolHapX and existing haplotype reconstruction tools in terms of F1-Score**. In all panels, the y-axis denotes the values of F1 score, and the x-axis denotes number of genetic variants in the haplotype. Boxes extend to the first and third quartile, and whiskers extend to the upper and lower value. **Figures a-d** compare PoolHapX against the viral haplotype reconstruction tools TenSQR, PredictHaplo and CliqueSNV. **a.** Sequencing coverage per pool = 2500X, number of pools = 25. Number of haplotypes for 50 loci, 100 loci and 200 loci are 32, 54 and 33, respectively. **b.** Sequencing coverage per pool = 5000X, number of pools = 25. Number of haplotypes for 50 loci, 100 loci and 200 loci are 32, 54 and 33, respectively. **c.** Sequencing coverage per pool = 2500X, number of pools = 50. Number of haplotypes for 50 loci, 100 loci and 200 loci are 41, 73 and 42, respectively. **d.** Sequencing coverage per pool = 5000X, number of pools = 50. Number of haplotypes for 50 loci, 100 loci and 200 loci are 41, 73 and 42, respectively. **Figures e-h** compare PoolHapX against the bacterial haplotype reconstruction tools Bhap and EVORhA. For Bhap and EVORhA, when the number of loci equals to 100 and 200, their MCC values are all below 0.8, leading no true positives. Since there are no true positives, their F1 scores are all 0. **e.** Sequencing coverage per pool = 100X, number of pools = 25. Number of haplotypes for 50 loci, 100 loci and 200 loci are 36, 31 and 36, respectively. **f.** Sequencing coverage per pool = 250X, number of pools = 25. Number of haplotypes for 50 loci, 100 loci and 200 loci are 36, 31 and 36, respectively. **g.** Sequencing coverage per pool = 100X, number of pools = 50. Number of haplotypes for 50 loci, 100 loci and 200 loci are 42, 43 and 43, respectively. **h.** Sequencing coverage per pool = 250X, number of pools = 50. Number of haplotypes for 50 loci, 100 loci and 200 loci are 42, 43 and 43, respectively. **Figures i-l** compare PoolHapX against the metagenomic haplotype reconstruction tools Gretel and StrainEst. For StrainEst, it only reconstructed a reference haplotype, there is no TP values, so the MCC values cannot be calculated. Since both CRR and FDR are calculated based on the MCC values to define the true positives or false positives, the F1 scores cannot be calculated either. **i**. Sequencing coverage per pool = 25X, number of pools = 25. Number of haplotypes for 50 loci, 100 loci and 200 loci are 30, 30 and 31, respectively. **j.** Sequencing coverage per pool = 50X, number of pools = 25. Number of haplotypes for 50 loci, 100 loci and 200 loci are 30, 30 and 31, respectively. **k.** Sequencing coverage per pool = 25X, number of pools = 50. Number of haplotypes for 50 loci, 100 loci and 200 loci are 39, 37 and 36, respectively. **l.** Sequencing coverage per pool = 50X, number of pools = 50. Number of haplotypes for 50 loci, 100 loci and 200 loci are 39, 37 and 36, respectively.

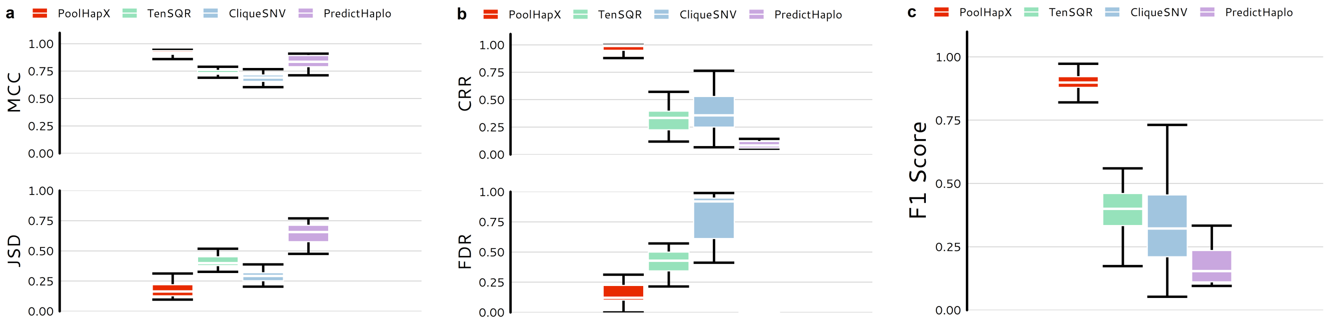


**Supplementary Figure 17: Comparison between PoolHapX and viral haplotype reconstruction tools using mocked data based on the 5-HIV strains [https://github.com/cbg-ethz/5-virus-mix].** For all panels, the x-axis denotes number of genetic variants in the haplotype. Boxes extend to the first and third quartile; whiskers extend to the upper and lower value. The sequencing coverage read length and insert size are 5000, 150, and 400, respectively. Comparing to viral reconstruction tools, TenSQR, PredictHaplo and CliqueSNV. Because PredictHaplo only reconstructed one or two haplotypes for each pool, the FDR was either 0 or 1. These results made it meaningless to represent with box plots; therefore, for FDR we only compared the PoolHapX with TenSQR and CliqueSNV in our figures. **a.** the upper half shows the accuracy for haplotype identity (MCC) and the lower half shows the accuracy for haplotype frequency (JSD). **b.** the upper set of box plots show the values of Correctly Reconstructed Rate (CRR) and the lower set shows the values of False Discovery Rate (FDR).  **c.** the y-axis denotes the values of F1 score. There are 25 haplotypes randomly distributed into 25 pools in this simulation.

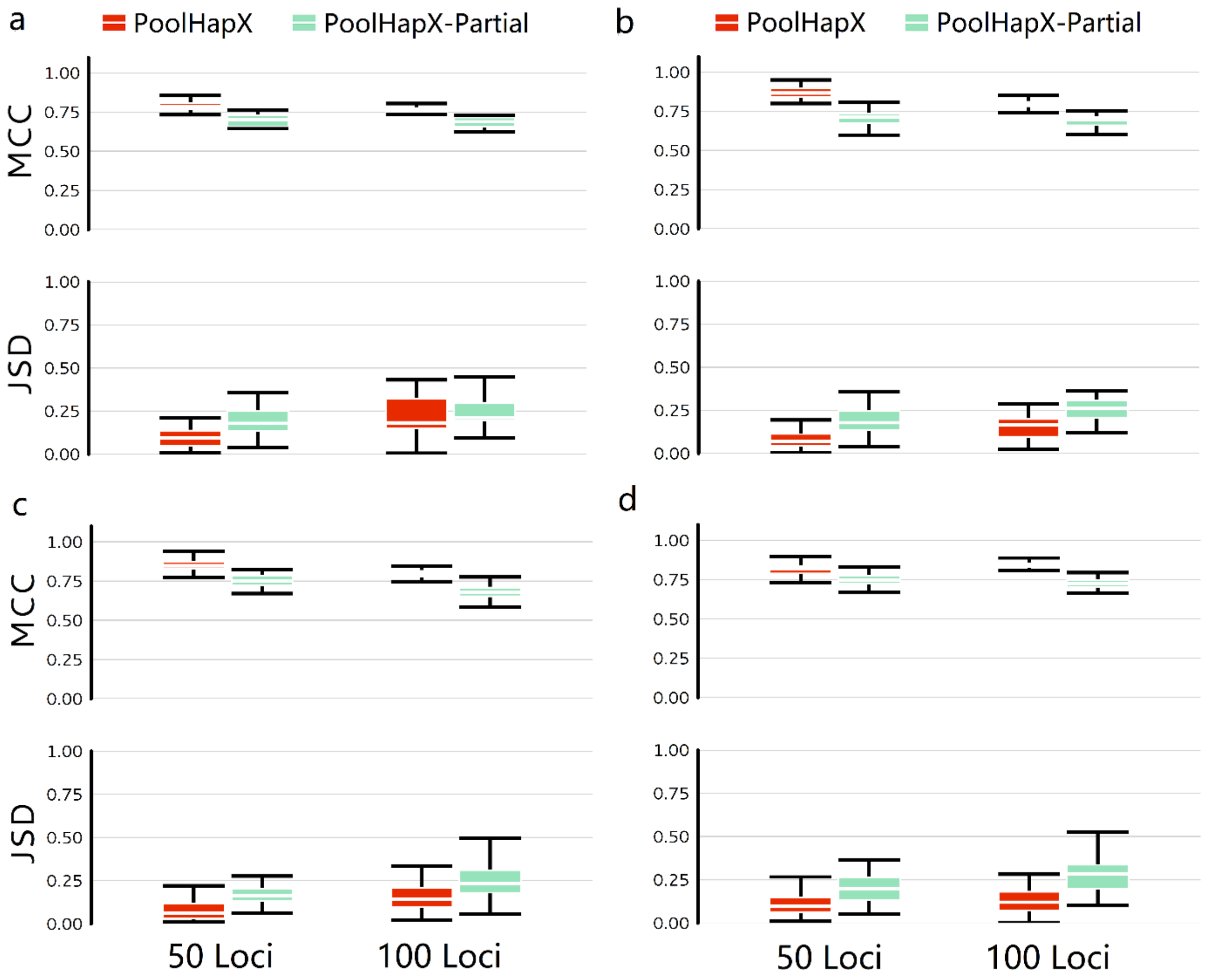


**Supplementary Figure 18: Comparison between PoolHapX and PoolHapX-Partial**. PoolHapX-Partial is the partial protocol of PoolHapX without running the regularization step. For all panels, the upper half shows the accuracy for haplotype identity (MCC) and the lower half shows the accuracy for haplotype frequency (JSD). The x-axis denotes number of genetic variants in the haplotype. Boxes extend to the first and third quartile; whiskers extend to the upper and lower value. Comparisons are done using simulated bacteria data. **a.** Sequencing coverage per pool = 100X, number of pools = 25. Number of haplotypes for 50 loci and 100 loci are 36 and 31, respectively. **b.** Sequencing coverage per pool = 250X, number of pools = 25. Number of haplotypes for 50 loci and 100 loci are 36 and 31, respectively. **c.** Sequencing coverage per pool = 100X, number of pools = 50. Number of haplotypes for 50 loci and 100 loci are 42 and 43, respectively. **d.** Sequencing coverage per pool = 250X, number of pools = 50. Number of haplotypes for 50 loci and 100 loci are 42 and 43, respectively.

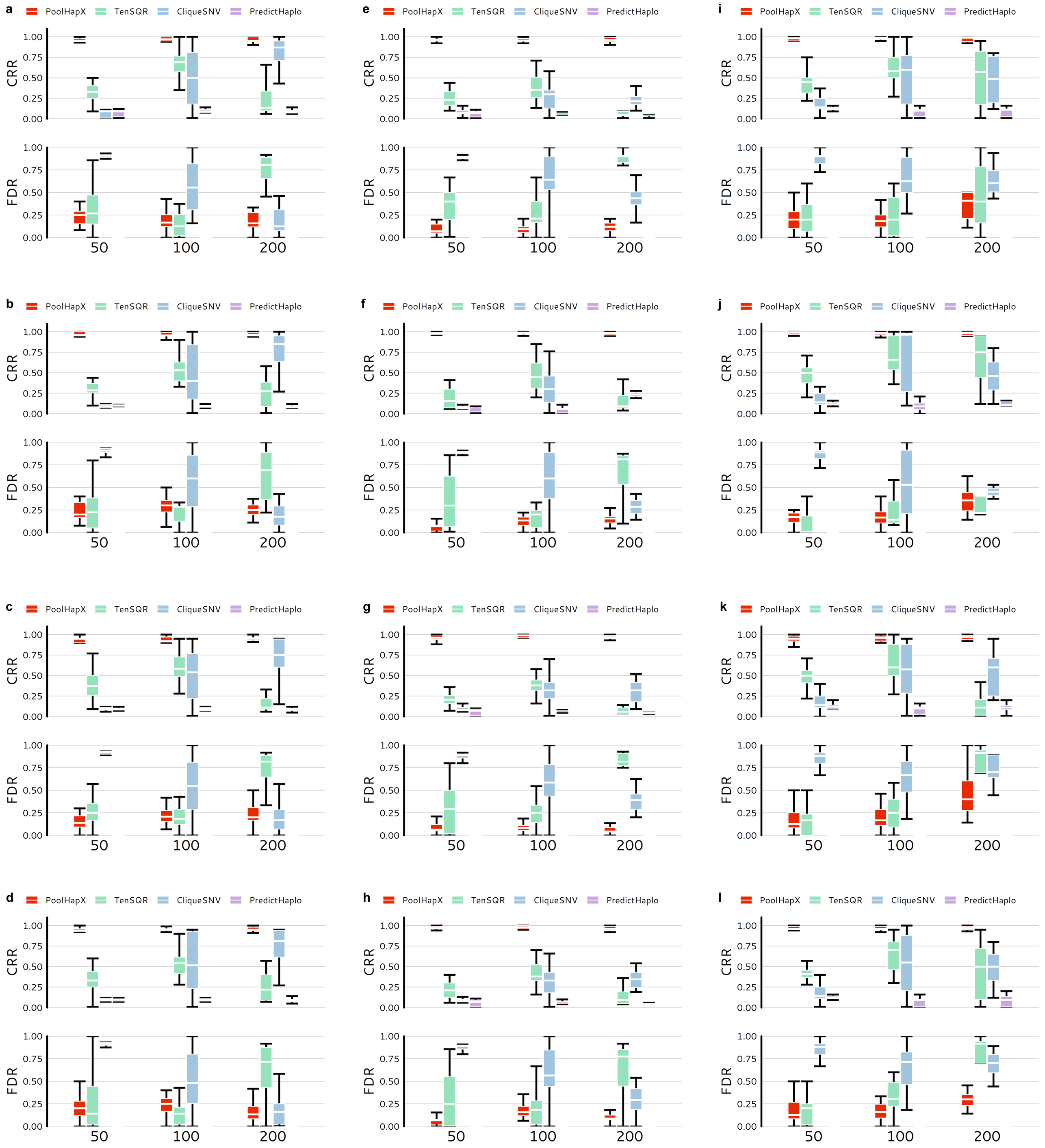


**Supplementary Figure 19: PoolHapX is robust to the within-host dynamics due to selective forces, measured by of Correctly Reconstructed Rate and False Discovery Rate**. In all panels, the upper set of box plots show the values of Correctly Reconstructed Rate (CRR) and the lower set shows the values of False Discovery Rate (FDR). The x-axis denotes number of genetic variants in the haplotype. Boxes extend to the first and third quartile, and whiskers extend to the upper and lower value. Comparing to viral reconstruction tools, TenSQR, PredictHaplo and CliqueSNV. Because PredictHaplo only reconstructed one or two haplotypes for each pool, the FDR was either 0 or 1. These results made it meaningless to represent with box plots, therefore, for FDR we only compared the PoolHapX with TenSQR and CliqueSNV in our figures. **Figures a-d** compare haplotype reconstruction tools under the scenario of negative selection. **a.** Sequencing coverage per pool = 2500X, number of pools = 25. Number of haplotypes for 50 loci, 100 loci and 200 loci are 31, 31 and 42, respectively. **b.** Sequencing coverage per pool = 5000X, number of pools = 25. Number of haplotypes for 50 loci, 100 loci and 200 loci are 31, 31 and 42, respectively. **c.** Sequencing coverage per pool = 2500X, number of pools = 50. Number of haplotypes for 50 loci, 100 loci and 200 loci are 36, 36 and 45, respectively. **d.** Sequencing coverage per pool = 5000X, number of pools = 50. Number of haplotypes for 50 loci, 100 loci and 200 loci are 36, 36 and 45, respectively. **Figures e-h** compare haplotype reconstruction tools under the scenario of positive selection. **e.** Sequencing coverage per pool = 2500X, number of pools = 25. Number of haplotypes for 50 loci, 100 loci and 200 loci are 32, 40 and 77, respectively. **f.** Sequencing coverage per pool = 5000X, number of pools = 25. Number of haplotypes for 50 loci, 100 loci and 200 loci are 32, 40 and 77, respectively. **g.** Sequencing coverage per pool = 2500X, number of pools = 50. Number of haplotypes for 50 loci, 100 loci and 200 loci are 41, 50 and 104, respectively. **h.** Sequencing coverage per pool = 5000X, number of pools = 50. Number of haplotypes for 50 loci, 100 loci and 200 loci are 41, 50 and 104, respectively. **Figures i-l** compare haplotype reconstruction tools under the scenario of selective sweep. **i.** Sequencing coverage per pool = 2500X, number of pools = 25. Number of haplotypes for 50 loci, 100 loci and 200 loci are 21, 30 and 24, respectively. **j.** Sequencing coverage per pool = 5000X, number of pools = 25. Number of haplotypes for 50 loci, 100 loci and 200 loci are 21, 30 and 24, respectively. **k.** Sequencing coverage per pool = 2500X, number of pools = 50. Number of haplotypes for 50 loci, 100 loci and 200 loci are 23, 36 and 27, respectively. **l.** Sequencing coverage per pool = 5000X, number of pools = 50. Number of haplotypes for 50 loci, 100 loci and 200 loci are 23, 36 and 27, respectively.

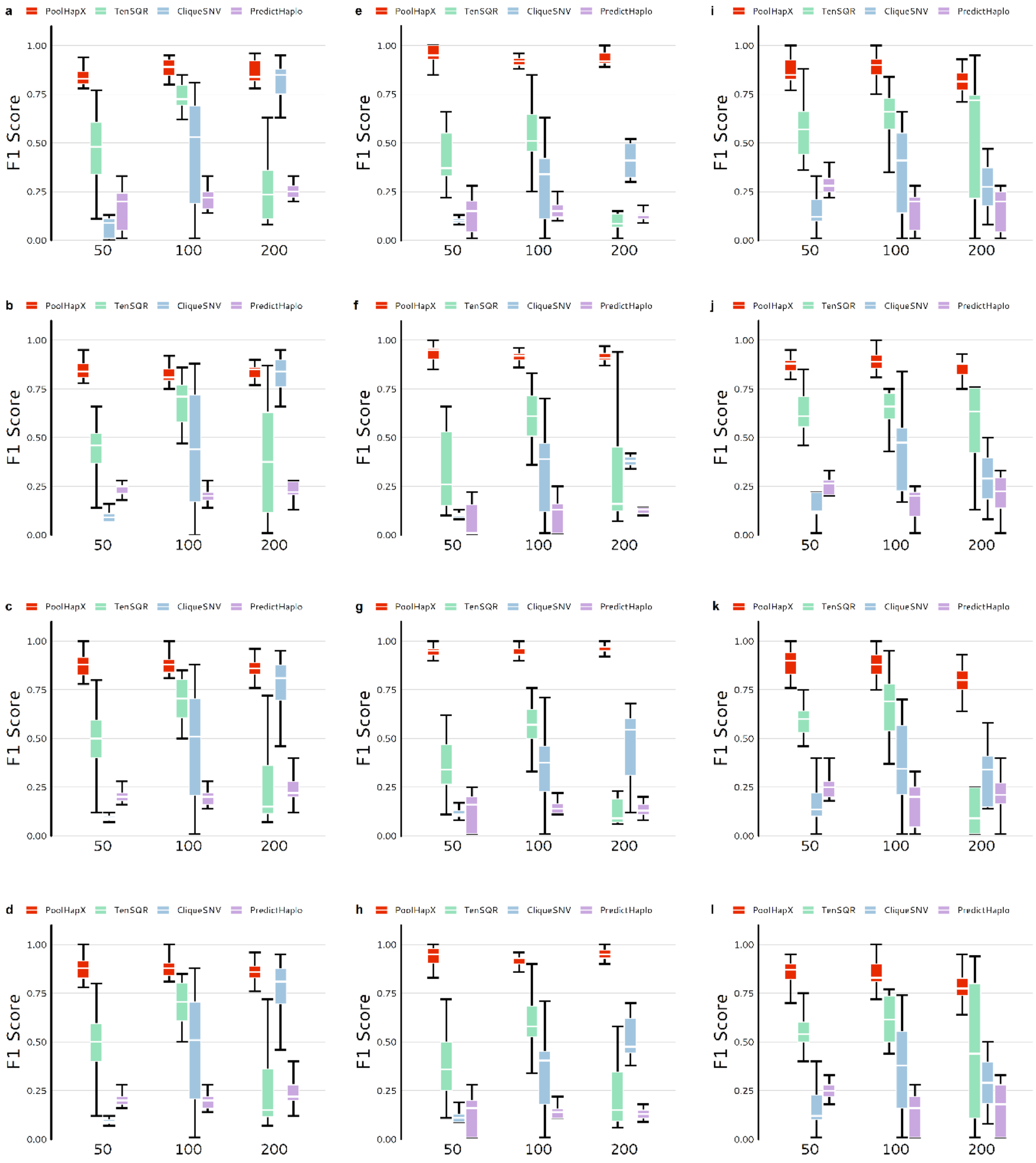


**Supplementary Figure 20: PoolHapX is robust to the within-host dynamics due to selective forces measured by F1-Score**. In all panels, the y-axis denotes the values of F1 score, and the x-axis denotes number of genetic variants in the haplotype. Boxes extend to the first and third quartile, and whiskers extend to the upper and lower value. Comparing to viral reconstruction tools, TenSQR, PredictHaplo and CliqueSNV. **Figures a-d** compare haplotype reconstruction tools under the scenario of negative selection. **a.** Sequencing coverage per pool = 2500X, number of pools = 25. Number of haplotypes for 50 loci, 100 loci and 200 loci are 31, 31 and 42, respectively. **b.** Sequencing coverage per pool = 5000X, number of pools = 25. Number of haplotypes for 50 loci, 100 loci and 200 loci are 31, 31 and 42, respectively. **c.** Sequencing coverage per pool = 2500X, number of pools = 50. Number of haplotypes for 50 loci, 100 loci and 200 loci are 36, 36 and 45, respectively. **d.** Sequencing coverage per pool = 5000X, number of pools = 50. Number of haplotypes for 50 loci, 100 loci and 200 loci are 36, 36 and 45, respectively. **Figures e-h** compare haplotype reconstruction tools under the scenario of positive selection. **e.** Sequencing coverage per pool = 2500X, number of pools = 25. Number of haplotypes for 50 loci, 100 loci and 200 loci are 32, 40 and 77, respectively. **f.** Sequencing coverage per pool = 5000X, number of pools = 25. Number of haplotypes for 50 loci, 100 loci and 200 loci are 32, 40 and 77, respectively. **g.** Sequencing coverage per pool = 2500X, number of pools = 50. Number of haplotypes for 50 loci, 100 loci and 200 loci are 41, 50 and 104, respectively. **h.** Sequencing coverage per pool = 5000X, number of pools = 50. Number of haplotypes for 50 loci, 100 loci and 200 loci are 41, 50 and 104, respectively. **Figures i-l** compare haplotype reconstruction tools under the scenario of selective sweep. **i.** Sequencing coverage per pool = 2500X, number of pools = 25. Number of haplotypes for 50 loci, 100 loci and 200 loci are 21, 30 and 24, respectively. **j.** Sequencing coverage per pool = 5000X, number of pools = 25. Number of haplotypes for 50 loci, 100 loci and 200 loci are 21, 30 and 24, respectively. **k.** Sequencing coverage per pool = 2500X, number of pools = 50. Number of haplotypes for 50 loci, 100 loci and 200 loci are 23, 36 and 27, respectively. **l.** Sequencing coverage per pool = 5000X, number of pools = 50. Number of haplotypes for 50 loci, 100 loci and 200 loci are 23, 36 and 27, respectively.

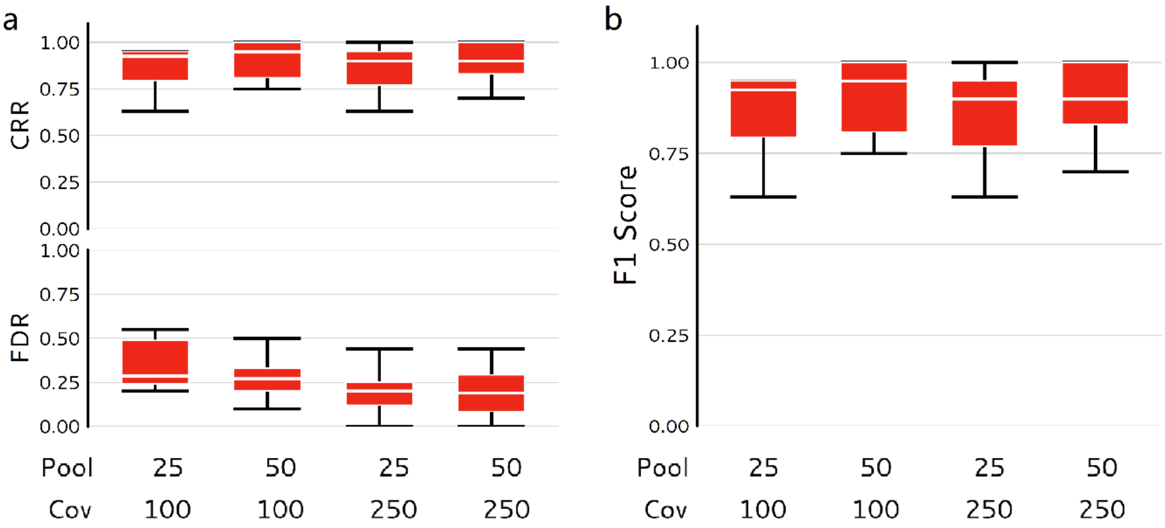


**Supplementary Figure 21: PoolHapX + single-cell linked-reads.** The PoolHapX is applying to simulated 10x linked-reads using based on combinations of different number of pools (25,50) and sequencing coverage (100, 250). The number of haplotypes is 31 for 570 loci. **a.** the upper set of box plots show the values of Correctly Reconstructed Rate (CRR) and the lower set shows the values of False Discovery Rate (FDR). **b.** the y-axis denotes the values of F1 score.

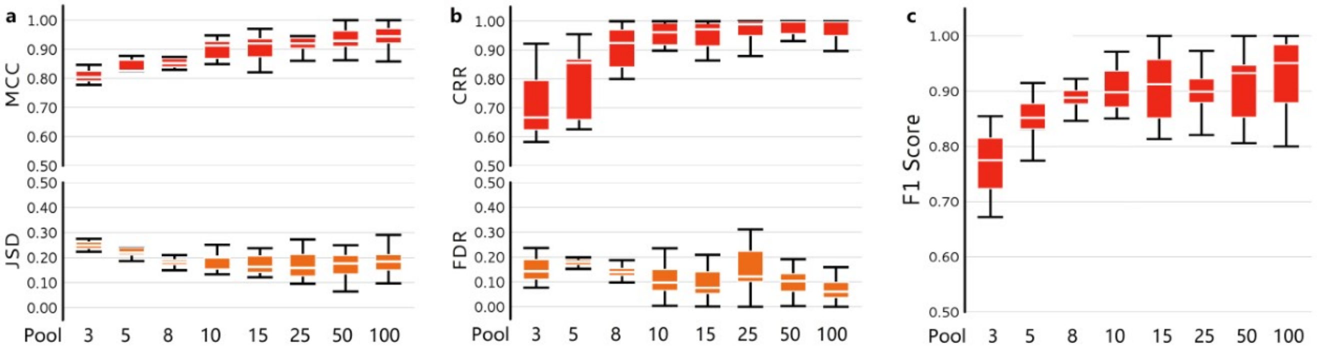


**Supplementary Figure 22: The performance of PoolHapX on different number of pools (3, 5, 8,10,15, 25, 50, 100) with sequencing coverage 5000 and 25 global haplotypes.** The 5-HIV-mix data is used as the template in simulations. **a.** the upper panel shows the accuracy for haplotype identity (MCC) and the lower panel shows the accuracy for haplotype frequency (JSD). **b.** the upper panel shows the values of Correctly Reconstructed Rate (CRR) and the lower panel shows the values of False Discovery Rate (FDR). **c.** the y-axis denotes the values of F1 score. In general, when the number of pools increased to 8, the performances of PoolHapX are relative stable with an average MCC, CRR and F1 score around 0.90, JSD around 0.20, and FDR around 0.10.

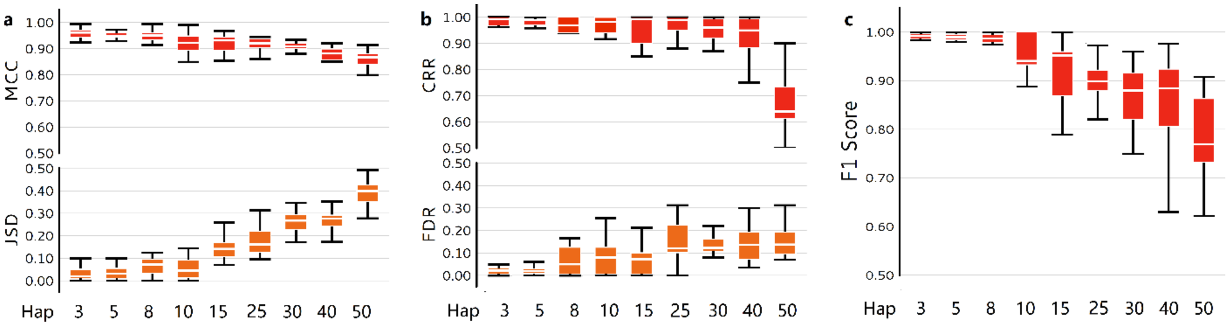


**Supplementary Figure 23: The performance of PoolHapX on different number of global haplotypes (3, 5, 8,10,15, 25, 30, 40, 50) with sequencing coverage 5000 and 25 pools**. The 5-virus-mix data is used in simulation. **a.** the upper panel shows the accuracy for haplotype identity (MCC) and the lower panel shows the accuracy for haplotype frequency (JSD). **b.** the upper panel shows the values of Correctly Reconstructed Rate (CRR) and the lower panel shows the values of False Discovery Rate (FDR). **c.** the y-axis denotes the values of F1 score. In general, PoolHapX performs better with less global haplotypes, when the number of haplotypes increased to 50, there is a bit of drop regarding to the overall performances of PoolHapX.
